# Diet-Dependent Cognitive Benefits of Exogenous Ketone Body Precursor, (R,S)-1,3,-Butanediol, in a Mouse Model of Tauopathy

**DOI:** 10.64898/2026.06.03.729999

**Authors:** Kyle Fulghum, Abdirahman Hayir, Ruusu Ankeriasniemi, Cynthia Shaddy-Gouvion, Cha Mee Vang, Sebastian F. Salathe, Eric D. Queathem, Curtis C. Hughey, Mohammed Haeri, John P. Thyfault, Patrycja Puchalska, Peter A. Crawford

## Abstract

Alzheimer’s disease and related tauopathies are escalating public health threats, particularly in the context of obesity and metabolic dysfunction, which accelerate cerebral glucose hypometabolism, tau pathology, neurodegeneration, and cognitive decline. Ketogenic therapies reconfigure systemic fuel metabolism, with emerging evidence for neuroprotection. (R,S)-1,3-butanediol (BD) raises circulating D- and L-β-hydroxybutyrate (βOHB) concentrations. To evaluate whether BD improves cognitive function across dietary contexts, male and female tau-transgenic mice and littermate controls received 10% BD in drinking water for 20 or 30 weeks starting at 6 weeks of age. BD rapidly induced ketosis (1.5-3.0 mM βOHB) in chow-fed mice, with L-βOHB contributing to ∼75% of the circulating βOHB pool. Despite minimal effects of BD on body weight and glucose homeostasis, and no effect on histopathological tau signal, 20-week BD treatment improved memory to control levels in chow-fed female tauopathy mice. Isotope-tracing untargeted metabolomics revealed that BD-treatment differentially affected glucose-derived ^13^C-enrichment of metabolites in brains of male and female mice. BD-induced cognitive benefits in tau-transgenic mice were abrogated when mice were maintained on BD for 30 weeks on standard chow or when mice were administered BD over 20 weeks while maintained on a high-fat, Western diet, Notably, BD-induced ketosis was blunted in mice consuming Western diet. Moreover, intermittent ketogenic diet-induced ketosis failed to improve cognition in Western diet-fed tauopathy mice. These results suggest BD-induced ketosis extends cognitive benefits in a manner dependent on biological sex and nutritional metabolic status. Taken together, these data contextualize the roles of βOHB as modulators of cognitive resilience in tauopathy.

## INTRODUCTION

Neurodegenerative diseases, such as Alzheimer’s disease (AD) and related tauopathies, present a growing public health crisis in the United States^1^. Important hallmarks of these diseases include progressive reduction in cerebral glucose metabolism and the accumulation of neurofibrillary tangles^2^, which together contribute to neuronal loss and cognitive decline^3,4^. Over the next 25 years, the occurrence of neurodegenerative diseases that lead to cognitive impairment is estimated to double^1^, with limited therapeutic options currently available totreat or prevent cognitive decline^5^. Therefore, decoding the mechanistic interplay between brain glucose metabolism, neurodegenerative pathology, and cognitive function remains central to developing effective interventions.

Aging is the greatest risk factor for cognitive decline^6^, but recent studies also support a strong link between mid-life obesity and the development of cognitive impairment^7^. In fact, neurodegenerative diseases share important pathological features with cardiometabolic diseases, such as insulin resistance^8^ and reductions in brain glucose metabolism^9,10^. Importantly, ketogenic therapies have been shown to reduce glucose utilization in the brain^11^, suggesting its use as a key potential strategy in mitigating the reductions in brain glucose metabolism associated with both obesity and neurodegenerative diseases.

Ketogenic therapies elevate circulating ketone bodies and are increasingly recognized as a promising candidate for neurodegenerative diseases treatment^12–14^. Ketone bodies could serve as an alternative energy substrate, bypassing neuronal reliance on glucose and helping to maintain neurological energy homeostasis. Furthermore, ketone bodies act as signaling molecules and redox regulators that extend their neuroprotective relevance well beyond their canonical role as oxidative fuels^15^. Ketogenic interventions are a promising therapeutic option for the treatment of neurodegenerative diseases by increasing both energy availability and signaling potential. One ketogenic therapy is (R,S)-1,3-butanediol (BD), a diol precursor that is hepatically converted to both D- and L- enantiomers of the ketone body β-hydroxybutyrate (βOHB), raising circulating concentration of each enantiomer^16,17^. While D-βOHB is canonically oxidized or involved in signaling, less is known about the contributions of L-βOHB to human health. In individuals with heightened risk for or with pre-existing neurodegeneration, BD or related ketone precursors may offer neuroprotection by modulating the metabolic, pathological, and cognitive features of neurodegenerative diseases.

To date, the therapeutic effects of chronic BD administration on cognitive function and brain metabolism have not been well-characterized. Specifically, the metabolic effects of exogenously supplied ketone bodies in the setting of tauopathy-related neurodegeneration remain elusive. In this study, we investigated the effects of chronic BD treatment on cognitive function and brain metabolism in tauopathy transgenic mice consuming either a normal chow diet or a high-fat, high-sucrose Western diet to induce cardiometabolic dysfunction. Cognitive improvements were evident in BD-treated female mice on normal chow with tauopathy, and BD treatment rewired cerebral glucose metabolism. However, BD treatment failed to elevate circulating ketones or improve cognitive outcomes in mice consuming the Western diet, indicating that diet-induced metabolic dysfunction can diminish the benefits of this specific ketogenic intervention in the brain.

## 2. Methods

### 2.1 Experimental animals

All procedures were approved by the Institutional Animal Care and Use Committee at the University of Minnesota. We utilized bi-transgenic rTg4510 mice (Jackson Labs; Stock #024854) with tetracycline-inhibited transactivator protein under hemizygous control of forebrain-specific calcium-calmodulin-dependent kinase II (Camk2a), and hemizygous tetracycline-responsive element prion protein promoter sequence for expression of P301L mutant of human microtubule-associated protein tau (MAPT*P301L). As such, these mice accumulate hyperphosphorylated human tau protein in the forebrain region and develop brain pathology reproducibly by 22 weeks of age^18^. We chose to use the tauopathy mouse model since neurofibrillary tangles (NFT) better predict cognitive dysfunction in AD patients than amyloid-β alone^19^. We maintained mice on a mixed background of FVBx129S6xC57BL/6 with male breeders hemizygous for Camk2a-tTA but noncarriers for tetO-MAPT*P301L, while female breeders were hemizygous for tetO-MAPT*P301L but noncarriers for Camk2a-tTA. Bi-transgenic mice (containing hemizygous Camk2a-tTA and hemizygous tetO-MAPT*P301L; tauopathy mice), single hemizygous transgenic littermate controls, and non-transgenic littermate controls were kept on 12h:12h light:dark cycle with both diet (in absence of doxycycline) and water provided *ad libitum.* At 6 weeks of age, mice were either maintained on a normal chow diet in the absence of doxycycline, were switched to a high-fat (42% kCal) sucrose-enriched Western diet (Teklad, TD.88137), or were switched to a weekly cycling of the Western diet with a zero-carbohydrate ketogenic diet. Upon completion of each experiment, mice were anesthetized with sodium pentobarbital (150 mg/kg, i.p.) and euthanized via decapitation and excision of the heart. Tissue masses were measured via gravimetry. These procedures are consistent with the AVMA *Guidelines on Euthanasia*. Transgenes were confirmed using genotyping, with the PCR primers for Camk2a and MAPT found in **Suppl Table 1**.

### 2.2 Administration of exogenous ketone body precursor

For exogenous ketone body administration, mice were given 10% (v/v) (R,S)-1,3-butanediol (Fisher Sci #AC107620025) dissolved in drinking water beginning at 6 weeks age and continued for up to 30 weeks. Weekly body weights were recorded, and monthly blood collections were acquired via mouse tail snip. All recurring measurements and samples were taken at the same time of day (ZT1-ZT3).

### 2.3 Circulating substrate measurements

Murine blood was collected via tail snip, and a drop of blood was used to measure circulating blood glucose using the Metene TD.4116 blood glucose monitoring system (Longhua District, Shenzhen City, China). About 10 µL tail blood was collected and serum was separated via centrifugation for analyses. For measurement of total ketone bodies (acetoacetate, β-hydroxybutyrate), internal standards [U-^13^C_4_]-AcAc and [3,4,4,4-D_4_]-βOHB were spiked into cold (1:1) acetonitrile (ACN):Methanol and metabolites were extracted from diluted serum samples. Following, extracted samples were separated via reverse-phase UHPLC and detected using Thermo Fisher Scientific QExactive Plus hybrid quadrupole-orbitrap mass spectrometer via parallel reaction monitoring, as described previously^20,21^. For enantioseparation and quantification of D- and L-βOHB, extracted serum samples were dried in a speed-vac and derivatization was adapted from previous reference^20,21^. Standards of D- and L-βOHB were dried, derivatized, and injected alongside samples to confirm enantiomeric identity using an LC-MS/MS system in positive ionization mode. Additional details are highlighted in the Supplemental Methods.

### 2.4 Glucose and ketone tolerance tests

For glucose tolerance testing (GTT), mice were fasted for 18h prior to intraperitoneal administration of unlabeled glucose (1 mg/g BW for males; 2 mg/g BW for females). The BD-administered group was maintained on BD-water during GTT. For ketone tolerance testing (KTT), random-fed mice were administered unlabeled sodium D-βOHB via intraperitoneal injection (10 µmol/g BW), and blood was collected for total ketone body measurements. The 10% BD treatment was replaced with normal drinking water 48 hours prior to KTT to wash out exogenous ketone bodies in circulation.

### 2.5 Body composition

Body fat distribution was measured using EchoMRI-100 Body Composition Analyzer (EchoMRI LLC, Houston, TX) in 36-week-old tauopathy-transgenic and non-transgenic littermate control mice following 30 weeks of consuming normal drinking water or 10% BD dissolved in drinking water.

### 2.6 Immunoblot

Frozen brain samples were homogenized in 10x volume per mass protein lysis buffer that contained 20 mM Tris-HCl, 150 mM NaCl, 1 mM EDTA, 1% Triton-X 100 at pH=7.5. The buffer was supplemented with protease inhibitor cocktail (Roche, Basel, Switzerland) and phosphatase inhibitor cocktail (Sigma, St. Louis, MO). Ten micrograms of total protein were dissolved in 2% lithium dodecyl sulfide sample loading buffer and prepared for separation via SDS-polyacrylamide gel electrophoresis. Separated proteins were transferred to 0.2 µm polyvinylidene fluoride membranes. Protein targets of interest were probed in the combined soluble and insoluble fraction using 1:5000 anti-phospho-tau (Ser202, Thr205) AT8 antibody (Invitrogen #MN1020) in 5% bovine serum albumin and 1:5000 anti-tau (Tau-5, total tau) antibody (Invitrogen #ahb0042) in 5% nonfat, dry milk. 1:10,000 anti-mouse secondary antibody was used for phospho-tau and 1:10,000 anti-mouse was used for total tau, both dissolved in 5% nonfat, dry milk. Images were acquired using the ChemiDoc MP Imaging System (BioRad, Hercules, CA).

### 2.7 Barnes maze testing

Cognitive function was assessed using a Barnes Maze protocol as previously described^22,23^ and in the Supplementary Methods. We used a maze with a tan background since rTg4510 mice present with white, agouti, or black fur. We recorded primary errors, primary latency to escape hole entry, and percentage of time spent in the goal quadrant derived from video acquisition using AnyMaze Software (Stoelting Co.; Wood Dale, IL) provided by the UMN Behavioral Phenotyping Core.

### 2.8 Brain immunohistochemistry

Murine brains were collected following 30-week treatment of 10% BD and were immersion-fixed in 10% formalin for 24 hours, followed by tissue blocking and processing in increasing concentrations of ethanol (50%-100%), xylene, and paraffin before embedding. Immunohistochemistry was performed similarly to previously described^24,25^. Formalin-fixed, paraffin-embedded mouse brain sections were cut and mounted on glass microscope slides, deparaffinized by standard methods, then treated with heat-induced epitope retrieval using 10 mM sodium citrate buffer, pH-6.0. Immunostaining for both mouse monoclonal antibodies, anti-Tau AT8 pS202/pT205 (1:500 dilution, Fisher Scientific, Walthan, MA) and anti-NeuN (1:500 dilution, Millipore, Burlington, MA), was done using the M.O.M.® (Mouse on Mouse) ImmPRESS® HRP Polymer Kit (Vector Laboratories, Burlingame, CA), while ImmPRESS Anti-Rabbit Ig (peroxidase) Polymer Detection Kit (Vector Laboratories, Burlingame, CA) was used for rabbit polyclonal anti-GFAP antibody (1:10,000 dilution in PBS, Protein Tech Group, Rosemont, IL). Per manufacturer’s instructions, sections were blocked, incubated with primary antibodies overnight at 4°C, then washed and incubated with respective polymer secondary antibodies. Staining was visualized by precipitation of diaminobenzidine using Dako Envision FLEX DAB chromagen plus substrate buffer (Agilent Technologies, Santa Clara, CA). Primary antibodies were omitted in negative controls. Sections were dehydrated and slides covered using Cytoseal-60 mounting media (Epredia, Kalamazoo, MI). Samples were compared to water control brains.

### 2.9 Isotope Tracing Untargeted Metabolomics (ITUM)

Mice were fasted 18h prior to isotope tracing experiments. Fifteen minutes after an intraperitoneal injection of 5 µmol/g BW [U-^13^C_6_]glucose, mice were euthanized with pentobarbital and decapitated for rapid collection of the brain frontal cortex, which was immediately freeze-clamped. Blood samples were collected at t=0 min and t=15 min to quantify plasma glucose concentration. Briefly, 20 mg brain samples were homogenized in (2:2:1) Methanol:ACN:H_2_O extraction buffer, and then metabolites extracted from 2 mg tissue homogenate in 1 mL extraction buffer by three cycles of vortexing, freeze-thawing, and water bath sonication. Samples were incubated at -20 C° for 1 hour, followed by 10 min centrifugation. Exactly 800 µL supernatant was transferred to new tubes, and the solvent was evaporated in a speed-vac. Finally, the dried metabolite pellet was reconstituted in 100 µL (1:1) ACN:H_2_O with the aid of vortexing and sonication, then incubated at 4 °C for 1 h before spinning samples and loading in vials for LC/MS analysis. ITUM workflow, metabolite identification, and analyses proceeded as previously described^26^ and as detailed in the Supplemental Methods. To confirm fractional enrichment of metabolites with low enrichment (<1%), manual integration using QuanBrowser on Thermo Fisher Xcalibur software was performed.

### 2.10 Shotgun lipidomics

Lipidomics extractions and analyses were performed as previously described^27^, and in the Supplemental Methods, using ∼20 mg frozen brain tissue. Bulk lipid extraction was completed using a modified Bligh-Dyer method^28^, before multidimensional MS-based shotgun lipidomics.

### 2.11 Statistical analyses

Statistical significance was determined where p<0.05, and statistical values were plotted when p<0.10. Analyses were performed using Student’s t-test or 2-way ANOVA with Tukey’s post-hoc test. For repeated measures, a 2-way ANOVA with mixed effects analysis was performed to identify significant differences between timepoints.

For correlation testing, we compared Barnes Maze area under the curve (AUC) against several variables measured throughout the study and computed Pearson’s r correlation using GraphPad Prism. Significant (p<0.05) and trending variables, along with correlation directionality, were used to construct a Metabolic Health Score (MHS) for each animal that could model Barnes Maze AUC and plotted the linear regression with 95% confidence interval. Individual measures were standardized using Z-score normalization to account for differences in scale and variance, then the MHS was calculated as the unweighted sum of standardized variables.

For lipidomics, we analyzed total concentrations of individual lipid species and grouped them by lipid classes, using R for analyses. Two-way ANOVA models (genotype × treatment) were fitted for concentration values of each lipid with post-hoc comparisons performed using estimated marginal means with pairwise contrasts, and p-values adjusted using the Benjamini–Hochberg false discovery rate method. Intragroup coefficient of variation and class-level lipid abundance plots were constructed using R. For partial least-squares discriminant analysis (PLS-DA) and variable importance projection (VIP) plots, we uploaded absolute concentrations of lipid species for analysis using Metaboanalyst 6.0 software (http://www.metaboanalyst.ca/)^29^. Metabolites with missing data were omitted and data were filtered by interquartile range, followed by log_10_ transformation to fit a normality curve and to buffer against skewed concentrations of certain lipid species before PLS-DA and VIP plots were made. For multiple comparison testing, q-values were calculated using a method embedded within Metaboanalyst that controls false discovery rate (FDR), where we used an FDR of p<0.10 to assume significance. Individual lipid species were plotted as absolute concentration with p-values indicated when p<0.05 following two-way ANOVA and Tukey’s post-hoc test. All other graphs were made using GraphPad Prism (GraphPad, Boston, MA), with statistical analyses performed in the software.

## 3. Results

### 3.1 (R,S)-1,3-Butanediol Promotes Physiological Ketosis via Sustained Elevation in L-βOHB

To optimize BD delivery in male and female mice, we first tested the acute effects of administering BD dissolved in drinking water to non-transgenic, littermate control mice (**Fig 1A**). To characterize the kinetics of acute 10% BD administration, previously BD-naïve male and female mice were given access to 10% BD in drinking water for 10 min, after which BD was removed. Serum total ketone bodies were measured over time. Thirty minutes after exposure, circulating βOHB concentrations were markedly elevated (∼30-fold increase in females and ∼4-fold increase in males), but rapidly decreased at the 60 min timepoint and returned to baseline at 180 min (**Fig 1B**). The percentage of L-βOHB enantiomer in serum was increased in BD-treated mice, but with delayed clearance relative to overall clearance of βOHB in the mice (**Fig 1B**).

**Figure 1.**
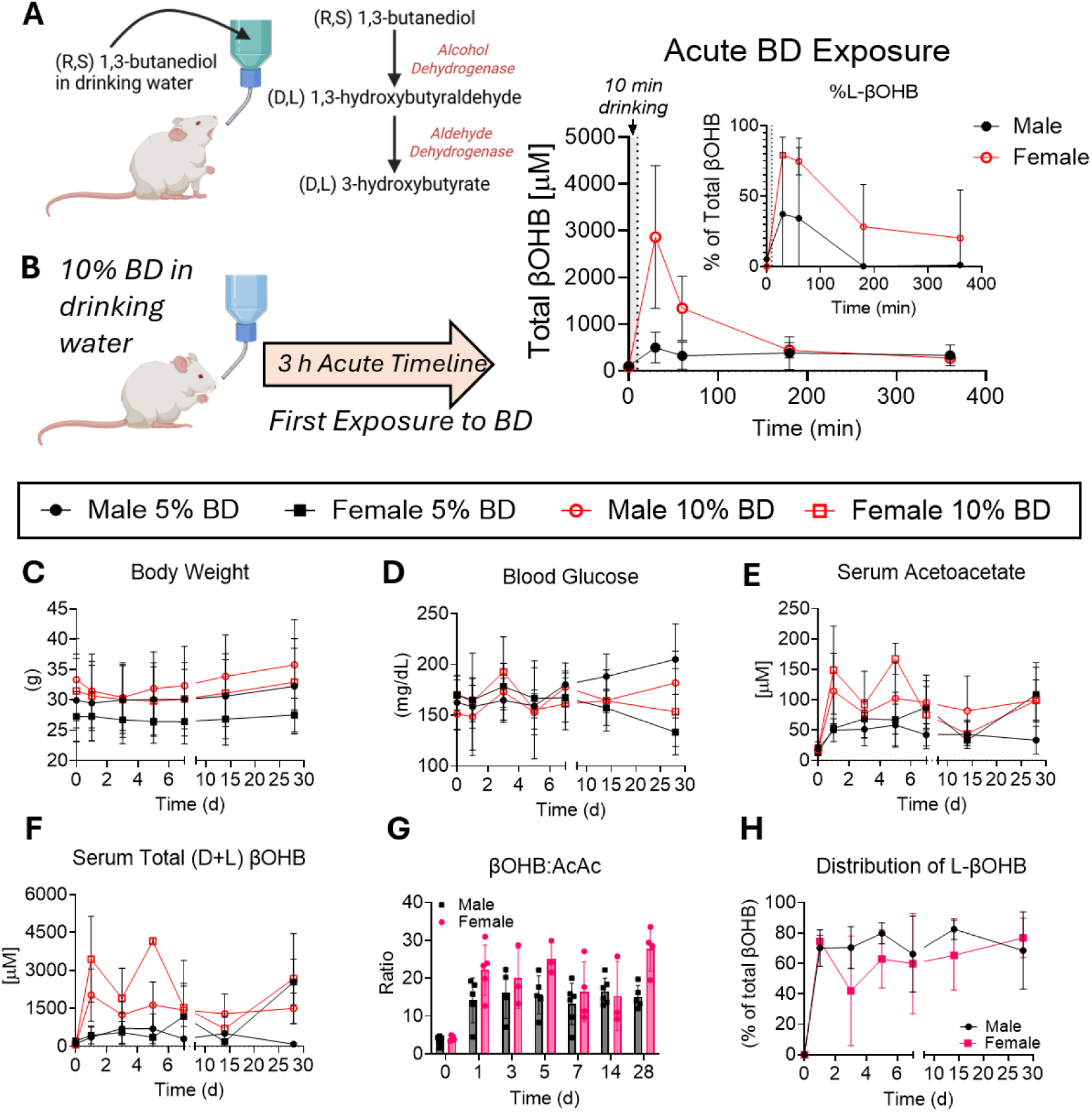
(R,S)-1,3-butanediol (BD) in drinking water promotes euglycemic, physiological ketosis. (**A**) Male and female mice were given either 5% or 10% BD in drinking water for 28 days. Schematic of 1,3-butanediol metabolism in mice. (**B**) BD-naïve mice were given a 10 min exposure to 10% BD in the drinking water, and the elevation of serum βOHB was measured, along with the distribution of D- and L-βOHB. An additional cohort of mice was treated with 5% BD or 10% BD in the drinking water for 28 days, with male and female body weight (**C**), blood glucose (**D**), serum acetoacetate (**E**), and total serum βOHB (**F**) measured throughout the 28-day administration. (**G**) 10% BD consumption increased the ratio of blood βOHB:AcAc and circulating L-βOHB distribution (**H**) during 28-day administration. Data are expressed as mean ± SD, *n*=3-8/group. Statistical tests included two-way ANOVA with Tukey’s post-hoc test. BD = 1,3-butanediol, AcAc = acetoacetate; βOHB= β-hydroxybutyrate.

To measure the effects of sustained ketosis and the distribution of D- and L-βOHB enantiomers, we treated male and female mice for 28 days with 5% BD or 10% BD. Compared to water-treated controls, BD-treated male and female mice showed normal body weights and blood glucose over a 28-day treatment period (**Fig 1C-D** and **Suppl Fig 1**). BD treatment caused a non-statistically significant reduction in body weight ∼3 days following administration of 10% BD (<10% change in both males and females), returning to baseline values after day 5 (**Fig 1C**). In mice administered 5% BD, circulating AcAc increased modestly above baseline values with an overall, 28-d average concentration of 46.4 ± 8.7 µM in male mice and 69.7 ± 26.3 µM in female mice. The 10% BD increased AcAc to an overall average of 99.9 ± 32.6 µM in males and 103.7 ± 47.4 µM in females over the 28-d treatment (**Fig 1E**). Total βOHB (D+L enantiomers) was 442.2 ± 243.0 µM in males and 875.3 ± 886.3 µM in females mice given 5% BD, and 1.5 ± 0.3 mM or 2.4 ± 1.3 mM in male and female mice, respectively, receiving 10% BD over 28 days (**Fig 1F**). Because total serum βOHB concentrations in 5% BD-treated mice were minimally elevated compared to water-control mice (**Fig 1E-F, Suppl Fig 1C, F**), all further experiments were performed using 10% BD. Metabolism of the exogenous ketone body precursor, BD, yielded a sustained increase in the ratio of total (D+L) βOHB:AcAc in circulation, with males averaging a ratio of 15.1 ± 4.7 while females averaged a ratio of 21.2 ± 7.6 (**Fig 1G**). Importantly, since BD metabolism produces both D-βOHB and L-βOHB, we quantified the enantiomeric percentage of both βOHB enantiomers in circulation at all times while mice consumed 10% BD, with males averaging 73.0 ± 16.7% distribution of L-βOHB while females averaged 63.6 ± 24.7% distribution of L-βOHB throughout the 28-day BD administration (**Fig 1H**). L-βOHB was undetectable in the circulation of water control mice.

To determine if chronic BD-induced ketosis provokes a change in ketone body disposal, we performed a ketone tolerance test (KTT) in littermate control mice via intraperitoneal injection of 10 µmol/g BW sodium D-βOHB dissolved in water (**Fig 2A**). Prior to KTT, male and female mice were maintained on either normal drinking water or 10% BD for 20 weeks. First, we measured the clearance of βOHB in circulation derived from the 20-week BD treatment by removing the BD for 48 hours and measuring ketone bodies in serum over time. Most of the βOHB was cleared by 180 min (3.57-fold reduction in females and 3.98-fold reduction in males, **Fig 2B**). The L-βOHB enantiomer, which is typically low under physiological conditions in serum, was cleared by 540 min in both male and female mice (**Fig 2B**). During the KTT, concentrations of βOHB were not significantly different in BD-treated mice compared to water control-treated mice at any time point (**Fig 2C**); however, area under the curve (AUC) for βOHB tolerance was lower in BD-treated males compared to normal water controls, indicating more rapid clearance of ketone bodies in the BD-treated mice (**Fig 2D**; *p=0.0256*). BD treatment did not influence the clearance of βOHB in females, but ketone clearance for female water controls was more rapid than that for males (*p=0.0522*) (**Fig 2D**). Since D-βOHB interconverts with AcAc through enzymatic β-hydroxybutyrate dehydrogenase 1 (BDH1), we also measured circulating concentrations of AcAc and observed lower AUCs during the KTT than normal water-treated males (**Fig 2E-F**; *p=0.0251*). There were no significant effects measured in BD-treated females compared to normal water treatment. No differences in total (D+L) βOHB:AcAc ratio between normal water and BD treatment were observed (**Fig 2G**). One week following the KTT, mice were sacrificed and tissue collected. Following the 20-week BD treatment, there were no long-term effects of BD on body weight (**Suppl Fig 2A**), blood glucose in *ad libitum* fed mice (**Suppl Fig 2B**), or circulating total βOHB at t=0 when starting the KTT (**Suppl Fig 2C**) in either male or female mice. BD treatment reduced brain mass of control females when normalized to body weight (**Suppl Fig 2D**); however, there were no effects of BD treatment on raw or tibia length-normalized brain mass or heart mass in male or female mice (**Suppl Fig 2E-I**).

**Figure 2.**
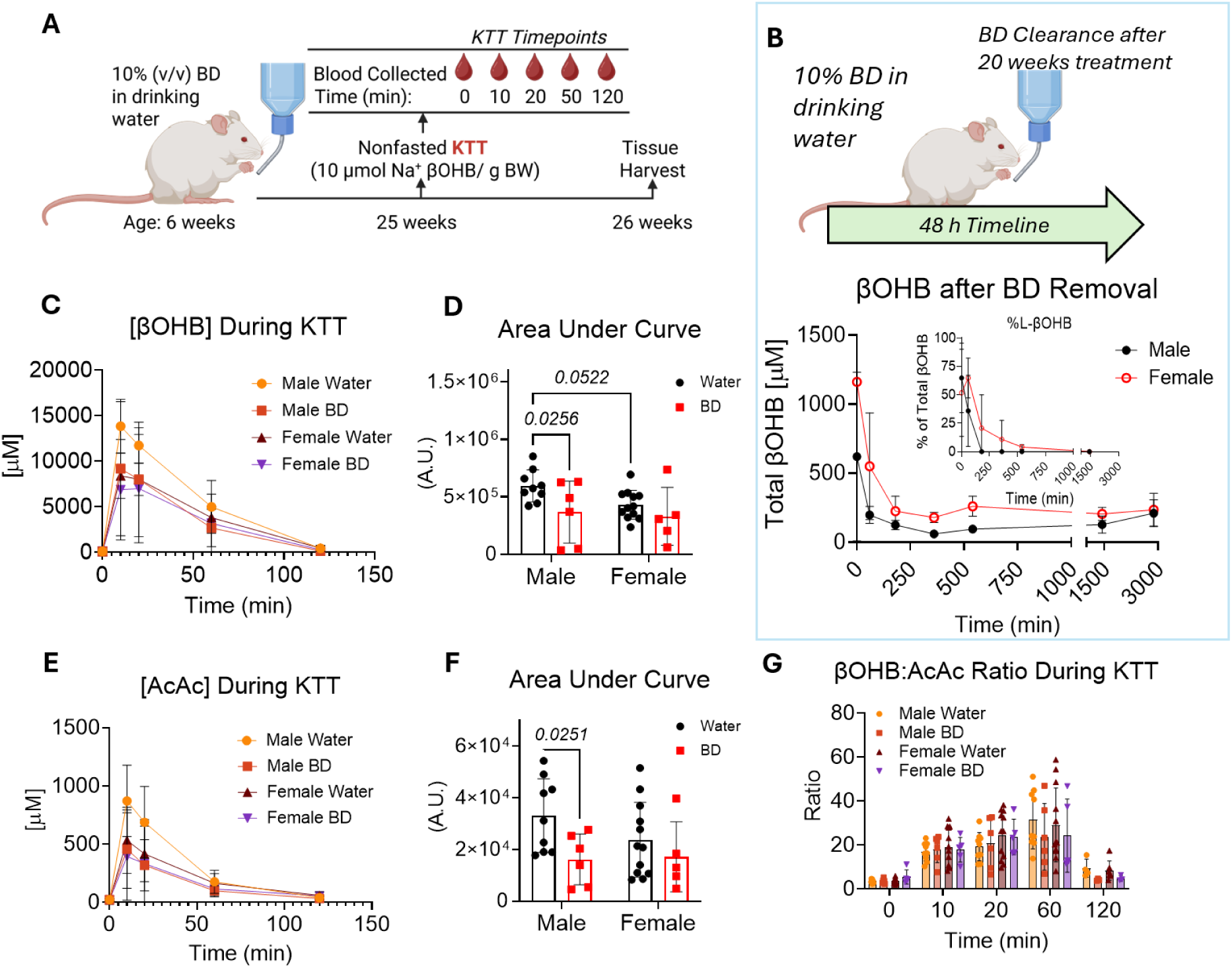
Adaptation to 20 weeks of physiological ketosis in male and female mice. (**A**) Schematic of 20 weeks 10% BD administration to assess chronic adaptation to sustained ketosis. (**B**) In one cohort of male and female mice, BD was removed after 20 weeks of treatment, and the concentration of total serum βOHB, as well as the distribution of D- and L-βOHB were measured to show ketone clearance kinetics. (**C-F**) A ketone tolerance test (KTT) was performed on a separate group of non-fasted mice via intraperitoneal injection of 10 µmol/g BW Na^+^ D-βOHB. BD was removed 48 h prior to the KTT. Serum levels of βOHB (**C-D**) and AcAc (**E-F**) with corresponding area under the curve during the 120 min KTT. (**G**) Ratio of circulating βOHB:AcAc shown during each time point of the KTT. For KTT tests, data are expressed as mean ± SD, *n*=5-11/group. P-values indicated where p<0.10 following two-way ANOVA and Tukey’s post-hoc test. BD = 1,3-butanediol, KTT = ketone tolerance test, A.U. = arbitrary units, AcAc = acetoacetate; βOHB= β-hydroxybutyrate.

### 3.2 Twenty Week BD Treatment Restores Cognitive Ability in Tauopathy Mice

After characterizing of the effects of 20-week BD administration, we sought to determine how BD influenced pathophysiology of the tauopathy mouse model, which develops neurofibrillary tangles in forebrain neurons and reproducibly displays measurable cognitive decline and brain pathology by 22 weeks of age^18^ (**Suppl Fig 3A**). Therefore, we administered 10% BD in drinking water to male and female bi-transgenic (tauopathy) mice, and respective littermate (either single transgenic or non-transgenic) controls without tauopathy, beginning at 6 weeks of age and continuing for 20 weeks (**Fig 3A**). At study conclusion of 26 weeks of age, male and female tauopathy mice showed marked elevation in both phosho-Tau (AT8) and total Tau expression in forebrain samples compared to non-tauopathy mice (**Suppl Fig 3B**). Using a Barnes Maze, we tested spatial memory and recall, noting that female tauopathy mice showed 1.88±0.35-fold increase in primary latency time to finding an escape hole during memory acquisition (**Fig 3B-C**; *p=0.0031*). Male tauopathy primary latency time was not statistically different compared to non-tauopathy littermate controls (**Suppl Fig 3C-D**). Strikingly, female tauopathy mice receiving BD treatment for 20 weeks showed improvement in primary latency (0.57 ± 0.36-fold, *p=0.0177*) and probe recall test (0.35 ± 0.41-fold, *p=0.0027*) when compared to tauopathy mice receiving normal drinking water, with values nearing those seen in non-tauopathy controls (**Fig 3C-D**). While we did not observe the same effects of BD treatment in male tauopathy mice, an improvement in primary latency to entry in littermate control mice receiving BD was observed compared to those receiving normal drinking water (0.61 ± 0.27-fold, *p=0.0409,* **Suppl Fig 3C-D**). No significant effects were observed in male mice performing the memory probe recall test (**Suppl Fig 3E**).

**Figure 3.**
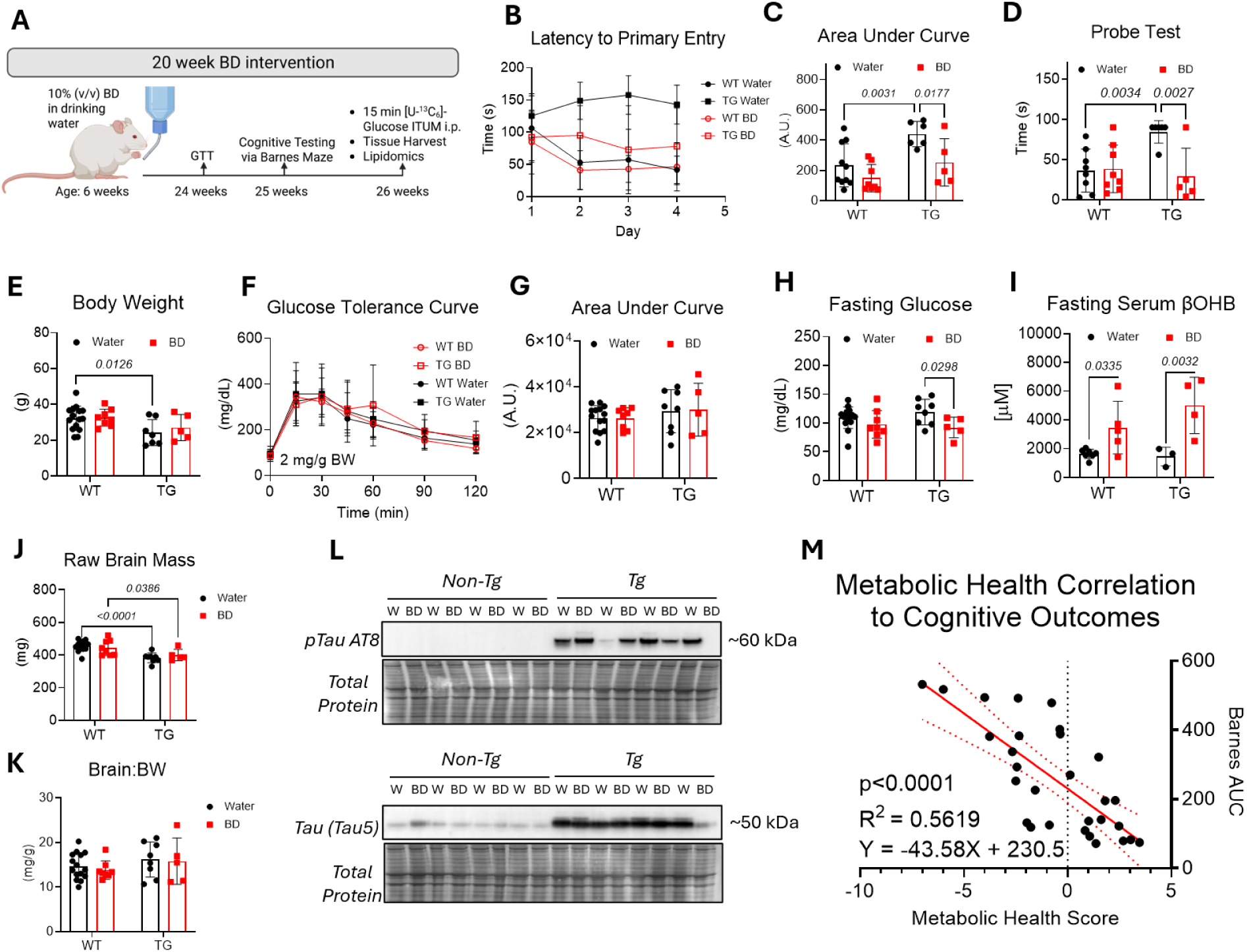
Twenty-week BD treatment improves cognitive outcomes in female tauopathy-transgenic mice. Tauopathy mice and littermate controls were provided with 10% BD in drinking water or normal drinking water for 20 weeks, and cognitive outcomes were assessed. (**A**) Schematic of experimental study design for 20-week BD treatment. (**B**-**C**) Mice were subjected to Barnes Maze behavioral testing following 20-week treatment, with latency to escape hole entry during the memory acquisition phase, days 1-4 (*n*=5-10/group) and corresponding area under the curve for the trials. (**D**) Memory recall probe test on day 5 (*n*=5-10/group) in female mice. (**E**) Effects of transgene and BD treatment on body weight (*n*=5-15/group), and (**F-G**) systemic glucose tolerance test (*n*=5-14/group) performed at 25 weeks of age with corresponding area under the curve for the 120 min test. Following the 20-week intervention, mice were euthanized. (**H)** Effects of 20-week BD treatment on blood glucose values following 18 h fast (*n*=5-15/group) and (**I)** serum βOHB following 18 h fast while consuming the BD-treated water (*n*=3-7/group). (**J-K**) Effect of the BD treatment on raw brain mass (*n*=5-15/group) and brain mass normalized to body weight (*n*=5-15/group) in female mice. (**L**) Immunoblot showing the effect of 20- week BD treatment on phosphorylated human Tau (AT8) and total Tau protein in forebrain regions (*n*=4/group). (**M**) A metabolic health score was constructed following correlation analyses with Barnes Maze AUC, and Z-scores were transformed to plot a linear regression of MHS vs cognitive outcomes. Data are presented as mean ± SD, and group sizes are as indicated. P-values indicated when p<0.10 following two-way ANOVA and Tukey’s post-hoc test. BD = 1,3-butanediol, ITUM = isotope tracing untargeted metabolomics, A.U. = arbitrary units, βOHB= β-hydroxybutyrate, BW = body weight, WT = non-tauopathy control mice, TG = tauopathy mice, MHS = metabolic health score.

Metabolic abnormalities exacerbate cognitive dysfunction^9,15,30^, and while tauopathy mice in both water- and BD-administered groups showed ∼20% lower body mass than littermate controls, there were no effects of BD treatment on body weight (**Fig 3E**). To determine whether variations of glucose homeostasis were linked to the effects of BD treatment in standard water- or BD-treated tauopathy and control mice, we performed glucose tolerance testing (GTT) and observed no effects of BD treatment on systemic glucose tolerance in female mice (**Fig 3F-G**). However, reduced fasting blood glucose was uniquely observed in BD-maintained tauopathy mice [22.5 ± 8.4% (SEM) compared to water-administered tauopathy mice, *p=0.0298;* **Fig 3H**]. BD-treated mice were maintained on BD treatment during fasting, and as expected, the BD treatment increased 18h fasted circulating βOHB in both tauopathy mice and littermate controls more than two-fold (**Fig 3I**). Although overall brain mass was ∼14% lower in tauopathy female mice, this effect was lost when normalized to total body weight, and there was no effect of BD treatment on raw or normalized brain mass in female mice (**Fig 3J-K**). Following the 20-week study, we assessed the effects of BD-treatment on Tau hyperphosphorylation and total Tau protein in forebrain regions and observed no differences between BD and normal water controls (**Fig 3L**).

To predict Barnes Maze AUC outcomes from metabolic indices, we constructed a Metabolic Health Score in female mice by generating correlations with measurements from the 20-week study (**Fig 3M**). Based on calculated Pearson’s correlations, r, we identified a significant positive correlation between Barnes Maze AUC and fasting blood glucose (higher glucose correlated with a poorer learning outcome; r = 0.39, *p=0.038*), and negative correlations with body weight (r = -0.44, *p=0.018*), brain mass normalized to tibia length (r = -0.60, *p=0.001*), heart mass normalized to tibia length (r = -0.44, *p=0.018*), and fasting βOHB concentration (higher ketone correlated with a more favorable learning outcome; r = -0.29, *p=0.13*, **Suppl Table 2, Suppl Fig 4**). These variables were transformed into Z-scores to load a Metabolic Health Score (MHS) = Z_BW_ + Z_brain/TL_ + Z_heart/TL_ + Z_fasting BOHB_ – Z_fasting BG_ to predict Barnes Maze AUC. Linear regression yielded a R^2^=0.56 to predict cognitive outcomes based on MHS (p<0.0001, **Fig 3M**).

As observed with the body weights of females, tauopathy male mice in both water- and BD-administered groups showed ∼20% lower body mass than non-tauopathy littermate controls, but there were no significant effects of BD treatment on body weight (**Suppl Fig 3F)**. Similar to female mice, tauopathy mice showed lower overall brain weight that was lost when normalized to total body weight (**Suppl Fig 3G-H**). Unlike BD-treated female tauopathy mice, BD treatment modestly improved glucose tolerance in male tauopathy mice (but not littermate controls, **Suppl Fig 3I-J**) and there were BD-associated, non-significant diminutions in fasting glucose in both tauopathy mice and littermate controls (**Suppl Fig 3K**). BD treatment increased fasting circulating βOHB in both tauopathy male mice and littermate controls approximately two-fold (**Suppl Fig 3L**). Pearson’s correlations for variables significantly correlated with Barnes Maze AUC scores were found to correlate significantly with body weight (r = -0.52, *p=0.01*), brain mass normalized to tibia length (r = -0.63, *p=0.001*), heart mass normalized to body weight (r = 0.61, *p=0.002*), and fasting βOHB concentration (r = -0.43, *p=0.*039, **Suppl Table 3, Suppl Fig 4**). Unlike in the female mice, we did not observe a significant correlation between Barnes Maze AUC and fasting blood glucose in male mice (*p=0.20*, **Suppl Table 3, Suppl Fig 4**). As such, we constructed a male-specific MHS where MHS = Z_BW_ + Z_brain/TL_ + Z_fasting BOHB_ – Z_heart/BW_ to plot against Barnes Maze AUC. We found this health score to have a significant (*p<0.0001*) linear regression with R^2^=0.75 from our resulting equation, Y = -39.03X + 207.6 (**Suppl Fig 3M**).

### 3.3 Changes in Glucose Metabolism are Associated with Cognitive Function

Impaired brain glucose metabolism is a hallmark feature of progressive neurodegeneration in AD-related dementias^10,31^. To determine how BD treatment modulates brain glucose utilization in rTg4510 tauopathy mice, we performed isotope tracing untargeted metabolomics (ITUM). Twenty-six-week-old female and male tauopathy and littermate control mice were administered BD- or water-control for 20 weeks. Mice were fasted for 18 h, then 5 µmol/g BW [U-^13^C_6_]-glucose was administered via i.p. injection. Fifteen minutes following administration, forebrains were excised and freeze-clamped for analyses. In forebrain regions of male and female brains, we traced glucose-derived ^13^C into metabolites related to glycolysis, biosynthetic pathways, TCA cycle, and cataplerotic fates (**Fig 4A**). A modest increase in ^13^C-enrichment of 2-hydroxyglutarate (2-HG) was observed in female tauopathy brains of water-administered controls and was the only metabolite with significant differences in ^13^C-enrichment; however, this effect was abrogated in BD-treated tauopathy mice (∼53% reduction in ^13^C-enrichment compared to water-treated tauopathy mice, *p=0.0168*; **Fig 4B**). Previous studies identified L-2-hydroxyglutarate as a redox-responsive metabolite that is formed under conditions of reductive stress^32^; our finding suggests that tauopathy-transgenic mice have elevated ^13^C-enrichment compared to non-tauopathy controls and that BD treatment normalizes glucose-derived ^13^C-enrichment of this metabolite.

**Figure 4.**
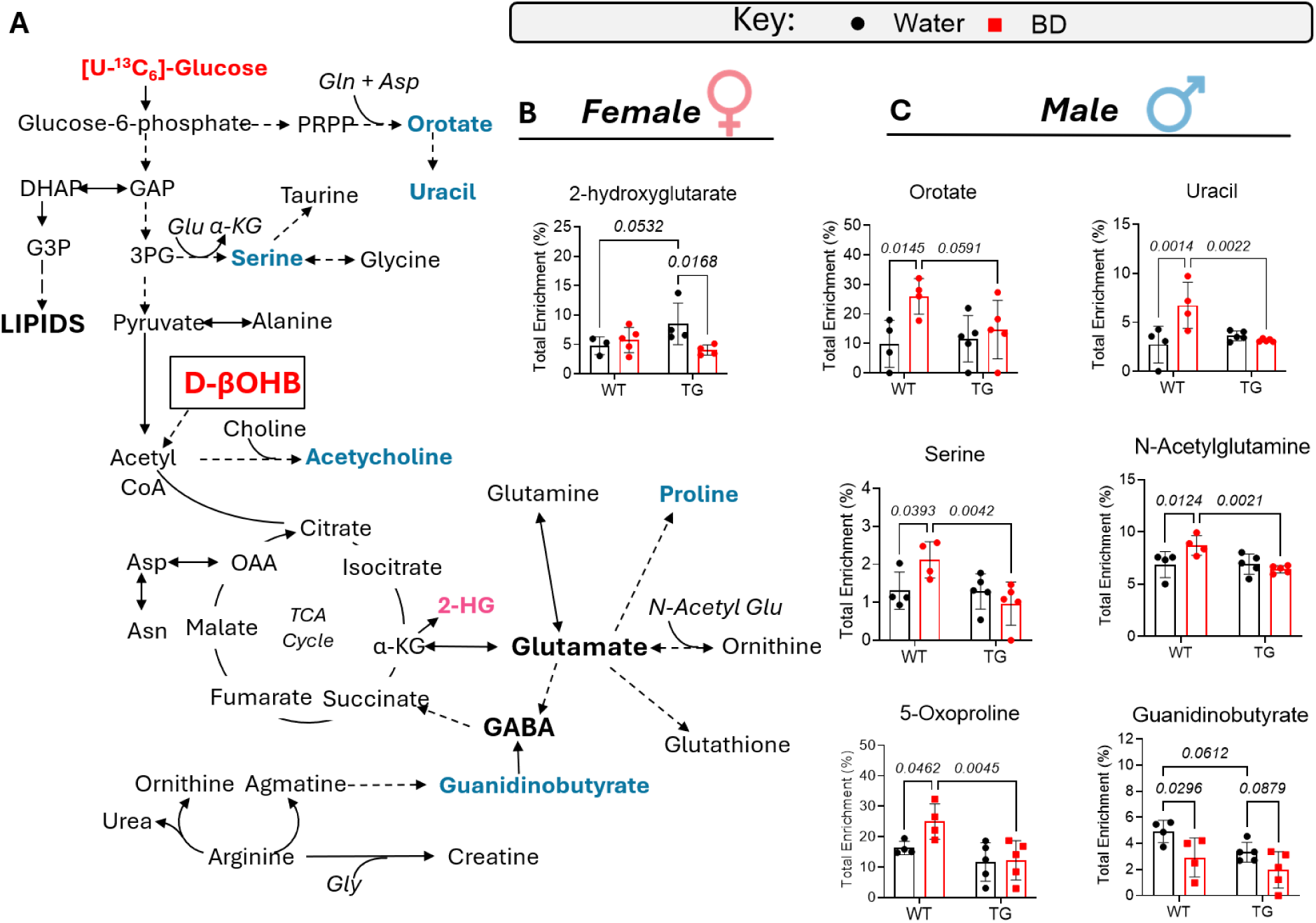
Effects of twenty-week BD treatment on acute glucose utilization in forebrains of mice. Following 20-week BD treatment, male and female mice underwent ITUM with [U-^13^C_6_]-glucose delivered via I.P. injection. Total ^13^C enrichment from glucose-derived carbon was measured in metabolites from freeze-clamped, forebrain sections of mice collected 15-min after glucose administration. (**A**) Schematic of glucose metabolic pathway and corresponding metabolites significantly changed in female (pink) (**B**) or male (blue) (**C**) forebrains following the tracing study. Data are expressed as mean ± SD, *n*=3-5/group. P-values indicated when p<0.10 following two-way ANOVA and Tukey’s post-hoc test. GAP = glyceraldehyde-3-phosphate, DHAP = dihydroxyacetone phosphate, G3P = glycerol-3-phosphate, PRPP = phosphoribosyl pyrophosphate, Gln = glutamine, Asp = aspartate, βOHB= β-hydroxybutyrate, 3PG = 3-phosphoglycerate, Glu = glutamate, Gly = glycine, α-KG = α-ketoglutarate, Asn = asparagine, GABA = gamma-aminobutyric acid, OAA = oxaloacetate, 2-HG = 2-hydroxyglutarate, TCA = tricarboxylic acid, WT = non-tauopathy control mice, TG = tauopathy mice

In BD-treated male mice, there were a greater number of metabolites with significant differences in glucose-derived ^13^C-enrichment of forebrain regions. We traced the fate of ^13^C-glucose into metabolites linked to glycolysis, biosynthetic pathways, TCA cycle, and cataplerotic metabolites. In non-tauopathy male mice, 20 weeks of BD treatment was associated with increased glucose-derived ^13^C-enrichment in orotate (*p=0.0145*), uracil (*p=0.0014*), serine (*p=0.0393*), N-acetylglutamate (*p=0.0124*), and 5-oxoproline (*p=0.0462*) in non-tauopathy control animals, while enrichment in guanidinobutyrate was less than in normal water controls (*p=0.0296*, **Fig 4C**). Elevated ^13^C-enrichment in orotate and uracil could suggest greater *de novo* pyrimidine flux, which could confirm the importance of pyrimidines in brain function, as previously reviewed^33^, while increased brain abundance of N-acetylglutamate is consistent with balanced nitrogen buffering^34^. These BD effects were abrogated in BD-treated tauopathy mice. In male tauopathy brains, significantly lower ^13^C-enrichment was observed in alanine, aspartate, and GABA when compared to non-tauopathy controls (**Suppl Fig 5A-H**). There was little observed effect of BD treatment on glucose-derived ^13^C-enrichment in males. Taken together, the cognitive benefits of BD treatment in male mice promote glucose-derived ^13^C-enrichment of metabolites associated with brain biosynthesis, but these effects are blunted in male mice with tauopathy, which did not derive a cognitive benefit from BD treatment.

### 3.4 Biological Sex, Tau Hyperphosphorylation, and BD Treatment have Minimal Influence on Brain Lipidome

Neurodegenerative diseases, including AD, are known to influence lipid architecture in the brain^35^. As such, we mapped major lipid classes in the forebrain that could be influenced by glucose-derived precursors or by brain uptake of lipids from circulation, focusing on key phospholipids, cholesterol, and sphingolipids (**Fig 5A**). To determine the effects of tauopathy and BD treatment on the brain lipidome, we performed shotgun lipidomics on forebrains from 26-week-old tauopathy and littermate control mice administered BD or normal water for 20 weeks. Analysis of class-level lipid abundances revealed that BD treatment had minimal impact on the overall lipidome in both male and female brains, regardless of genotype (**Fig 5B**). The absence of major treatment effects is potentially attributable to high intragroup variability observed across lipid species (**Suppl Fig 6A**), which likely limits statistical detection of subtle changes. In fact, when correcting for multiple statistical tests (FDR q<0.05), there were no significant changes in abundance of any individual lipid species in female or male brains that aligned with the cognitive benefits we observed (**Suppl Table 4**).

**Figure 5.**
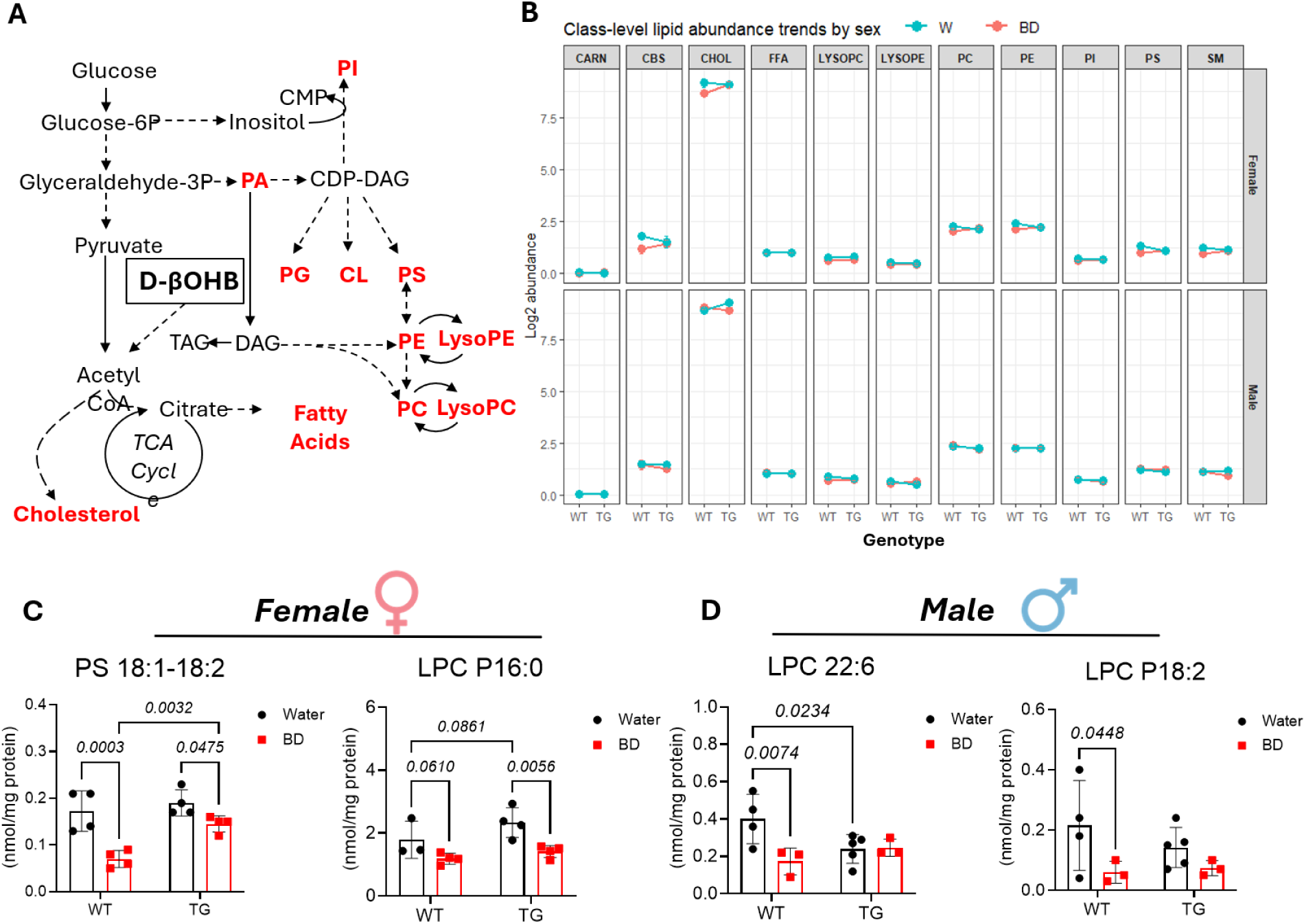
BD treatment influences the brain lipidome of mice in a treatment and genotype-specific manner. (**A**) Schematic depicting lipid pathways queried in the brain. (**B**) Absolute concentrations of lipid species were analyzed using RStudio to highlight changes in class-level lipid abundances based on biological sex, genotype, and water treatment. Concentrations of selected phospholipid species that were significantly changed in groups with observed cognitive benefits, before FDR cutoff, are shown in the female brain (**C**) and in the male forebrain (**D**) following 20-week BD treatment. Data are expressed as mean ± SD, *n*=4/group. Group analyses performed using two-way ANOVA model with post-hoc comparisons performed using estimated marginal means and p-values adjusted using Benjamini-Hochberg false-discovery rate method. P-values in **C** and **D** indicated when p<0.10 following two-way ANOVA with Tukey’s post-hoc test, and before false discovery rate cutoff, where q>0.05. WT = non-tauopathy control mice, TG = tauopathy mice, PC = phosphatidylcholine, PE = phosphatidylethanolamine, PA = phosphatidic acid, PI = phosphatidylinositol, PS = phosphatidylserine, SM = sphingomyelin, CBS = cerebroside, FFA = free fatty-acid, LPC = lysoPC, LPE = lysoPE, Car = acylcarnitine

A few individual lipid species showed significant alterations before the FDR cutoff. BD treatment reduced forebrain levels of PS 18:1-18:2 (*p=0.0475*) and LysoPC P16:0 (*p=0.0056*) in female tauopathy mice compared to littermate controls (**Fig 5C**). While higher levels of PS species in the brain are typically associated with positive effects on synaptic function and cognition^36^, the observation that BD treatment lowered PS 18:1-18:2 in brains of both tauopathy (24% reduction) and non-tauopathy (59% reduction, *p=0.0003*) mice when compared to normal water controls may indicate its effect in shifting lipid composition away from unsaturated species that are typically more reactive^37^. In male mice, BD treatment lowered LysoPC 22:6 (*p=0.0074*) and LysoPC P18:2 (*p=0.0448*) in brains of non-tauopathy littermate controls but not in brains of tauopathy mice (**Fig 5D**). LysoPCs have been shown to be elevated in the brains of those with neurodegenerative diseases, potentially contributing to neuroinflammation^38^. While overall LysoPC levels were not significantly changed in any group, our data suggest changes in individual LysoPC species are associated with cognitive benefits of BD treatment.

Additionally, we performed partial least squares discriminant analyses (PLS-DA) of brain lipid species from female and male brains, with corresponding variable importance projection (VIP) scores plots to determine standout metabolites with greatest influence on group separation (**Suppl Fig 6B-E**). In female brains, we found VIP scores >2.5 for phosphatidylcholine (PC) species D18:0-20:1/P18:1-22:6, phosphatidylethanolamine (PE) species P18:2-22:6/D18:1-20:1, and phosphatidylserine (PS) species 18:2-18:2 (**Suppl Fig 6C**). In male brains, we found VIP scores >3 for PC D18:0-22:4/D20:0-20:4/D20:2-20:2, PS P18:0-22:5, and PC D18:0-20:1/P18:1- 22:5 (**Suppl Fig 6E**). Abundances of these lipid species contributed to small, overall group differences; however, none of these individual lipids were significant.

### 3.5 Thirty-week BD Treatment Does Not Improve Cognitive Outcomes in Tauopathy Mice

We also sought to determine whether cognitive impairment could be improved by administering 10% BD in drinking water to tauopathy and non-tauopathy littermate control mice for 30 weeks, beginning at 6 weeks of age, with the study concluding when mice were 36 weeks of age (**Fig 6A**). This is due to the progressive nature of neurofibrillary tangle formation in the tauopathy mouse model, and the inherent nature of many therapeutic interventions to delay but not prevent disease progression in AD and related dementias. Female tauopathy mice weighed significantly less than littermate controls at the start of the study (*p=0.006*) and retained lower body weight throughout the 30-week treatment (**Fig 6B**), but tauopathy was not associated with lower body mass in male mice until after 9 weeks of age (**Suppl Fig 7A**). BD treatment did not influence circulating blood glucose values (**Fig 6C**), but it elevated circulating AcAc (**Fig 6D**) and circulating total βOHB (**Fig 6E**) at all time points throughout the study. Similar effects of BD were measured in males on circulating glucose, AcAc, and βOHB, though BD induced modestly lower ketosis in males compared to females (**Suppl Fig 7B-D**). No differences in circulating ketone bodies were evident in normal water- or BD-treated tauopathy animals when compared to littermate controls (**Fig 6D-E**). We measured the distribution of D- and L-βOHB just prior to BD administration (6 weeks of age), at the midpoint of the BD administration period (18 weeks of age), and near the end of the study (30 weeks of age) and found circulating L-βOHB to account for <2% of all circulating βOHB at 6 weeks of age in all groups, while it also accounts for <1% of all βOHB in normal water controls at 18 weeks and 30 weeks of age (**Fig 6F**). In BD-treated female mice, L-βOHB accounted for >50% of total circulating βOHB in both non-tauopathy and tauopathy females at 18 weeks of age. At 30 weeks of age, L-βOHB accounted for >70% of total circulating βOHB in both non-tauopathy and tauopathy females.

**Figure 6.**
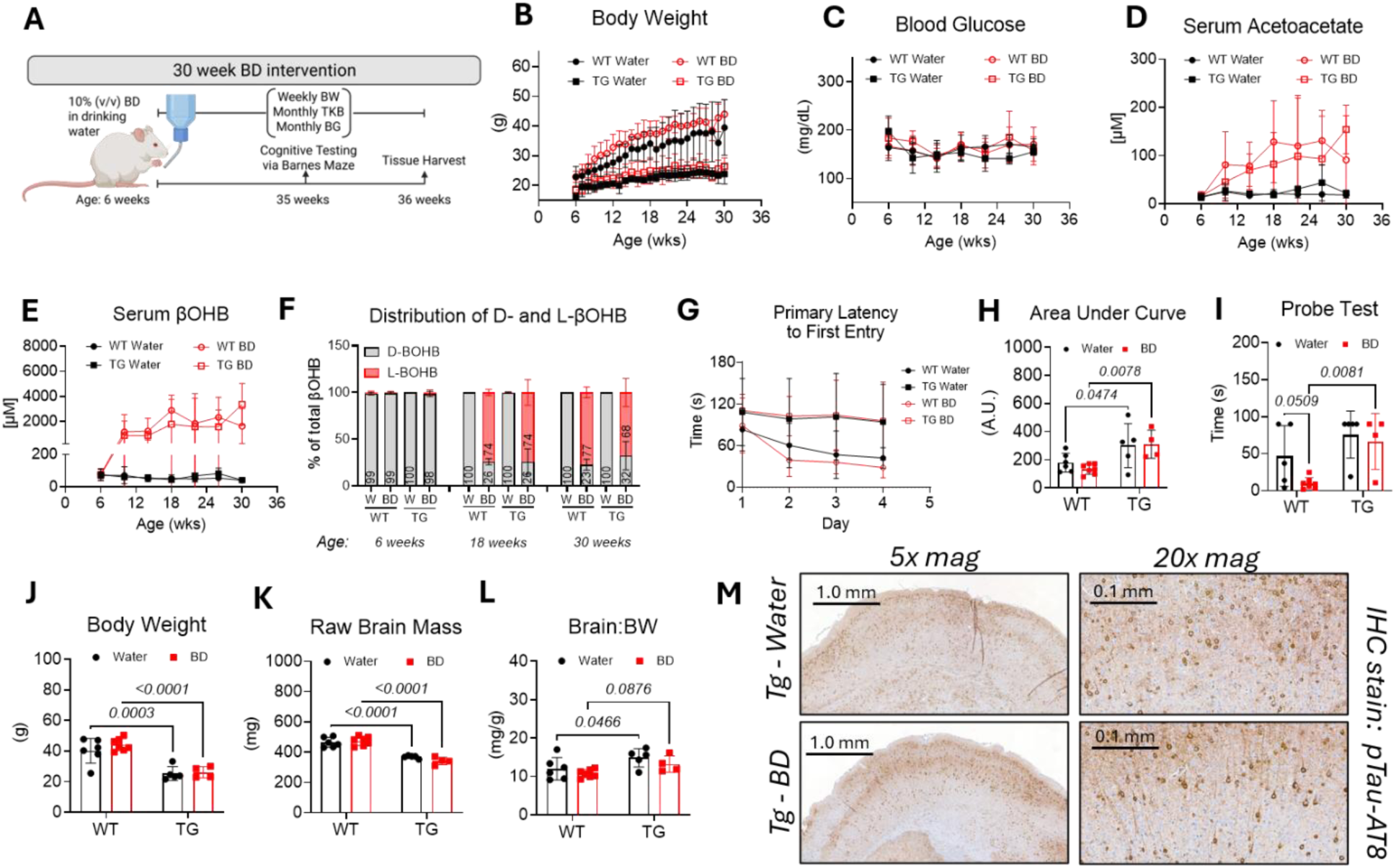
Thirty-week BD treatment has no effect on cognitive outcomes in female tauopathy-transgenic mice. (**A**) Female tauopathy mice and non-transgenic littermate controls were provided with 10% BD in drinking water or normal drinking water for 30 weeks and cognitive outcomes were assessed. Effects of the transgene and BD treatment on body mass (**B**), non-fasted blood glucose (**C**), serum AcAc (**D**), and serum βOHB (**E**) over 30-week treatment. (**F**) Total percentages of D- and L-βOHB in circulation at baseline (6 weeks of age), 12-week treatment (18 weeks of age), and 24-week treatment (30 weeks of age, *n*=3/group). (**G-H**) Mice were subjected to the Barnes- Maze for cognitive testing. Latency to escape hole entry during the memory acquisition phase, days 1-4, and corresponding area under the curve for the trials. (**I**) Latency to escape the hole entrance during day 5, memory probe. Thirty-week BD treatment did not affect body mass (**J**), raw brain mass (**K**), or brain mass normalized to body weight (**L**) in transgenic or non-transgenic female mice at tissue harvest. (**M**) Representative images from immunohistochemistry staining of paired helical filaments in the forebrain region from tauopathy-transgenic mice exposed to BD treatment or normal drinking water. Data are expressed as mean ± SD, *n*=4-7/group unless otherwise indicated. P-values indicated where p<0.10 following two-way ANOVA and Tukey’s post-hoc test. BD = 1,3-butanediol, BW = body weight, BG = blood glucose, βOHB= β-hydroxybutyrate, TKB = total ketone bodies, A.U. = arbitrary units, mag = magnification, IHC = immunohistochemistry, WT = non-tauopathy control mice, TG = tauopathy mice.

We next tested the cognitive impact of a 30-week BD treatment using a Barnes Maze. As expected, tauopathy female mice consuming normal drinking water showed severe impairment (1.67 ± 0.87-fold increase vs non-transgenic, *p<0.05*) in primary latency to escape hole entry during the memory acquisition phase (days 1-4, **Fig 6G-H**) and in the memory probe recall (*p=0.05*, day 5, **Fig 6I**). The same effect was observed in tauopathy male mice consuming normal drinking water (**Suppl Fig 7F-H**; 1.74±0.81-fold increase vs non-tauopathy, *p<0.05*). BD treatment did not confer cognitive benefits to tauopathy or control mice, as measured using the Barnes Maze protocol in female or male mice. No effects of the BD treatment were observed on final body weight or brain mass, and tauopathy mice consuming normal drinking water retained 36.9 ± 7.3% (SEM) lower body mass (*p=0.0003*) and 20.5 ± 2.5% (SEM) lower raw brain mass (*p<0.0001*) than non-tauopathy controls (**Fig 6J-K**). The effect of tauopathy on brain mass was lost when normalized to body weight (**Fig 6L**). In males, we noticed a similar effect of the transgene on body weight, raw brain mass, and brain mass normalized to body weight (**Suppl Fig 7I-K**). Tauopathy female mice consuming normal drinking water exhibited 66.2 ± 14.7% (SEM) lower adiposity than littermate controls (*p<0.001*), with no BD effect (**Suppl Fig 8A**). BD-treated female mice had lower non-fasted glucose values at tissue harvest when compared to normal water controls (**Suppl Fig 8B**); however, there were no significant effects of the transgene or BD treatment on male body fat composition or non-fasted glucose values (**Suppl Fig 8C-D**). Brains from the tauopathy mice were fixed in formalin, paraffin-embedded, and then subjected to immunohistochemical staining of Tau-AT8, with no significant visual effects of BD treatment on tau pathology after 30 weeks in female (**Fig 6M**) or male mice (**Suppl Fig 7L**).

### 3.6 Western Diet Diminishes BD-Induced Ketosis and Prevents Cognitive Benefits in Tauopathy Mice

To determine whether the cognitive benefits of BD are preserved in mice with diet-induced cardiometabolic dysfunction, we next tested the effects of BD on mice maintained on a high-fat (40%kcal fat), sucrose-enriched Western diet (WD), initiated at 6 weeks of age and continued for 20 weeks - the same duration in which BD improved Barnes Maze performance in female tauopathy mice and in male control mice (**Fig 7A**). BD treatment (versus water control) was also initiated at 6 weeks of age. As previously observed, tauopathy mice exhibited a significant reduction in body weight at the start of intervention (**Fig 7B**; *p=0.0438* tauopathy WD+BD vs. non-tauopathy controls on WD+BD), but there were no significant effects of BD treatment throughout the study when comparing to normal water controls on WD after performing a mixed-effects analysis. BD treatment had no effect on random-fed, circulating blood glucose values (**Fig 7C**), and, surprisingly, only modest effects on circulating AcAc (**Fig 7D**) and total βOHB (**Fig 7E**) when compared to normal water controls consuming WD. Following a mixed-effects analysis, WD+BD did not significantly elevate circulating βOHB compared to normal water controls consuming WD in either tauopathy or littermate control mice. At 18 weeks of age (12 weeks into study), the percentage of L-βOHB enantiomer was 59 ± 14.2% (SEM) of total βOHB in the non-tauopathy mice, while it was only 30 ± 13.9% (SEM) in the tauopathy mice (**Fig. 7F**). Neither total βOHB nor the percentage of L-βOHB was significantly different between groups. To determine if BD treatment improved cognitive outcomes in mice fed the WD, we performed Barnes Maze testing and found that, as expected, tauopathy mice showed delayed memory acquisition compared to littermate controls. However, no cognitive benefits of BD treatment were observed in either tauopathy or littermate control mice (**Fig 7G-H**). Gross mass of brains harvested from tauopathy mice was diminished compared to littermate controls, and no effect of BD treatment was observed (**Fig 7I**); however, this effect was lost when normalized to body weight (**Fig 7J**). GTT of mice maintained on WD showed, as expected, generally higher AUCs for WD mice compared to chow-fed mice we previously reported, and while there was a modest (non-statistically significant) improvement of whole-body glucose disposal in female tauopathy mice compared to littermate controls, there was no effect of BD in either genotype (**Suppl Fig 9A-C**).The BD treatment group exhibited a reduction in 18h fasting blood glucose in tauopathy mice (*p=0.0134*; **Fig 7K**). While fed state circulating ketones were not significantly increased in WD+BD mice following an 18h fast, the WD+BD significantly increased circulating βOHB to 4.3 ± 3.6 mM in non-tauopathy littermate mice compared to 0.97 ± 0.80 mM in normal water controls (*p=0.056*), while it elevated circulating βOHB to 5.4 ± 2.5 mM in tauopathy mice compared to 1.3 ± 0.4 mM in normal water controls (*p=0.034*; **Fig 7L**). L-βOHB was detected only in mice receiving BD treatment while fasting, contributing ∼50% of the circulating βOHB pool in both tauopathy mice and littermate controls (**Fig 7M**).

**Figure 7.**
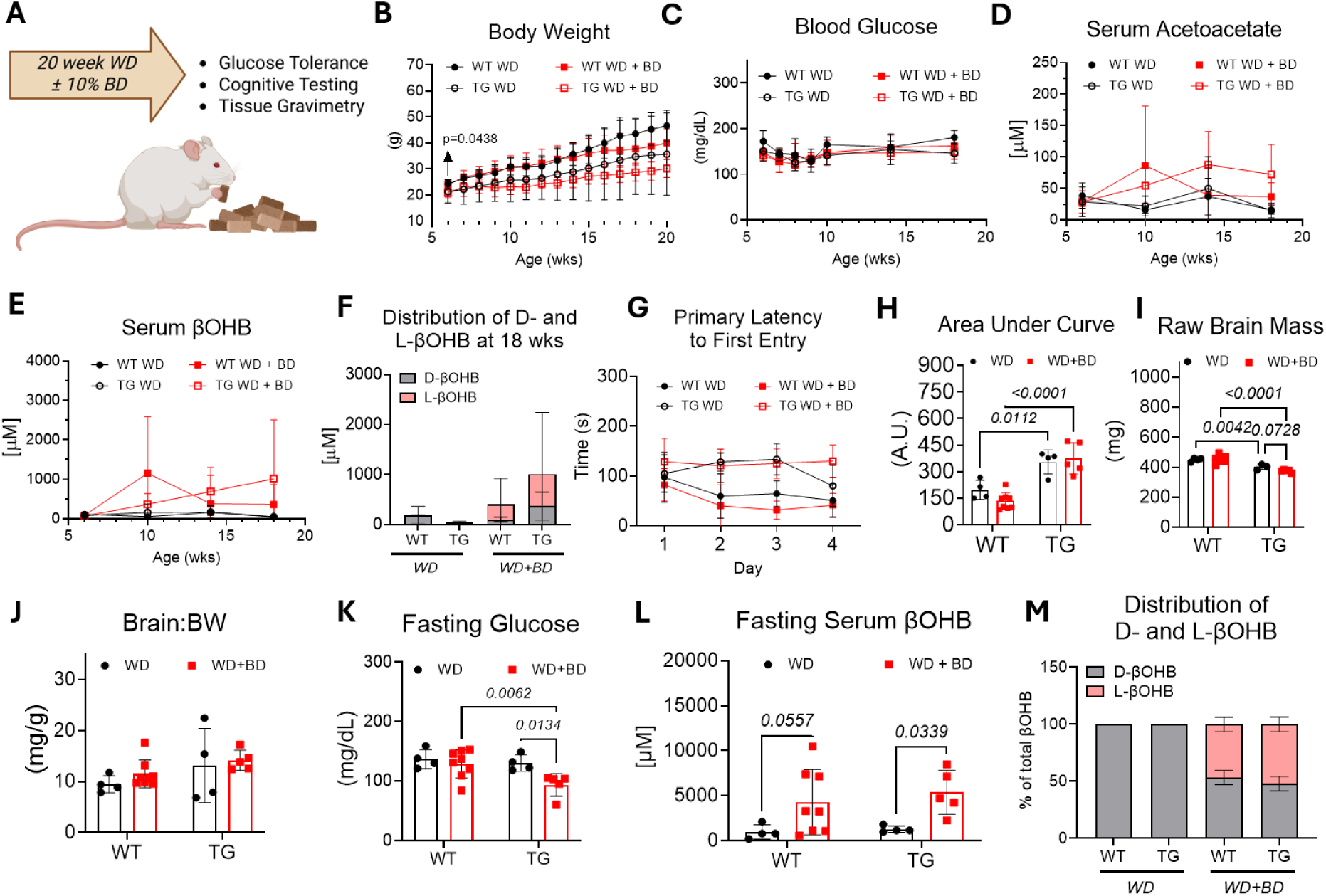
High-fat, Western diet abolishes cognitive benefits of BD treatment in female mice. (**A**) Female tauopathy mice and non-transgenic littermate controls were subjected to a 20-week high-fat Western diet (WD) while consuming 10% BD in drinking water or normal drinking water. Cognitive function was assessed using a Barnes Maze before tissues were harvested for analyses. (**B-C**) Effects of the dietary intervention, with or without BD treatment, on body weight and fed blood glucose throughout the study. (**D-F**) BD treatment effects on serum acetoacetate, serum βOHB, as well as the concentration and distributions of D-and L-βOHB at study midpoint (12- week treatment, 18 weeks of age). (**G-H**) Latencies to escape hole entry during the memory acquisition phase, days 1-4 of Barnes Maze, with corresponding area under the curve. Following cognitive testing, mice were euthanized, and the raw brain mass (**I**) of female mice was plotted with brain mass normalized to body weight (**J**). 18-hour fasted blood glucose (**K**) and fasted serum βOHB (**L**), along with enantiomeric distribution of βOHB, shown in (**M**). Data are expressed as mean ± SD, *n*=4-8/group. P-values indicated when p<0.10 following two-way ANOVA and Tukey’s post-hoc test. WD = high-fat, Western diet, BD = 1,3-butanediol, A.U. = arbitrary units, βOHB= β-hydroxybutyrate, WT = non-tauopathy control mice, TG = tauopathy mice.

In male mice, there were no genotype or BD-induced effects on rate of body weight increase in WD-fed mice, and tauopathy mice had lower body weights that non-tauopathy controls (**Suppl Fig 10A**). Furthermore, BD treatment had no significant effects on circulating glucose, AcAc, or βOHB in either tauopathy or non-tauopathy control mice (**Suppl Fig 10B-D**). At 18 weeks of age (12 weeks into study), the percentage of L-βOHB was 28 ± 15.5% (SEM) and 68 ± 4.1% (SEM) of total circulating βOHB in non-tauopathy and tauopathy mice receiving the BD treatment, respectively (**Suppl Fig 10E**), although there were no significant differences among groups. Importantly, there were no effects of BD treatment on cognitive outcomes or brain mass in male mice fed WD (**Suppl Fig 10F-I**). Additionally, there were no effects of BD treatment on 18h fasted blood glucose (**Suppl Fig 10J**), blood βOHB (**Suppl Fig 10K**) or systemic glucose tolerance in these mice (**Suppl Fig 9D-F**). Surprisingly, in male mice receiving the BD treatment while fasting, we observed lower L-βOHB distribution than in female mice. The contribution of L-βOHB was only ∼3% of total βOHB measured in fasted, tauopathy mice and only ∼16% of total βOHB in fasted, non-tauopathy mice (**Suppl Fig 10L**).

When considering variables that correlate with cognitive benefits in WD-fed mice with or without BD treatment, fasting blood glucose [r = -0.065 (*p=0.761*) and -0.582 (*p=0.006*) for male and female, respectively] and systemic glucose tolerance [r = -0.156 (*p=0.466*) and -0.529 (*p=0.014*) for male and female, respectively] were negatively correlated with Barnes Maze AUC (**Suppl Fig 11, Suppl Tables 5-6**). In WD-fed mice, chronic ketosis [r = 0.396 (*p=0.045*) and 0.278 (*p=0.223*) for male and female, respectively, as measured by βOHB AUC over 18 wks] and Barnes Maze AUC were positively correlated (**Suppl Fig 11**). Unlike chow-fed mice, these observations imply that in WD-fed mice, impaired glucose metabolism and lower sustained concentration of βOHB throughout the 20-week treatment period may predict better cognitive outcomes.

### 3.7 Ketogenic diet does not ameliorate cognitive impairment in a tauopathy mouse model

Unlike mice maintained on normal chow for 20 weeks, BD delivery did not improve cognition in tauopathy mice or littermate controls fed WD. However, because BD-induced ketosis was relatively blunted in mice maintained on WD, we next determined whether a distinct approach to augment ketogenesis would increase ketosis and improve cognitive outcomes. Endogenous ketogenesis was induced by weekly oscillations between WD and a zero-carbohydrate, very high-fat ketogenic diet (KD), an approach previously employed to improve cognitive and health span outcomes in mice^39^ (**Fig 8A**). Diets were cycled weekly for 20 weeks, beginning at 6 weeks of age. Body weights were reduced in male and female mice consuming the cyclical dietary strategy when compared to cohorts consuming only WD, and as observed on other regimens, male and female tauopathy mice showed generally lower body weights than littermate controls at the end of the 20 week regimen, by 16.9 ± 7.7% (SEM, *p=0.1007*) and 31.2 ± 7.4% (SEM, *p=0.0014*), respectively (**Fig 8B-C**). Only during weeks when consuming KD, circulating AcAc and total βOHB were markedly elevated in all groups (**Fig 8D-E**), and given that ketosis was endogenously provoked, the percentage of L-βOHB from ketogenic diet was <3% in all groups treated (**Fig 8F**). While on a KD, circulating βOHB was reduced in female tauopathy mice compared with non-tauopathy controls (*p=0.0450*). Circulating blood glucose values were not significantly different among the groups maintained on this regimen (**Fig 8G**).

**Figure 8.**
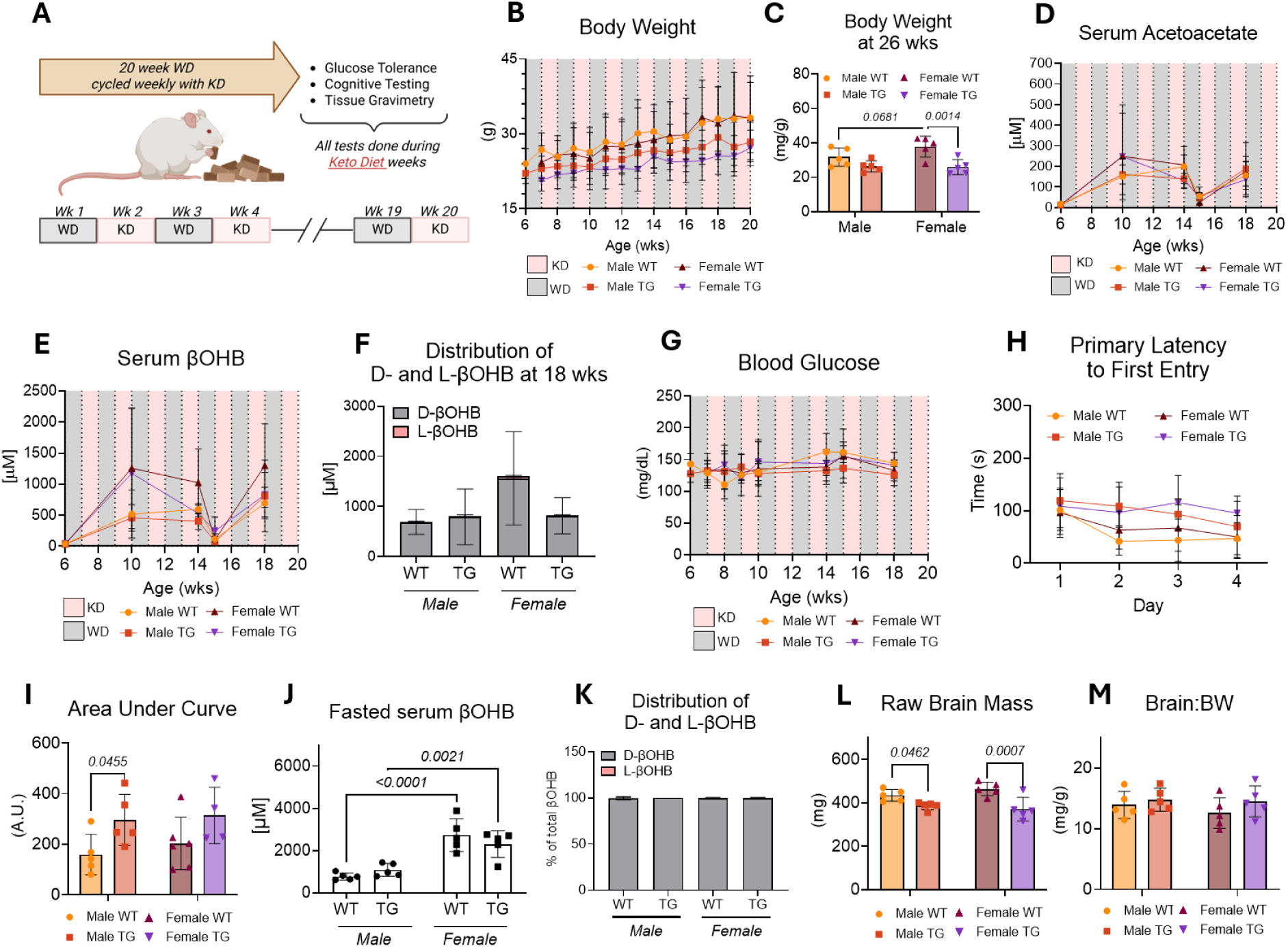
Weekly cycling of a very high-fat, zero-carbohydrate ketogenic diet (KD) with high-fat, Western diet (WD) does not improve cognitive outcomes in tauopathy mice. (**A**) Male and female tauopathy mice and littermate controls, were fed high-fat, Western diet (WD) that cycled weekly with a high-fat, very low-carbohydrate ketogenic diet (KD) for a total of 20 weeks. All measurements were performed during the ketogenic diet weeks unless otherwise indicated. (**C-D**) Body weights throughout the study and at the conclusion of the study (aged 26 weeks). (**D-E**) Random fed serum acetoacetate and βOHB throughout the study. (**F**) Distributions of D-and L-βOHB midway through the study (12 weeks into the study, 18 weeks of age). (**G**) Random fed blood glucose throughout the study. (**H-I**) Barnes Maze cognitive testing, with latency to escape hole entry plotted for memory acquisition phase, days 1-4, and corresponding area under the curve. Following cognitive testing, mice were fasted for 18 h and then euthanized. (**J-K**) Fasting serum βOHB and enantiomeric distribution of D/L βOHB. (**L-M**) Raw brain mass, and brain mass normalized to body weight. Data are expressed as mean ± SD, *n*=4-8/group. P-values indicated where p<0.10 following two-way ANOVA with Tukey’s post-hoc test. βOHB= β-hydroxybutyrate, WD = high-fat, Western diet; KD = ketogenic diet; WT = non-tauopathy control mice; TG = tauopathy mice

At the end of the 20-week regimen, cognitive performance was assessed using the Barnes maze while mice were consuming KD. As expected, tauopathy mice showed significant impairment in memory acquisition, more pronounced in males (*p=0.0455*) than females (*p=0.1037*, **Fig 8H-I**). The cycling KD regimen did not show differences between tauopathy and littermate control mice in baseline glycemia (**Suppl Fig 12A, 12D**), systemic glucose tolerance (**Suppl Fig 12B-C, 12E-F**), or fasted total βOHB, although females showed higher βOHB than males (**Fig 8J**). When considering the distribution of D-βOHB and L-βOHB in fasted mice, no group showed more than 1% contribution of L-βOHB to total βOHB measured (**Fig 8K**). Raw brain masses were lower in tauopathy mice than littermate controls, but this effect was lost after normalization to body weight (**Fig 8L-M**).

For mice consuming the WD cycling weekly with KD, we correlated various measurements throughout the 20-week study with Barnes Maze AUC scores (**Suppl Fig 11**). In female mice, we observed a negative correlation between fasting blood glucose and Barnes Maze AUC (r = -0.326, *p=0.359*), as well as a negative correlation between fasting βOHB and Barnes Maze AUC (r = - 0.617, *p=0.077*), together indicating that higher substrate concentrations correlated with better cognitive outcomes (**Suppl Table 5**). Male mice showed no correlation between fasting blood glucose and Barnes Maze AUC (r = 0.068, *p=0.852*) while a positive correlation between fasting βOHB and Barnes Maze AUC (r = 0.428, *p=0.217*) was observed (**Suppl Table 6**), supporting that fasting glucose may have minimal association with cognitive outcomes while lower fasting βOHB concentrations associate with better cognitive outcomes in mice cycling weekly between WD and KD.

## 4. Discussion

In this study, we evaluated the delivery of an exogenous ketone body precursor, (R,S)-1,3-butanediol (BD), to forebrain-specific, tauopathy mice and non-tauopathy littermate controls. We showed that 10% BD treatment promoted rapid, euglycemic ketosis in a unique physiological condition that promotes sustained circulatory elevations of the L-βOHB enantiomer. Cognitive benefits of BD were observed only in mice consuming standard chow and before severe cognitive decline had progressed, suggesting a therapeutic window may exist for intervention. BD treatment also altered glucose-derived carbon enrichment into biosynthetic pathways in the brain while reducing accumulation of lipids associated with neuroinflammation. BD-derived cognitive benefits were lost in advanced disease and during consumption of WD. Similarly, intermittent endogenous ketosis induced by cycling WD with a KD did not improve cognition. Together, these findings highlight that responsiveness to ketogenic interventions depend on metabolic health, disease progression, and biological sex.

Both acute and chronic exposure to 10% BD elevated circulating βOHB to physiological concentrations (∼2 mM) while markedly increasing the ratio of βOHB:AcAc in circulation beyond levels typically observed with normal fasting or ketogenic diets (**Fig 1**). Because oxidation of D-βOHB requires fewer mol NAD+ per acetyl-CoA generated than glucose, BD treatment could alter mitochondrial redox status and glucose utilization^40,41^. Modulating this ratio could affect intracellular TCA cycle flux and intracellular activity from greater mitochondrial NAD^+^ availability^42^, where oxidation of D-βOHB benefitted survival and memory in the hAPPJ20 AD mouse model^43^. However, while consuming BD, L-βOHB was predominant in circulation (>60% of the D+L total βOHB pool). This dominance in circulation results from slower, BDH1-independent clearance of L-βOHB compared to the D-βOHB enantiomer. As such, redox changes typically associated with ketogenic interventions may be less meaningful than expected since only D-βOHB is oxidized via BDH1^44,45^. It remains unclear what extent the L-enantiomer executes its functions as a metabolic fuel versus a signaling molecule^46,47^. Recently, it was reported that pharmacologic augmentation of NAD^+^ via P7C3-A20 reversed tau phosphorylation and oxidative stress in brains of the 5xFAD mouse model of AD^48^, and a recent preprint suggests that neuronal benefits of βOHB could be independent of D-βOHB oxidation, as the less avidly-oxidized L-βOHB enantiomer ameliorated pathophysiological consequences of tau in another tauopathy mouse model^49^. Furthermore, chronic BD treatment reduced βOHB AUC in male mice during a KTT, suggesting adaptive enhancement of ketone body disposal or diminished endogenous ketogenesis in BD-treated mice. Although effects were less pronounced in females, these findings support the possibility that chronic BD exposure modifies systemic ketone handling and neuronal redox balance. Together, this could imply a synergistic effect of elevating both D- and L-βOHB in neurons.

In our study, biological sex, treatment duration, and diet all affected cognitive outcomes (**Fig 3**, **Fig 6**). While a sustained, euglycemic, physiological ketosis was achieved in both male and female mice maintained on standard chow over the course of 30-week BD treatment, there were no cognitive benefits observed and no visual effects on tau pathology from brain immunohistochemistry. Due to the progressive nature of tau hyperphosphorylation and NFT formation in the rTg4510 model, it is likely that the age of mice following a 30-week intervention (aged 36 weeks) was outside of a therapeutic window for metabolic effects of BD treatment. This is supported by our findings that a 20-week BD treatment resulted in significant benefits to female, tauopathy mice and to male, control mice when compared to respective, normal water controls. It is also possible that the 30 week BD treatment could increase expression of monocarboxylate transporters^50^ or influence epigenetic remodeling^51^ in the brain compared to the shorter, 20 week treatment duration and that this adaptation forfeits benefits observed in the shorter treatment.

Since neuronal glucose hypometabolism is hallmark of neurodegenerative disease^10^ and that ketone bodies influence glucose metabolism^11^, we performed ITUM to test the effects of chronic BD treatment on brain glucose utilization (**Fig 4**). Our key findings suggest a biological sex-dependent effect driven both by the transgene and BD treatment. Previous reports link tauopathy to mitochondrial dysfunction^52^ and oxidative stress^53^ in the brain, with elevations in the specific metabolite 2-hydroxyglutarate being associated with neurodegeneration in mice^54^. We found that in females, BD treatment appeared to correct a tau-driven elevation in glucose-derived ^13^C-enrichment of 2-hydroxyglutarate in the brain. Reductions in 2-hydroxyglutarate have been reported to mitigate neuronal dysfunction in *Drosophila* species^55^, supporting a neuroprotective role of BD treatment in tauopathy mice. BD also lowered ^13^C-enrichment in glutathione of female brains, while overall labeling of glycolytic and TCA cycle metabolites remained unaffected (data not shown). Together, these changes could suggest that BD treatment may reduce the need for diverting glucose toward antioxidant pathways in the brain.

In males, BD treatment augmented glucose-derived anabolic activity in the brain (**Fig 4**). BD treatment did not restore cognitive function in male tauopathy mice; however, most BD-induced changes we observed in forebrain glucose utilization were associated with cognitive benefits observed in non-tauopathy male mice. In mice that showed cognitive benefits, glucose-derived ^13^C-enrichment of orotate, uracil, serine, and 5-oxoproline suggests that βOHB does not just serve as a replacement for glucose oxidation in the brain, but instead may redirect glucose utilization for biosynthetic purposes that promote synaptic function and preservation of plasticity^56^ in neuroinflammatory settings like AD. Interestingly, a UMP-enriched diet, which increases uridine availability and expands uracil pools, rescued cognitive decline in an impoverished rat model^57^, further linking our observations to brain health. Also consistent with our findings, studies associate higher serine brain levels with beneficial cognitive outcomes^58^, and a recent genome-wide association study showed a causative association between fasting plasma 5-oxoproline and cognitive function in humans^59^. Interestingly, a recent study showed that tauopathy mice shift glucose-derived ^13^C-enrichment of glutamate and GABA in the brain, promoting a hyperexcitability in neurons of the P301S tauopathy mouse model^60^. However, we did not observe any effects on glucose-sourced ^13^C-enrichments of GABA and glutamate using our mouse model of tauopathy (**Suppl Fig 5I**). These data support that ketosis may benefit cognitive outcomes through changes in brain glucose utilization.

Neurodegenerative diseases, including Alzheimer’s Disease, are associated with alterations in the brain lipidome^35,61^. Our data revealed selective lipid remodeling influenced by biological sex, genotype, and BD treatment. Overall lipidomic changes were modest, but female tauopathy mice treated with BD showed reductions in LysoPCs and selective phospholipid species associated with neuroinflammatory states^36,38,61^ (**Suppl Fig 6**). Similarly, while long-chain acylcarnitine species have been implicated as predictive biomarkers for AD^62^, we saw a modest reduction in the acylcarnitine 18:2 when mice were given BD treatment (**Suppl Table 4**). Lipidomic changes were less pronounced in brains of male mice; however, tauopathy male mice showed ∼30% increase in brain free cholesterol, which was normalized by BD treatment (**Suppl Table 7**). Dysregulated cholesterol metabolism in the brain is associated with inhibited synaptic vesicle formation^63^, suggesting that BD treatment may benefit neurotransmission, but it remains unclear whether brain cholesterol in tauopathy mice is a compensatory response for pathology or a direct result of such. Corresponding to cognitive improvements of BD treatment in non-tauopathy male mice were elevations in PI 16:0-18:0 and PS P20:0-22:5, which could indicate that BD modulates neuroinflammatory states^36,64^. Taken together, BD treatment could minimize lipid signatures associated with neurodegeneration and metabolic stress conditions.

To integrate systemic metabolic changes with cognitive outcomes, we developed a composite metabolic health score (MHS) incorporating multiple physiological variables. While brain mass had the strongest correlation with Barnes Maze performance (**Suppl Fig 4**, **Suppl Tables 2-3**), a single variable often fails to capture the organism’s holistic metabolic state. By integrating multiple physiological parameters into a single score, we accounted for the dynamic contributions of tau progression and BD intervention to cognitive outcomes. We found that a simple additive model using raw Z-scores outperformed weighted variants, suggesting a synergistic, rather than hierarchical, contribution of metabolic indicators to cognitive outcomes. The strong correlations observed between MHS and Barnes Maze performance (R^2^ = 0.56 and 0.75 for females and males, respectively) reinforce that the cognitive benefits of BD treatment are due to an integrated shift toward a metabolically favorable state rather than being a sole result of increased ketone availability. While our modeling is not intended to serve as a clinical predictor, the strong correlation with functional outcomes in the rTg4510 model validates its use as a descriptive metric for assessing our intervention.

Importantly, BD treatment failed to improve cognition in mice that were fed a WD for the same 20-week duration. The WD we used causes insulin resistance, obesity^66^, and cognitive impairment^67^ in mice, which is consistent with cardiometabolic dysfunction. We found that BD-treated mice consuming WD did not achieve significant elevations in circulating βOHB and showed a lower relative abundance of L-βOHB in circulation when compared to mice on normal chow. Blunted BD-induced ketosis suggests that cardiometabolic dysfunction may diminish the neuroprotective or cognitive benefits of ketone bodies. Intriguingly, the directionality of correlations between Barnes Maze AUC and fasting blood glucose or fasting βOHB in WD-fed mice shows that metabolic inflexibility was surprisingly linked to beneficial cognitive outcomes (**Suppl Fig 11, Suppl Tables 5-6**), an outcome that was reversed in normal chow-fed mice. Despite these observations, it remains unknown if and how substrate preferences compensate for the setting of cardiometabolic dysfunction.

Pathological states, like those resulting from WD, may suppress classical ketogenesis due to sustained elevations in insulin and subsequent reductions in peripheral ketolysis from mitochondrial dysfunction^15^. This could limit BD metabolism or limit delivery of ketone bodies to the brain. Based on our results, we conclude that metabolic inflexibility associated with WD causes variations in absorption and disposal of exogenously delivered ketone bodies and negatively influences the response to these ketones, similar to conclusions recently reported^68^. Furthermore, the finding that BD treatment did not significantly influence brain mass or fasting blood glucose values (except in tauopathy females) supports that diet-induced metabolic dysfunction likely impairs the BD-induced effects in brain glucose utilization and lipid architecture that we associated with cognitive benefits in normal chow-fed mice.

We also tested the metabolic and cognitive effects of intermittently augmenting endogenous ketogenesis in tauopathy mice (**Fig 8**). Weekly cycling between WD and a KD produced predictable elevations in circulating ketone bodies, and unlike BD-treatment, increased only the D-βOHB in circulation. Cognitive function was unimproved, underscoring that the duration and timing of ketosis, as well as underlying metabolic health, may dictate functional outcomes associated with ketogenic interventions. Similar to the WD+BD mice, negative correlations between Barnes Maze AUC and fasting blood glucose were observed for both male and female mice (**Suppl Fig 11**) cycling WD with KD. However, in female tauopathy mice, but not males, there was a salutary effect of fasting βOHB on cognitive outcomes, which was absent in the WD+BD mice. These data imply that promoting physiological ketosis from either exogenous or endogenous sources does not produce a consistent cognitive phenotype in tauopathy mice with cardiometabolic dysfunction. Benefits may depend on the ability to produce and maintain consistent ketone body levels that allow brain uptake and subsequent metabolic changes supporting cognition. In WD-fed mice, we did not observe any cognitive benefits from exogenous or endogenous ketogenic interventions, which underscores the importance of systemic metabolic health in determining responsiveness to therapeutic, ketogenic strategies for combatting neurodegenerative diseases.

## 5. Limitations

Several limitations should be considered when interpreting the findings of this study. First, the BD intervention we used produced a mixture of D- and L-βOHB, making it difficult to associate the outcomes we measured with the canonical oxidative fate of D-βOHB or roles of L-βOHB. Each enantiomer may exert its own influence on brain metabolism and behavioral outcomes, where future work may consider isolating the effects of each enantiomer. Second, since this tauopathy mouse model produces hyperphosphorylated tau protein and neurofibrillary tangles in the forebrain region, we performed our metabolomics and lipidomics studies on freeze-clamped sections from the general forebrain region. While this produced a heterogeneous mixture of tissue and cell types from the brain, future work may benefit from isolating cell types that have differential effects from BD treatment. For example, early reports are now attempting to model metabolic contributions of neurons, astrocytes, microglia, and oligodendrocytes in neurodegenerative conditions^65,69^. Additionally, while we only reported fractional enrichment from our ITUM analyses, we did not measure pool totals or metabolic flux, which could yield greater insight into overall metabolic activity. Third, we chose the Barnes Maze as the primary cognitive outcome in this study. This is a hippocampus-dependent, spatial learning test performed over 5 days. Future work may consider incorporating additional behavioral studies to illuminate BD effects on other aspects relevant to Tau pathology, such as anxiety or sociability^70^. The tauopathy mice we used are hyperactive^71^, which may confound findings and support a rationale for additional behavioral analyses. Finally, we used a high-fat Western diet to stimulate a phenotype of general cardiometabolic dysfunction. There are other strategies to identify how specific contributors to cardiometabolic dysfunction, such as insulin resistance or cardiovascular disease, affect cognitive outcomes and are influenced by BD treatment. Understanding the unique contributions of each factor to metabolic diseases could identify conditions where ketogenic interventions are reliably beneficial to support brain metabolism and cognitive resilience in neurodegenerative diseases.

## Acknowledgements

We thank Erin Lind (UMN) for scientific insight and technical assistance, and we thank Russel H. Swerdlow (KUMC) for helpful conversations with study design. This work was supported by the following: National Institute of Health HL144472 (KF), DK140753 (AH), DK091538 (PAC), HL166142 (EDQ), DK136772 (CCH), and AG069781 (PAC and JPT). PAC has served as an external consultant for Selah Therapeutics, Pfizer Inc., Abbott Laboratories, and Janssen Research and Development. The other authors declare no competing interests. BioRender was used in creating figures and artificial intelligence was used to assist code generation for lipidomics analyses.

## Declaration of generative AI and AI-assisted technologies in the manuscript preparation process

During the preparation of this work the authors used generative AI in order to produce code for lipidomics analysis and generation of figures. After using this tool/service, the authors reviewed and edited the content as needed and take full responsibility for the content of the published article.

## Supplemental Methods

### Enantioseparation and Quantification of D/L-βOHB

Extracted serum samples were dried in a speed-vac and derivatization was adapted from previous reference^20,21^. Briefly, freshly prepared 5 mM (S)-(+)-1-(2-Pyrrolidinylmethyl)-pyrrolidine (PMP), 10 mM triphenylphosphine (TPP), and 10 mM 2,2′- dipyridyl disulfide (DPS) were dissolved separately in ACN and equal volumes were added to the dried samples, vortexed, and incubated for 90 min at room temperature. Samples were dried again in a speedvac and then resuspended in 40 μL of 2% Methanol, 98% H_2_O, and 0.0125% acetic acid. Samples were vortexed, sonicated for 5 min, centrifuged, and supernatant injected into LC-MS/MS system using positive ionization mode. Standards of D- or L-βOHB were also dried, derivatized, and injected alongside samples to confirm enantiomeric identity.

### Barnes Maze Testing

Cognitive function was assessed using a Barnes Maze protocol as previously described^22,23^. The Barnes Maze test assesses spatial learning and memory in rodents by placing mice on an elevated, circular platform (36-inch high, 36-inch diameter) with 20 x 2-inch holes. One of the holes leads to an escape box. We used a maze with a tan background since rTg4510 mice present with white, agouti, or black fur. Testing occurred in a brightly lit room with a 300-lux overhead light to motivate mice to find the dark escape box. Four contextual markers were positioned around the maze. Mice were first habituated to the maze platform and guided to the escape box. For each trial, a mouse started at the center of the platform. Between trials, the maze was cleaned with 70% ethanol to minimize scent trails. Then, from days 1 to 4, mice completed four acquisition trials daily, with a 15-minute inter-trial interval. Trials ended when the mouse found the escape box or after 3 minutes, whichever occurred sooner. Primary latency to escape hole entry was recorded for each trial. If the mouse did not find the escape hole within the 3-minute trial, the mouse was gently guided to the escape box and allowed 30 seconds rest inside the escape box. On day 5, mice were subjected to a 90-second probe trial with the escape box removed. We recorded primary errors, primary latency to escape hole entry, and percentage of time spent in the goal quadrant derived from video acquisition using AnyMaze Software (Stoelting Co.; Wood Dale, IL) provided by the UMN Behavioral Phenotyping Core.

### Isotope Tracing Untargeted Metabolomics (ITUM)

Mice were fasted for 18h prior to isotope tracing experiments. Fifteen minutes after an intraperitoneal injection of 5 µmol/g BW [U-^13^C_6_]glucose, mice were euthanized with pentobarbital and decapitated for rapid collection of the brain frontal cortex, which was immediately freeze-clamped. Blood samples were collected at t=0 min and t=15 min to quantify plasma glucose concentration. ITUM workflow proceeded as previously described^26^. Briefly, 20 mg brain samples were homogenized in (2:2:1) Methanol:ACN:H_2_O extraction buffer, and then metabolites extracted from 2 mg tissue homogenate in 1 mL extraction buffer by three cycles of vortexing, freeze-thawing, and water bath sonication. Samples were incubated at -20 °C for 1 hour, followed by 10 min centrifugation. Exactly 800 µL supernatant was transferred to new tubes, and the solvent was evaporated in a speed-vac. Finally, the dried metabolite pellet was reconstituted in 100 µL (1:1) ACN:H_2_O with the aid of vortexing and sonication, then incubated at 4 °C for 1 h before spinning samples and loading in vials for LC/MS analysis. To aid metabolite identification, an in-house standard mixture of 100 metabolites and a pooled sample mix were analyzed by data-dependent acquisition (DDA). Identifications were based on retention time compared to authentic standard, m/z that matches theoretical m/z predicted from molecular formula (±10 ppm), and MS2 fragmentation patter matched to online databases (e.g. mzCloud). Electrospray ionization drift was monitored by injecting a pooled quality control (QC) sample at regular intervals.

Extracted metabolites were separated using the Thermo Fisher Scientific Vanquish UHPLC system, followed by detection using high mass resolution mass spectrometry on the Thermo Fisher Scientific QExactive plus hybrid quadrupole-orbitrap mass spectrometer, fitted with heated electrospray ionization source. Atlantis Premier BEH Z-HILIC column (1.7 µm particle size, 2.1 mm x 100 mm) was used to separate metabolites using hydrophilic interaction liquid chromatography (HILIC) mode using following mobile phases: MPA = 15 mM NH_4_HCO_3_ in 100% H_2_O, pH = 9.0; MPB = 15 mM NH_4_HCO_3_ in 90% ACN, pH = 9.0. Total run time was 10 min, flow rate was 0.5 mL/min for the first 6 minutes and then increased to 1 mL/min for the remainder of the run. Column chamber was maintained at 30 °C. Mobile phase gradient was as follows: 0 to 0.1 min, 90% MPB; 0.1 to 5 min, 90 → 65% MPB; 5 to 6 min, 65% MPB; 6 to 6.5 min, 65 → 90% MPB; 6.5 to 10 min, 90% MPB. For analyses, both blanks and QC samples were injected periodically throughout the run. Blank samples containing only ACN:H_2_O (1:1)^26^. Thermo Compound Discoverer 3.3 was used for the analysis. To confirm fractional enrichment of metabolites with low enrichment (<1%), manual integration using QuanBrowser on Thermo Fisher Xcalibur software was performed.

### Shotgun lipidomics

Lipidomics extractions and analyses were performed as previously described^27^. Briefly, brains were freeze-clamped, weighed (∼20 mg) and homogenized using zirconium oxide beads and an automated tissue homogenizer (Omni International, Bead Ruptor 12, 19-050A) using 1.0 mL ice-cold, 0.1x PBS lacking both MgCl_2_ and CaCl_2_. Total protein was determined using BCA assay, then internal standards were added based on the sample protein amount. Bulk lipid extraction was completed using a modified Bligh-Dyer method, as described previously^28^. Finally, samples were directly infused into a TSQ Vantage triple quadrupole mass spectrometer (Thermo Fisher Scientific, San Jose, CA) equipped with an automated Triversa Nanomate nanospray device (Advion Biosciences, Ithaca, NY) for multidimensional MS-based shotgun lipidomics.

---

**Supplemental Figure 1.**
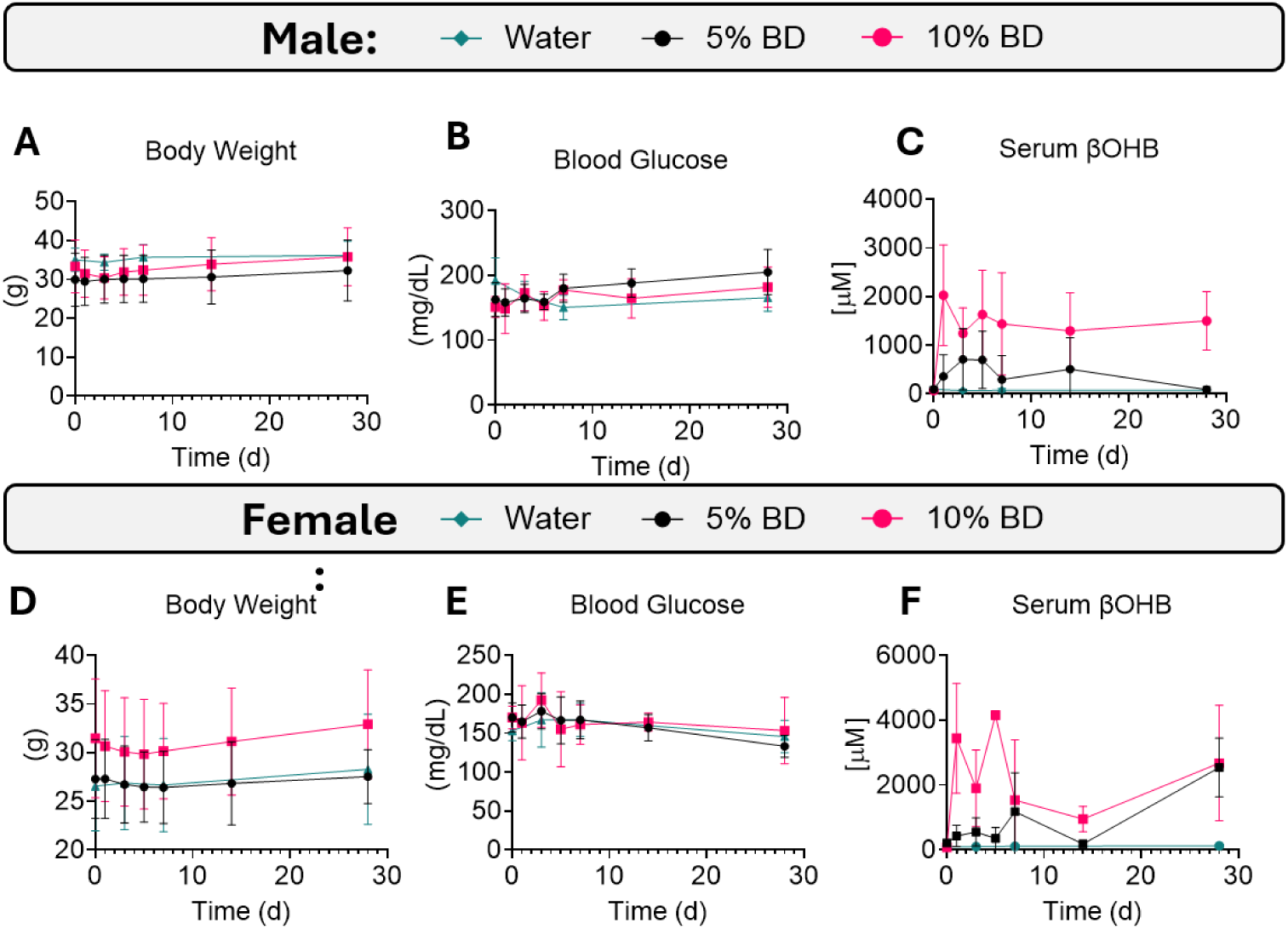
Kinetics of 1,3-butanediol (BD) in male and female mice. Male and female mice were treated acutely with 5% or 10% BD in drinking water for 28 days. In male mice given the BD treatment, water controls showed no effects on body weight (**A**), blood glucose (**B**), or serum βOHB (**C**). Similarly, in female mice given the BD treatment, water controls showed no effects on body weight (**D**), blood glucose (**E**), or serum βOHB (**F**) compared to baseline measures. Data for this figure were re-plotted from figure 1 to show comparison to water controls (*n*=4-5 water controls per panel). BD = 1,3-butanediol, βOHB= β-hydroxybutyrate.

**Supplemental Figure 2.**
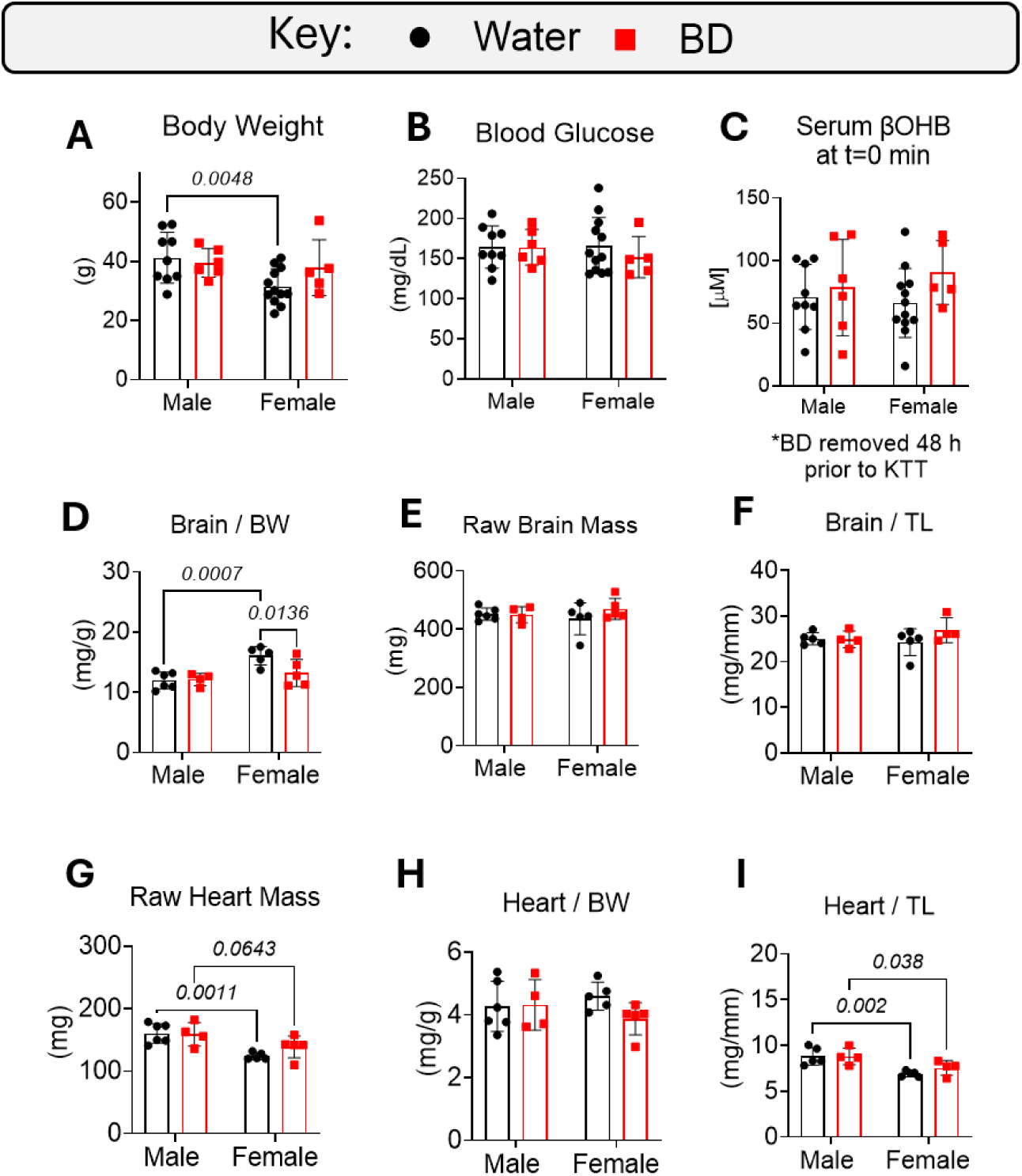
Chronic BD treatment has minimal influence on body weight and tissue mass in healthy mice. Male and female mice were treated with 10% BD in drinking water for 20 weeks and then subjected to a ketone tolerance test (KTT). Effects of treatment on body weight (**A**) and non-fasted blood glucose (**B**) in BD-treated mice compared to normal water controls. After 48 h removal of BD water, serum βOHB was unchanged between groups (**C**). BD treatment had no effect on raw or normalized brain mass (**D-F**), and there were no effects of BD treatment on raw or normalized heart (**G-I**). Data are expressed as mean ± SD, *n*=4-5/group. P-values indicated where p<0.10 following two-way ANOVA and Tukey’s post-hoc test. BD = 1,3-butanediol, βOHB= β-hydroxybutyrate, TL = tibia length.

**Supplemental Figure 3.**
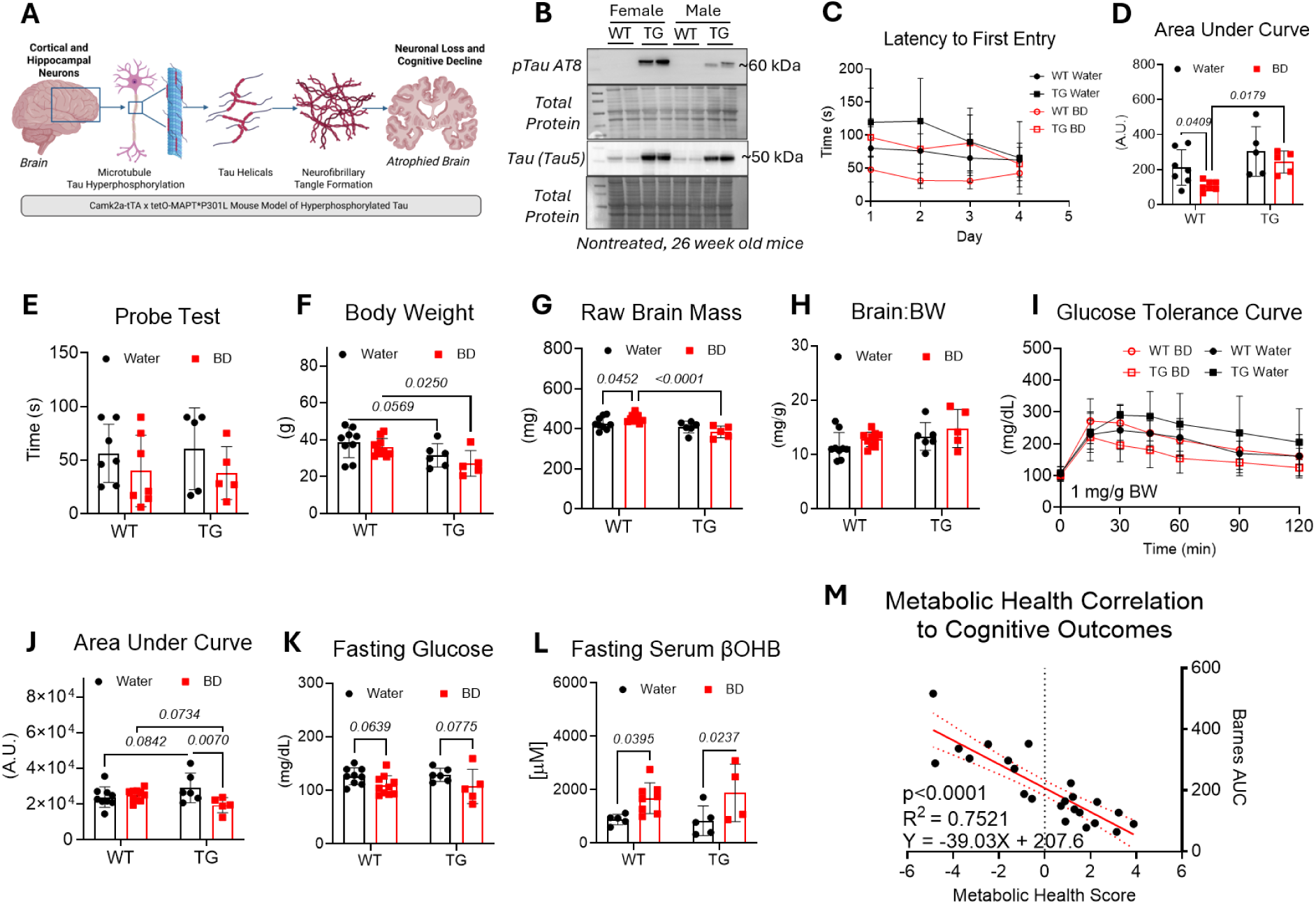
Twenty-week BD treatment may improve cognitive outcomes in non-tauopathy, but not tauopathy, male mice. Schematic of tauopathy mouse model used in this study (**A**) with representative immunoblot of phosphorylated human Tau (AT8) and total Tau (Tau5) in nontreated, 26-week aged male and female tauopathy mice and non-transgenic control mice (**B**, *n*=2/group). Male tauopathy mice and non-transgenic littermate controls were also provided with 10% BD in drinking water or normal drinking water for 20 weeks, and cognitive outcomes were assessed. Mice were subjected to Barnes Maze behavioral testing following 20-week treatment. Latency to escape hole entry during memory acquisition phase, days 1-4 (**C**, *n*=5-7/group) and corresponding area under the curve for the trials (**D**). Memory recall probe test, day 5 (**E**, *n*=5-7/group) in male mice. Effects of genotype and 20 weeks BD treatment on body weight (**F**), raw brain mass (**G**), and brain mass normalized to body weight (**H**). Systemic glucose tolerance test (**I**) performed at 25 weeks of age with corresponding area under the curve (**J**) for the 120-min test. Following the 20-week intervention, mice were fasted for 18 h and then euthanized. Effects of 20-week BD treatment on fasting blood glucose (**K**) and serum βOHB while consuming BD-treated water (**L**). Pearson’s correlations were calculated for variables, and a metabolic health score (MHS) was constructed to predict cognitive outcomes in male mice, as shown in (**M**). Data are presented as mean ± SD, *n*=5-10/group unless otherwise indicated. P-values indicated when p<0.10 following two-way ANOVA and Tukey’s post-hoc test. BD = 1,3-butanediol, βOHB= β-hydroxybutyrate, ITUM = isotope tracing untargeted metabolomics, A.U. = arbitrary units, BW = body weight, WT = non-tauopathy control mice, TG = tauopathy mice, MHS = metabolic health score

**Supplemental Figure 4.**
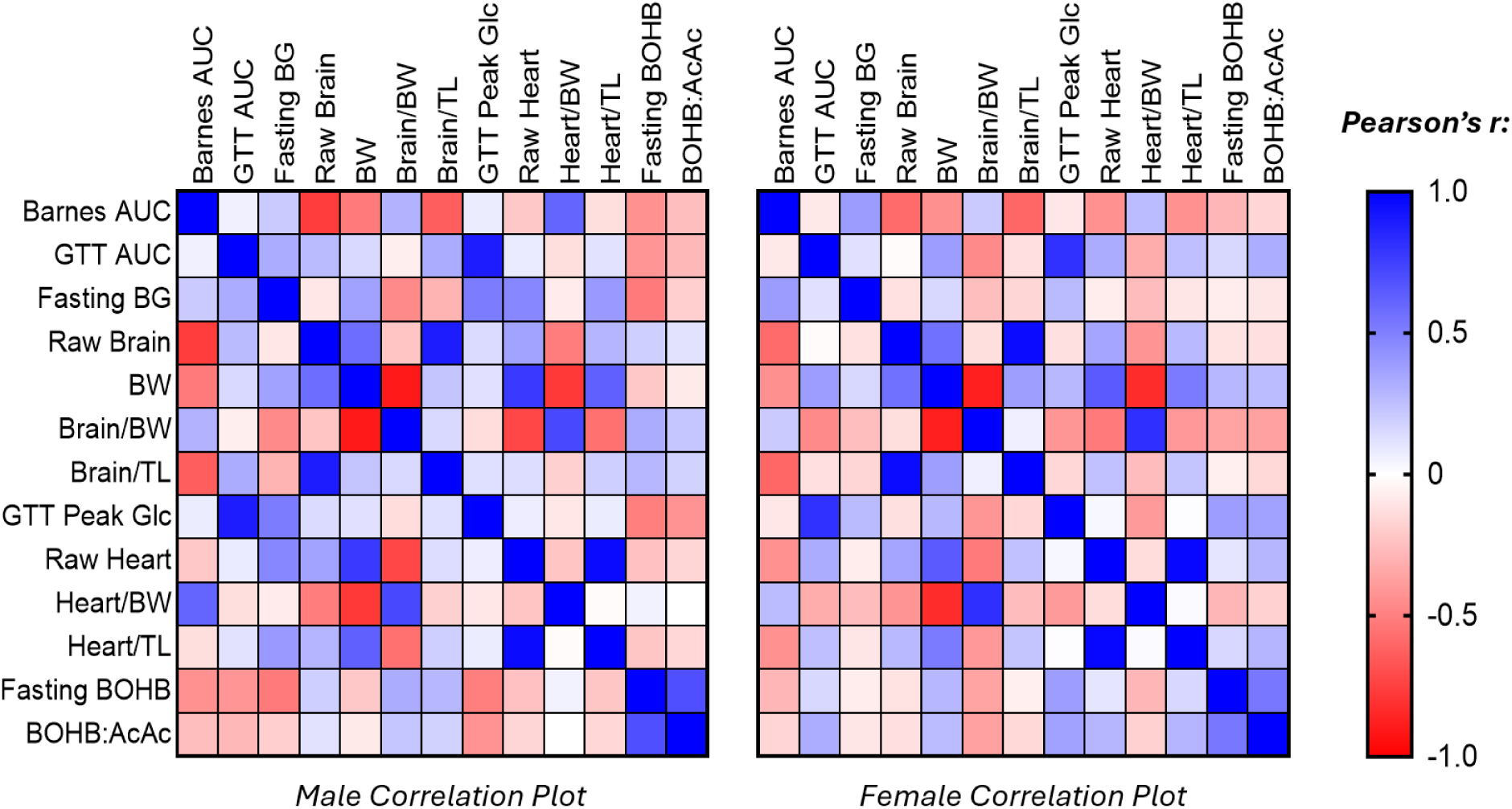
Correlation heatmaps for male and female mice consuming normal chow with or without BD for 20 weeks. Heatmaps showing correlations among variables in male and female mice, with or without 10% BD in the drinking water for 20-week treatment. Data represented as Pearson’s r, *n*=23-29/group. AUC = area under curve, βOHB= β-hydroxybutyrate, GTT = glucose tolerance test, BG = blood glucose, BW = body weight, Glc = glucose, TL = tibia length for normalization.

**Supplemental Figure 5.**
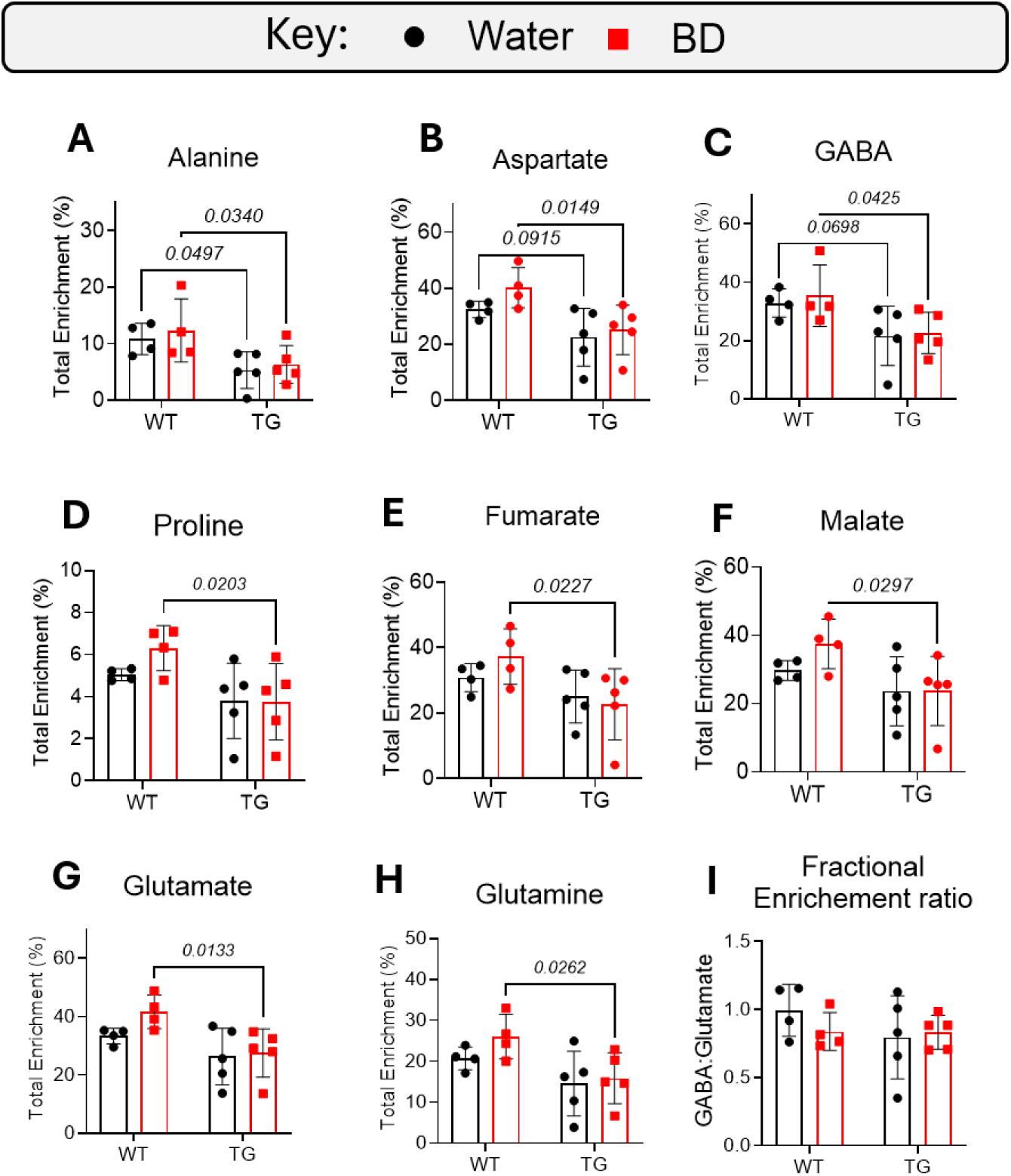
Twenty-week BD treatment influences acute glucose utilization into anabolic pathways in the forebrain of male mice. Following 20-week BD treatment, male mice underwent 15-min ITUM with [U-^13^C_6_]-glucose delivered via I.P. injection. Total ^13^C enrichment from glucose-derived carbon was measured in metabolites from freeze-clamped, forebrain sections of male mice. Fractional enrichment of ^13^C was traced in Alanine (**A**), Aspartate (**B**), GABA (**C**), Proline (**D**), Fumarate (**E**), Malate (**F**), Glutamate (**G**), and Glutamine (**H**) in forebrain regions. Ratio of glucose-derived enrichment in GABA: Glutamate shown in (**I**). Data are expressed as mean ± SD, *n*=4-5/g. P-values indicated when p<0.10 following two-way ANOVA and Tukey’s post-hoc test. GABA = gamma-aminobutyric acid, WT = non-tauopathy control mice, TG = tauopathy mice

**Supplemental Figure 6.**
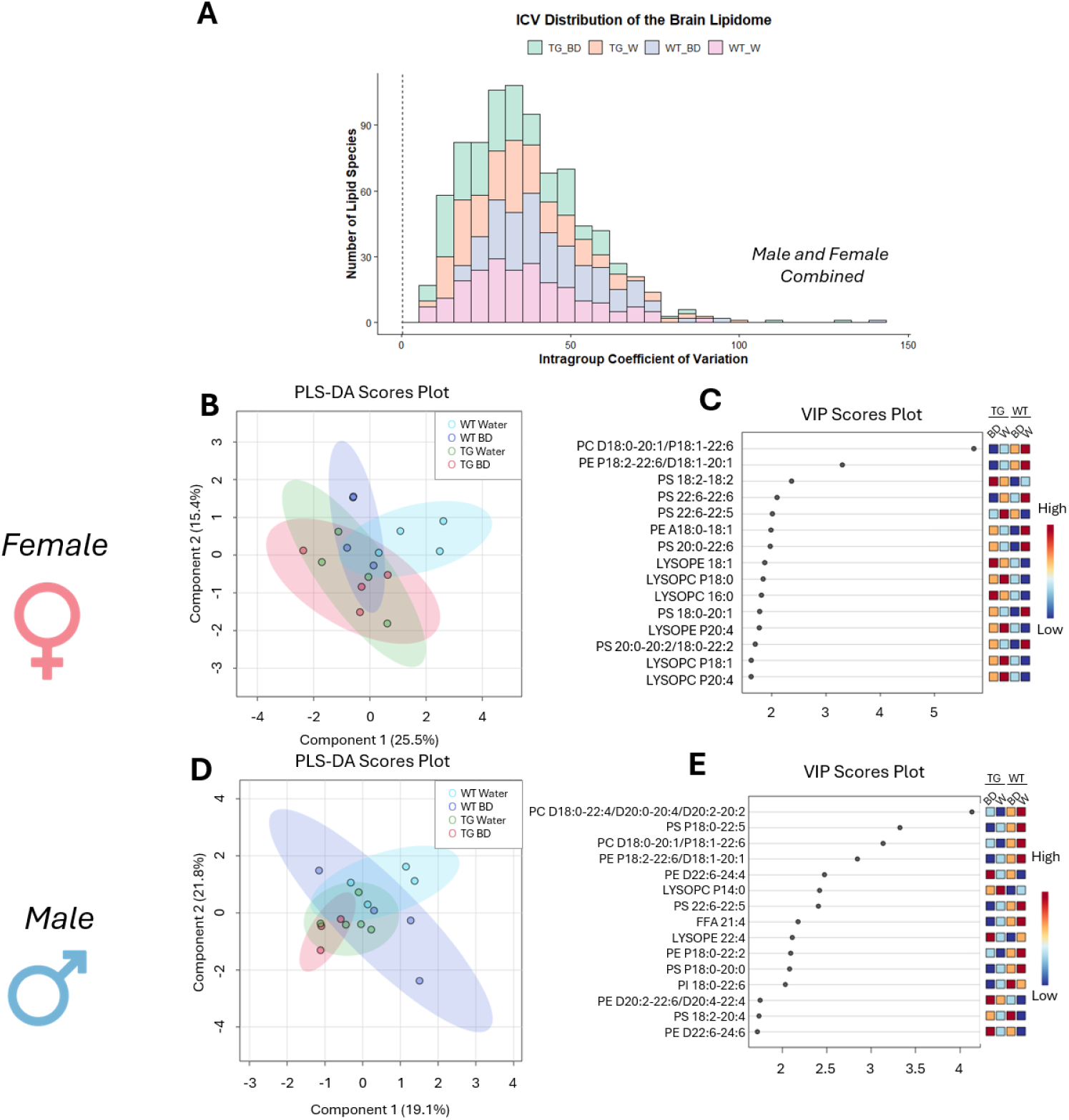
BD treatment modestly influences the brain lipidome of mice in a treatment and genotype-specific manner. Following 20-week BD treatment, a subset of male and female mouse brains was freeze-clamped, and forebrain sections were subjected to lipidomic analyses. Intragroup coefficient of variation for pooled male and female samples from each treatment group is shown in (**A**). Absolute concentrations of lipid species were analyzed using Metaboanalyst 6.0 software and were log-transformed to show group separation via partial least squares-discriminant analysis in female brains (PLS-DA, **B**), highlighting the corresponding variable importance plot (VIP) scores for the top 15 metabolites (**C**). A PLS-DA plot was also constructed for male brains (**D**) with corresponding VIP scores for the top 15 metabolites (**E**). Data are expressed as *n*=3-5/group. WT = non-tauopathy control mice, TG = tauopathy mice, PC = phosphatidylcholine, PE = phosphatidylethanolamine, PA = phosphatidic acid, PI = phosphatidylinositol, PS = phosphatidylserine, SM = sphingomyelin, CBS = cerebroside, FFA = free fatty-acid, LPC = lysoPC, LPE = lysoPE, Car = acylcarnitine, W = water treatment, BD = 1,3-butanediol treatment

**Supplemental Figure 7.**
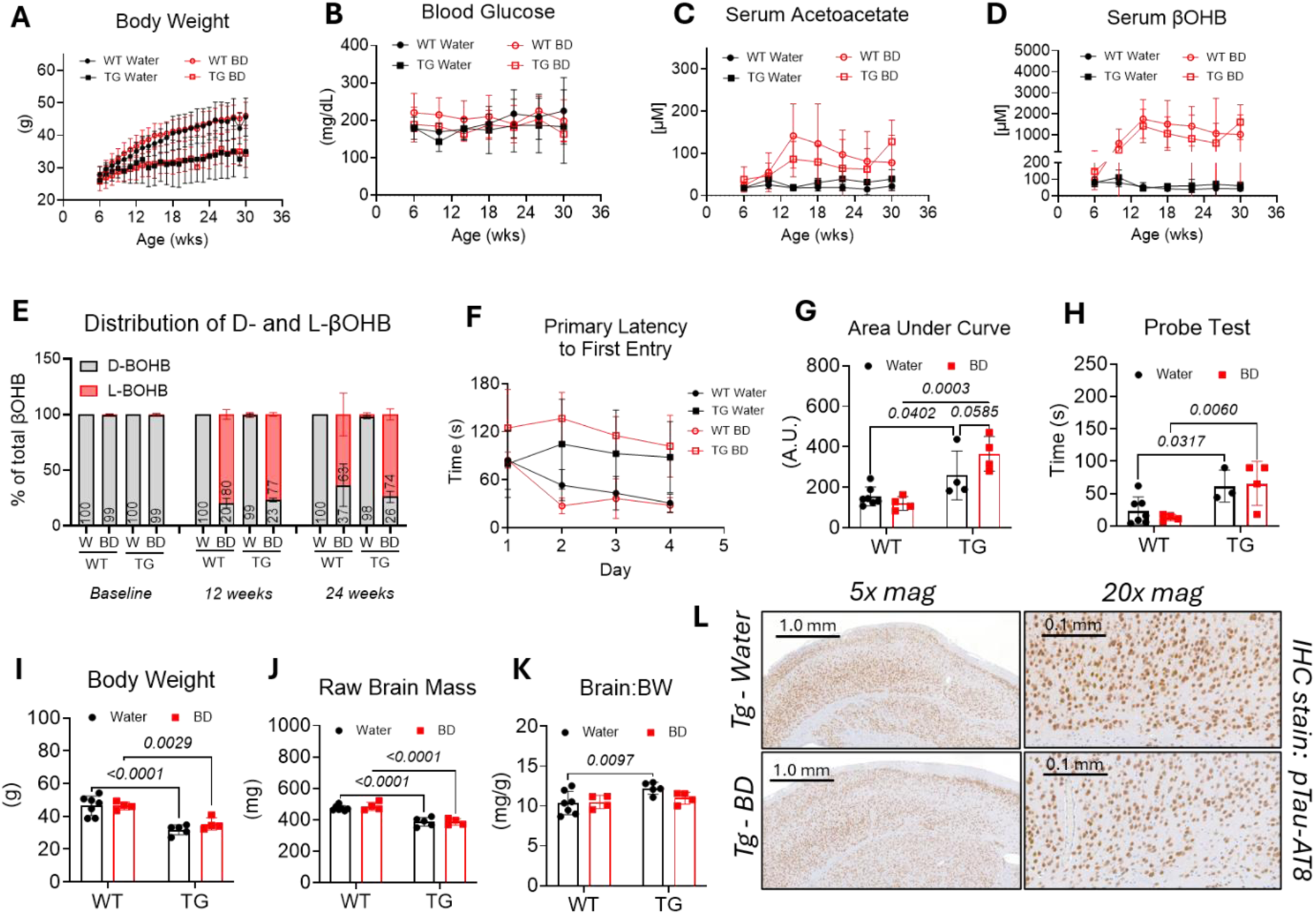
Thirty-week BD treatment has no significant effects on cognitive or brain outcomes in male tauopathy-transgenic mice. Male tauopathy mice and non-transgenic littermate controls were also provided with 10% BD in drinking water or normal drinking water for 30 weeks, and cognitive outcomes were assessed. Effects of the transgene and BD treatment on body mass (**A**), non-fasted blood glucose (**B**), serum Acetoacetate (**C**), and serum βOHB (**D**) over 30-week treatment. Total percentage of D- and L-βOHB in circulation at baseline (6 weeks of age), 12-week treatment (18 weeks of age), and 24-week treatment (30 weeks of age, **E**, *n*=3/group). Mice were subjected to the Barnes-Maze for cognitive testing. Latency to escape hole entry during memory acquisition phase, days 1-4 (**F**) and corresponding area under the curve for the trials (**G**). A memory probe test was conducted on day 5 and is shown in (**H**). Thirty-week BD treatment did not influence body mass (**I**), raw brain mass (**J**), or brain mass normalized to body weight (**K**) in transgenic or non-transgenic male mice at tissue harvest. Representative images from immunohistochemistry staining of paired helical filaments in the forebrain region (**L**) from tauopathy-transgenic mice exposed to BD treatment or normal drinking water. Data are expressed as mean ± SD, *n*=4-7/group unless otherwise indicated. P-values indicated where p<0.10 following two-way ANOVA and Tukey’s post-hoc test. BD = 1,3-butanediol, A.U. = arbitrary units, βOHB= β-hydroxybutyrate, mag = magnification, IHC = immunohistochemistry, WT = non-tauopathy control mice, TG = tauopathy mice

**Supplemental Figure 8.**
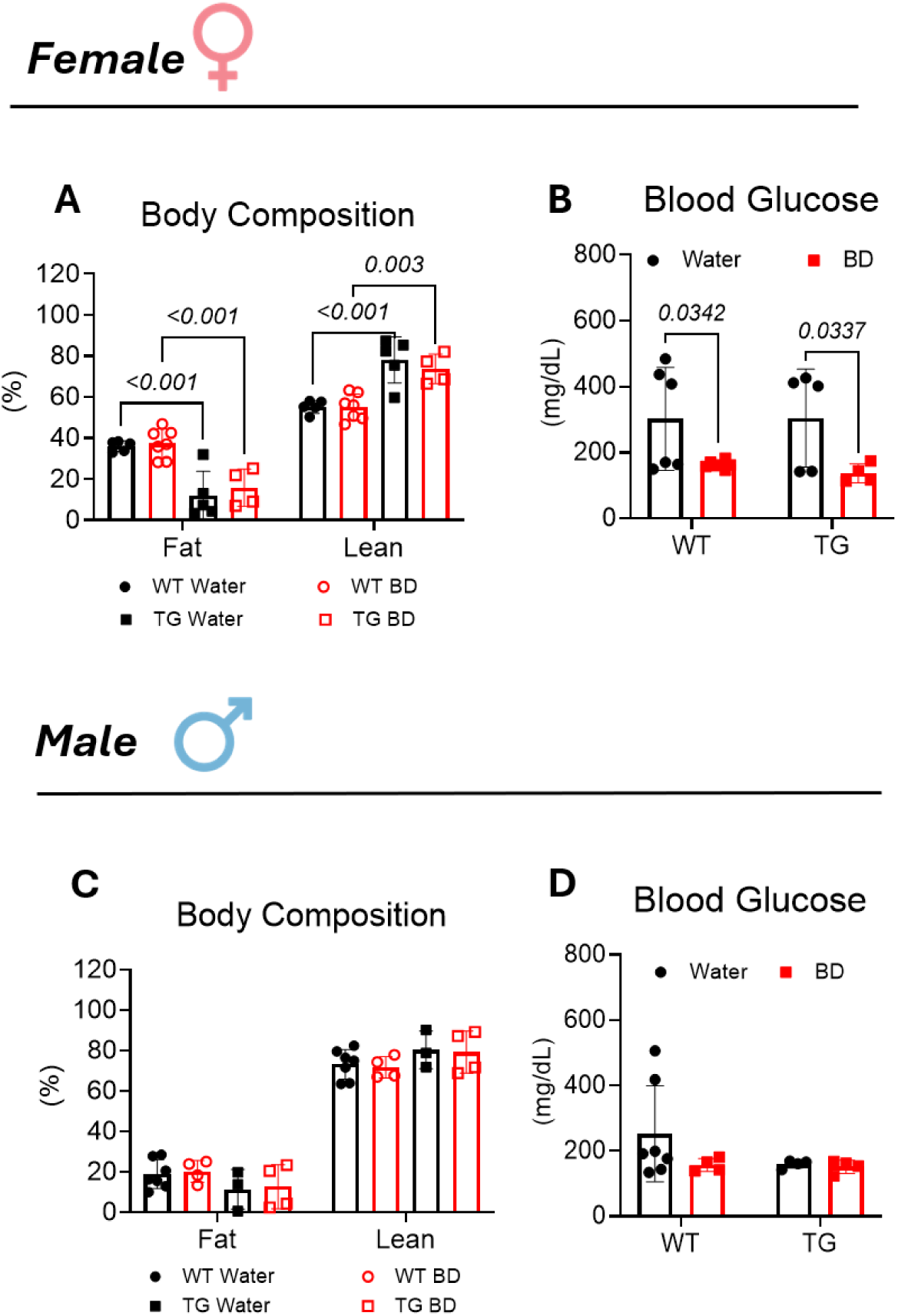
Sex-dependent effects of 30-week BD treatment on body composition and blood glucose. Following 30-week BD treatment in tauopathy and non-tauopathy mice, we measured body composition in female mice (**A**) and recorded the effects of chronic BD treatment on non-fasting blood glucose (**B**). We also measured body composition in male mice (**C**) and recorded effects of chronic BD treatment on non-fasting blood glucose (**D**). Data are expressed as mean ± SD, *n*=4-7/group. P-values indicated where p<0.10 following two-way ANOVA and Tukey’s post-hoc test. WT = non-tauopathy control mice, TG = tauopathy mice

**Supplemental Figure 9.**
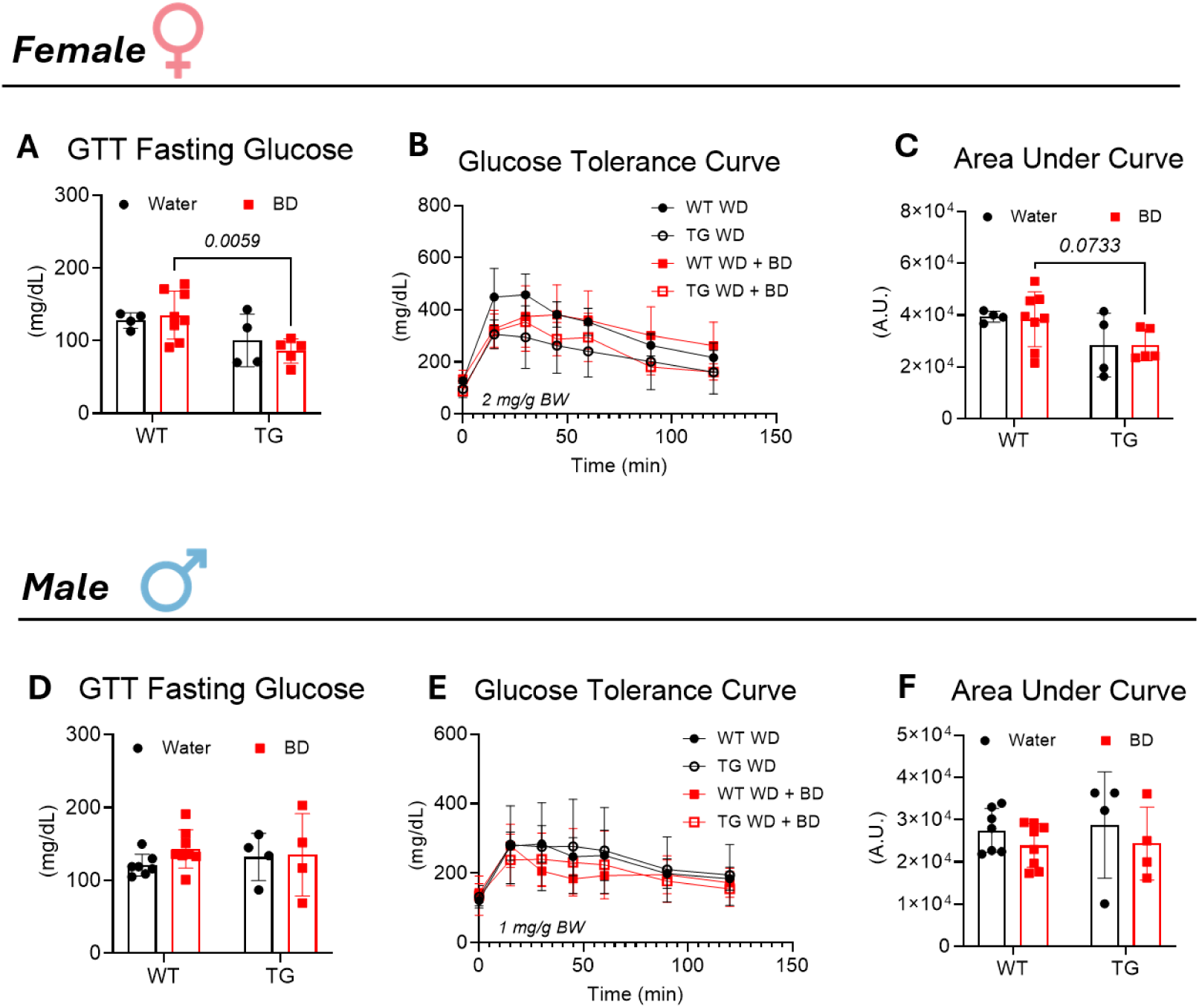
BD treatment does not influence systemic glucose tolerance in mice on a high-fat, Western diet (WD). Following a 20-week high-fat, Western diet either with or without BD treatment, mice were fasted for 18 h and then subjected to a glucose tolerance test (GTT) via I.P. injection of glucose (1 mg/g BW males; 2 mg/g BW females). Effects of BD treatment on female mouse fasted blood glucose values (**A**) or glucose tolerance curve (**B**) with corresponding area under the curve (**C**) when consuming the high-fat, Western diet. Effects of the BD treatment on male mouse fasted blood glucose values (**D**) or glucose tolerance curve (**E**) with corresponding area under the curve (**F**) when mice were consuming a high-fat, Western diet. Data are expressed as mean ± SD, *n*=4-8/group. P-values indicated where p<0.10 following two-way ANOVA with Tukey’s post-hoc test. WD = Western diet, BD = 1,3-butanediol, GTT = glucose tolerance test, WT = non-tauopathy control mice, TG = tauopathy mice

**Supplemental Figure 10.**
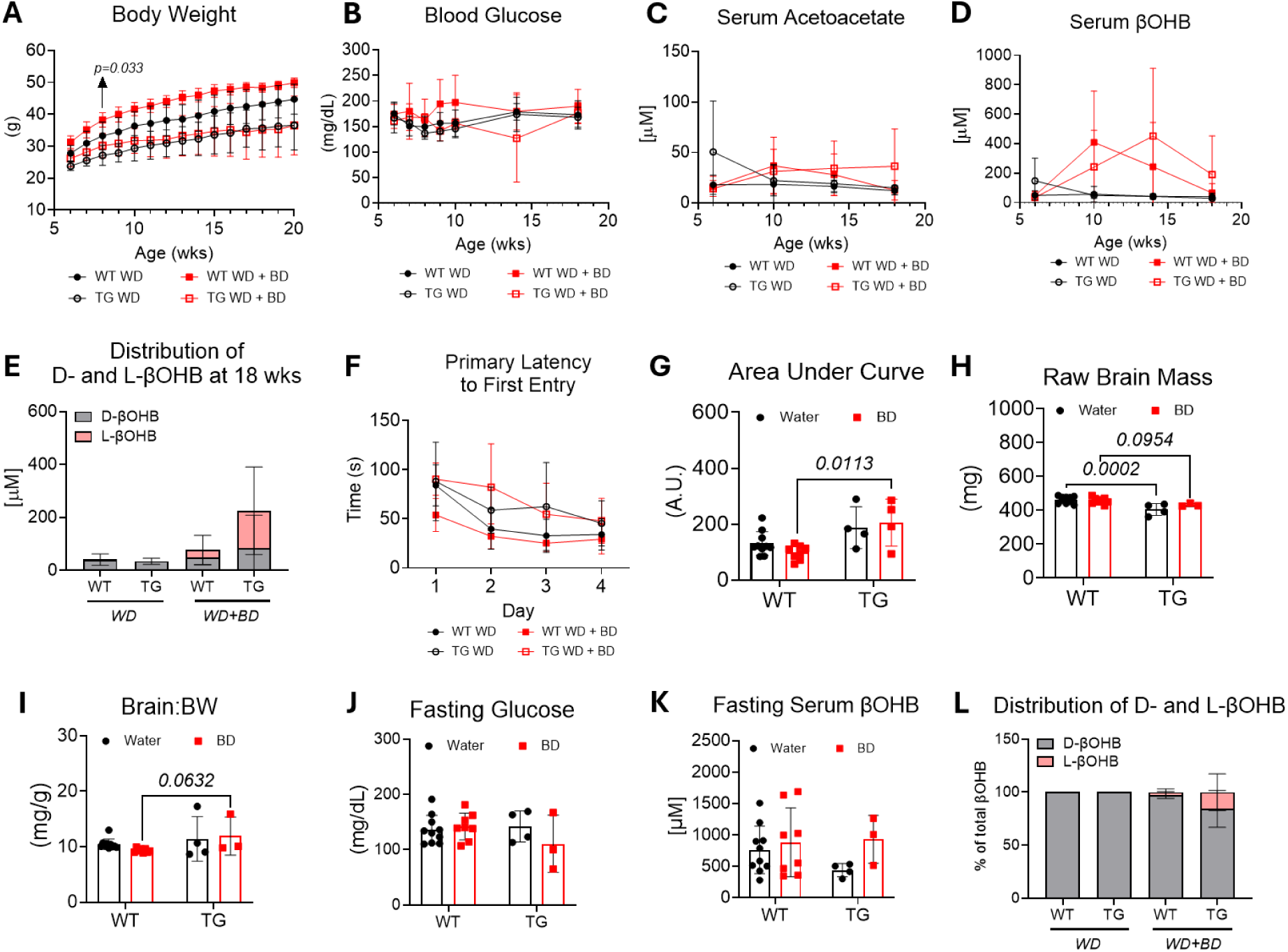
High-fat, Western diet abolishes cognitive benefits of BD treatment in male mice. Male tauopathy mice and non-transgenic littermate controls, were also subjected to a 20-week high-fat Western diet while consuming 10% BD or normal drinking water. Cognitive function was assessed using a Barnes Maze before tissues were harvested for analyses. Effects of high-fat, Western diet with or without BD treatment on body weight (**A**) and non-fasted blood glucose (**B**) throughout the study. BD treatment effects on serum acetoacetate (**C**), serum βOHB (**D**), as well as the concentration and distribution of D- and L-βOHB at study midpoint (12-week treatment, 18 weeks of age, **E**). Latency to escape hole entry during memory acquisition phase, days 1-4 of Barnes Maze (**F**) with corresponding area under curve (**G**). Following cognitive testing, mice were euthanized, and the raw brain mass (**H**) was plotted with brain mass normalized to body weight (**I**). Following an 18 h fast, effects of BD treatment on fasted blood glucose (**J**), fasted serum βOHB (**K**), and distribution of D- and L-βOHB (**L**). Data are expressed as mean ± SD, *n*=3-10/group. P-values indicated when p<0.10 following two-way ANOVA and Tukey’s post-hoc test. βOHB= β-hydroxybutyrate, WD = high-fat, Western diet, BD = 1,3-butanediol, A.U. = arbitrary units, WT = non-tauopathy control mice, TG = tauopathy mice.

**Supplemental Figure 11.**
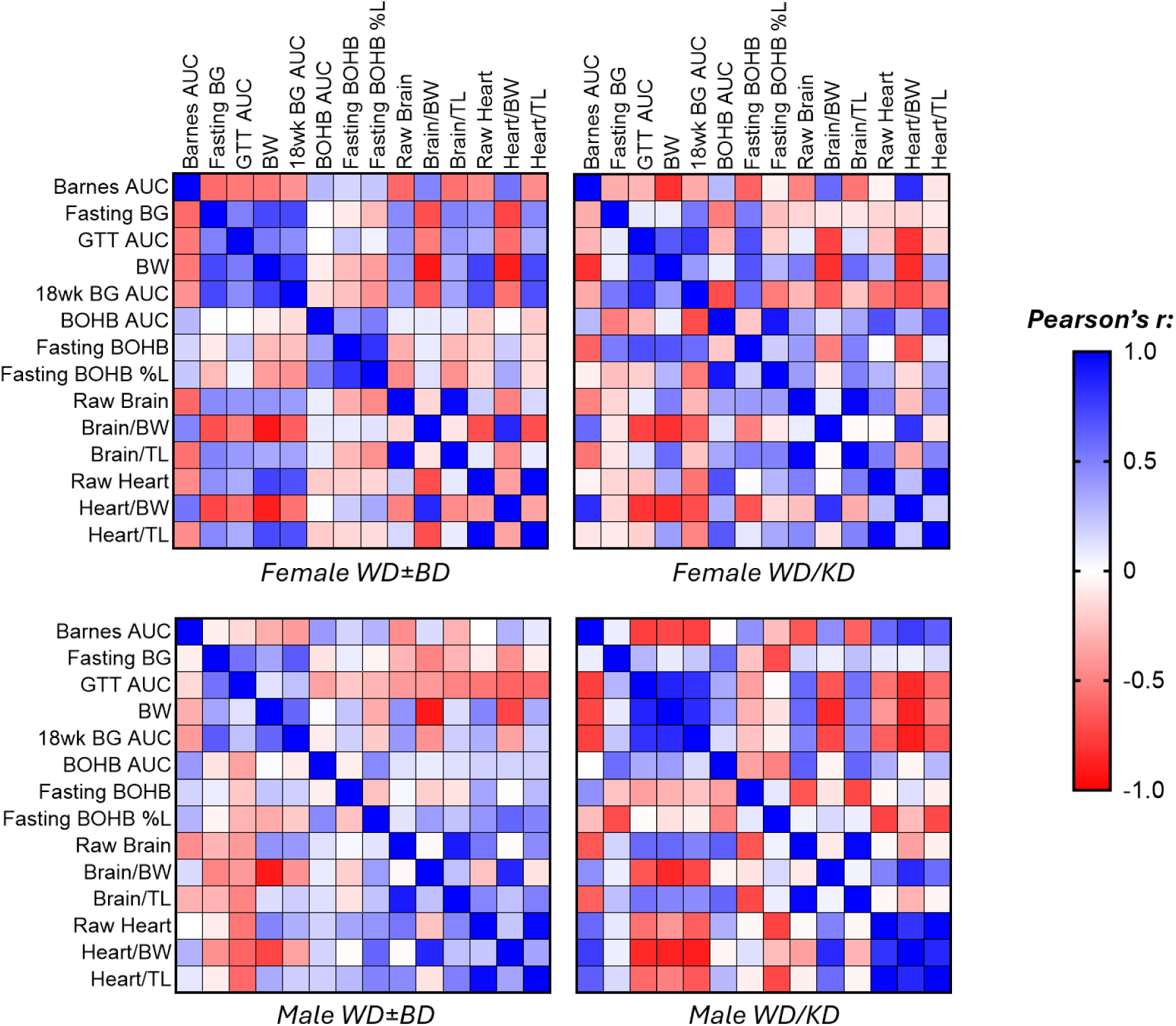
Correlation heatmap plots for Western diet-fed mice. Heatmaps showing Pearson’s r values for correlations in settings of Western diet (WD) with or without 10% BD in drinking water, and for weekly cycling of WD with a ketogenic diet (KD) for both male and female mice. Data are expressed as Pearson’s r, *n*=10-26/group. AUC = area under the curve, BG = blood glucose, GTT = glucose tolerance test, BW = body weight, TL = tibia length for normalization.

**Supplemental Figure 12.**
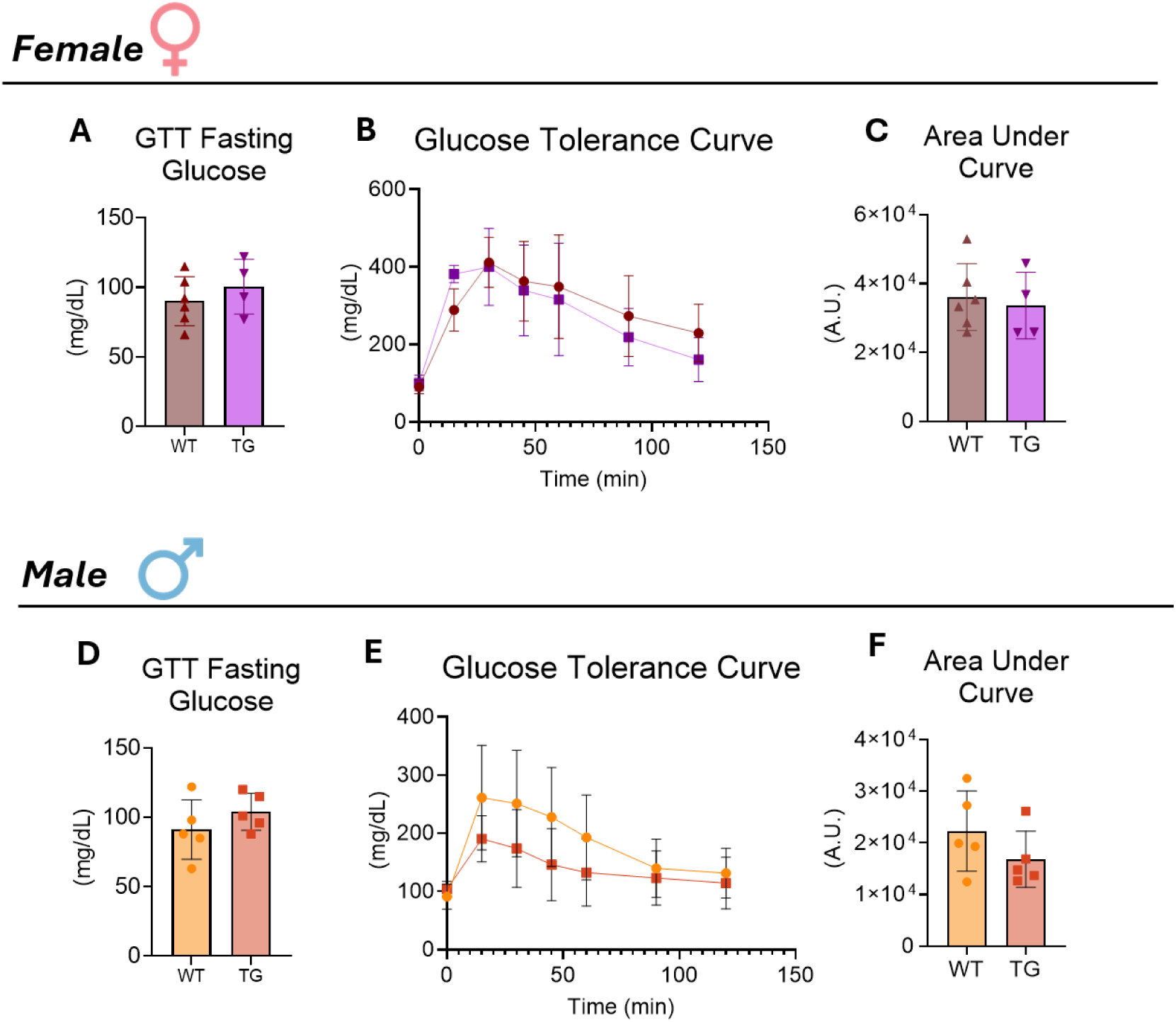
Cycling a high-fat, ketogenic diet with a high-fat, Western diet has minimal effects on glucose tolerance in tauopathy male and female mice. Male and female mice were fed a 20-week dietary strategy that cycled a high-fat, Western diet weekly with a high-fat, very low-carbohydrate ketogenic diet. During a ketogenic diet week near the end of the study, mice were fasted for 18 h and then subjected to a glucose tolerance test (GTT) via I.P. injection of glucose (1 mg/g BW males; 2 mg/g BW females). Fasted blood glucose values (**A**) and glucose tolerance curve (**B**) with corresponding area under the curve (**C**) were recorded for female transgenic and non-transgenic mice. Fasted blood glucose values (**D**) and glucose tolerance curve (**E**) with corresponding area under the curve (**F**) were also recorded for male transgenic and non-transgenic mice. Data are expressed as mean ± SD, *n*=4-6/group. GTT = glucose tolerance test, A.U. = arbitrary units, WT = non-tauopathy control mice, TG = tauopathy mice

**Supplemental Table 1:**
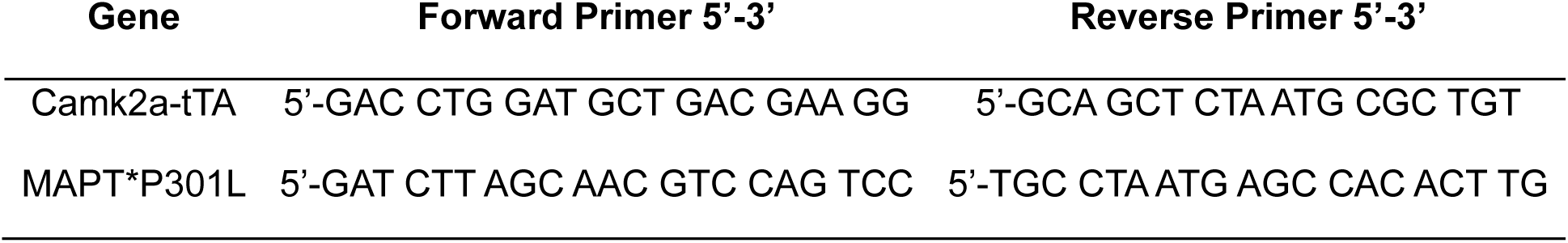
Genotyping primer sequences. Forward and reverse primer sequences (5’ – 3’) to confirm tauopathy transgenic mice. Tauopathy mice are positive for both Camk2a-tTA and MAPT*P301L transgenes.

**Supplemental Table 2:** Pearson’s correlation of metabolic variables with cognitive outcome in female mice. Pearson’s correlation coefficient, r, for various metabolic variables following a 20-week normal chow (NC) diet with or without 10% (v/v) 1,3-butanediol (BD) dissolved in drinking water. Positive values indicate positive correlation and negative values indicate negative correlation. *n*=29 females were used in correlation calculations. AUC = area under curve, GTT = glucose tolerance test, BG = blood glucose, BW = body weight, TL = tibia length, Glc = glucose, βOHB = total beta-hydroxybutyrate (both D and L enantiomers), AcAc = acetoacetate.

|  | Barnes AUC | GTT AUC | Fasting BG | Raw Brain | BW | Brain/BW | Brain/TL | GTT Peak Glc | Raw Heart | Heart/BW | Heart/TL | Fasting BOHB | BOHB/AcAc |
| --- | --- | --- | --- | --- | --- | --- | --- | --- | --- | --- | --- | --- | --- |
| **Barnes AUC** | 1.000 | -0.093 | 0.387 | -0.586 | -0.437 | 0.204 | -0.597 | -0.098 | -0.432 | 0.265 | -0.435 | -0.288 | -0.168 |
| **GTT AUC** | -0.093 | 1.000 | 0.116 | -0.017 | 0.383 | -0.459 | -0.128 | 0.810 | 0.323 | -0.329 | 0.250 | 0.150 | 0.319 |
| **Fasting BG** | 0.387 | 0.116 | 1.000 | -0.121 | 0.151 | -0.258 | -0.163 | 0.270 | -0.071 | -0.264 | -0.099 | -0.075 | -0.099 |
| **Raw Brain** | -0.586 | -0.017 | -0.121 | 1.000 | 0.556 | -0.132 | 0.954 | -0.125 | 0.350 | -0.423 | 0.271 | -0.114 | -0.129 |
| **BW** | -0.437 | 0.383 | 0.151 | 0.556 | 1.000 | -0.880 | 0.379 | 0.278 | 0.643 | -0.828 | 0.515 | 0.282 | 0.266 |
| **Brain/BW** | 0.204 | -0.459 | -0.258 | -0.132 | -0.880 | 1.000 | 0.061 | -0.413 | -0.524 | 0.813 | -0.402 | -0.362 | -0.369 |
| **Brain/TL** | -0.597 | -0.128 | -0.163 | 0.954 | 0.379 | 0.061 | 1.000 | -0.158 | 0.240 | -0.264 | 0.229 | -0.063 | -0.154 |
| **GTT Peak Glc** | -0.098 | 0.810 | 0.270 | -0.125 | 0.278 | -0.413 | -0.158 | 1.000 | 0.031 | -0.398 | 0.007 | 0.383 | 0.367 |
| **Raw Heart** | -0.432 | 0.323 | -0.071 | 0.350 | 0.643 | -0.524 | 0.240 | 0.031 | 1.000 | -0.134 | 0.971 | 0.099 | 0.284 |
| **Heart/BW** | 0.265 | -0.329 | -0.264 | -0.423 | -0.828 | 0.813 | -0.264 | -0.398 | -0.134 | 1.000 | 0.015 | -0.288 | -0.183 |
| **Heart/TL** | -0.435 | 0.250 | -0.099 | 0.271 | 0.515 | -0.402 | 0.229 | 0.007 | 0.971 | 0.015 | 1.000 | 0.155 | 0.291 |
| **Fasting BOHB** | -0.288 | 0.150 | -0.075 | -0.114 | 0.282 | -0.362 | -0.063 | 0.383 | 0.099 | -0.288 | 0.155 | 1.000 | 0.532 |
| **BOHB/AcAc** | -0.168 | 0.319 | -0.099 | -0.129 | 0.266 | -0.369 | -0.154 | 0.367 | 0.284 | -0.183 | 0.291 | 0.532 | 1.000 |

**Supplemental Table 3:** Pearson’s correlation of metabolic variables with cognitive outcome in male mice. Pearson’s correlation coefficient, r, for various metabolic variables following a 20-week normal chow (NC) diet with or without 10% (v/v) 1,3-butanediol (BD) dissolved in drinking water. Positive values indicate positive correlation and negative values indicate negative correlation. *n*=23 males were used in correlation calculations. AUC = area under curve, GTT = glucose tolerance test, BG = blood glucose, BW = body weight, TL = tibia length, Glc = glucose, βOHB = total beta-hydroxybutyrate (both D and L enantiomers), AcAc = acetoacetate.

|  | Barnes AUC | GTT AUC | Fasting BG | Raw Brain | BW | Brain/BW | Brain/TL | GTT Peak Glc | Raw Heart | Heart/BW | Heart/TL | Fasting BOHB | BOHB/AcAc |
| --- | --- | --- | --- | --- | --- | --- | --- | --- | --- | --- | --- | --- | --- |
| **Barnes AUC** | 1.000 | 0.056 | 0.204 | -0.764 | -0.525 | 0.298 | -0.632 | 0.074 | -0.218 | 0.608 | -0.130 | -0.432 | -0.259 |
| **GTT AUC** | 0.056 | 1.000 | 0.327 | 0.270 | 0.145 | -0.068 | 0.323 | 0.888 | 0.075 | -0.131 | 0.111 | -0.418 | -0.279 |
| **Fasting BG** | 0.204 | 0.327 | 1.000 | -0.096 | 0.365 | -0.461 | -0.295 | 0.512 | 0.467 | -0.079 | 0.399 | -0.523 | -0.189 |
| **Raw Brain** | -0.764 | 0.270 | -0.096 | 1.000 | 0.572 | -0.233 | 0.887 | 0.139 | 0.362 | -0.513 | 0.294 | 0.189 | 0.112 |
| **BW** | -0.525 | 0.145 | 0.365 | 0.572 | 1.000 | -0.905 | 0.228 | 0.115 | 0.776 | -0.783 | 0.620 | -0.217 | -0.086 |
| **Brain/BW** | 0.298 | -0.068 | -0.461 | -0.233 | -0.905 | 1.000 | 0.145 | -0.137 | -0.723 | 0.717 | -0.558 | 0.323 | 0.225 |
| **Brain/TL** | -0.632 | 0.323 | -0.295 | 0.887 | 0.228 | 0.145 | 1.000 | 0.123 | 0.133 | -0.186 | 0.190 | 0.280 | 0.176 |
| **GTT Peak Glc** | 0.074 | 0.888 | 0.512 | 0.139 | 0.115 | -0.137 | 0.123 | 1.000 | 0.070 | -0.097 | 0.071 | -0.504 | -0.429 |
| **Raw Heart** | -0.218 | 0.075 | 0.467 | 0.362 | 0.776 | -0.723 | 0.133 | 0.070 | 1.000 | -0.229 | 0.959 | -0.243 | -0.163 |
| **Heart/BW** | 0.608 | -0.131 | -0.079 | -0.513 | -0.783 | 0.717 | -0.186 | -0.097 | -0.229 | 1.000 | -0.017 | 0.050 | -0.001 |
| **Heart/TL** | -0.130 | 0.111 | 0.399 | 0.294 | 0.620 | -0.558 | 0.190 | 0.071 | 0.959 | -0.017 | 1.000 | -0.226 | -0.158 |
| **Fasting BOHB** | -0.432 | -0.418 | -0.523 | 0.189 | -0.217 | 0.323 | 0.280 | -0.504 | -0.243 | 0.050 | -0.226 | 1.000 | 0.687 |
| **BOHB/AcAc** | -0.259 | -0.279 | -0.189 | 0.112 | -0.086 | 0.225 | 0.176 | -0.429 | -0.163 | -0.001 | -0.158 | 0.687 | 1.000 |

**Supplemental Table 4:**
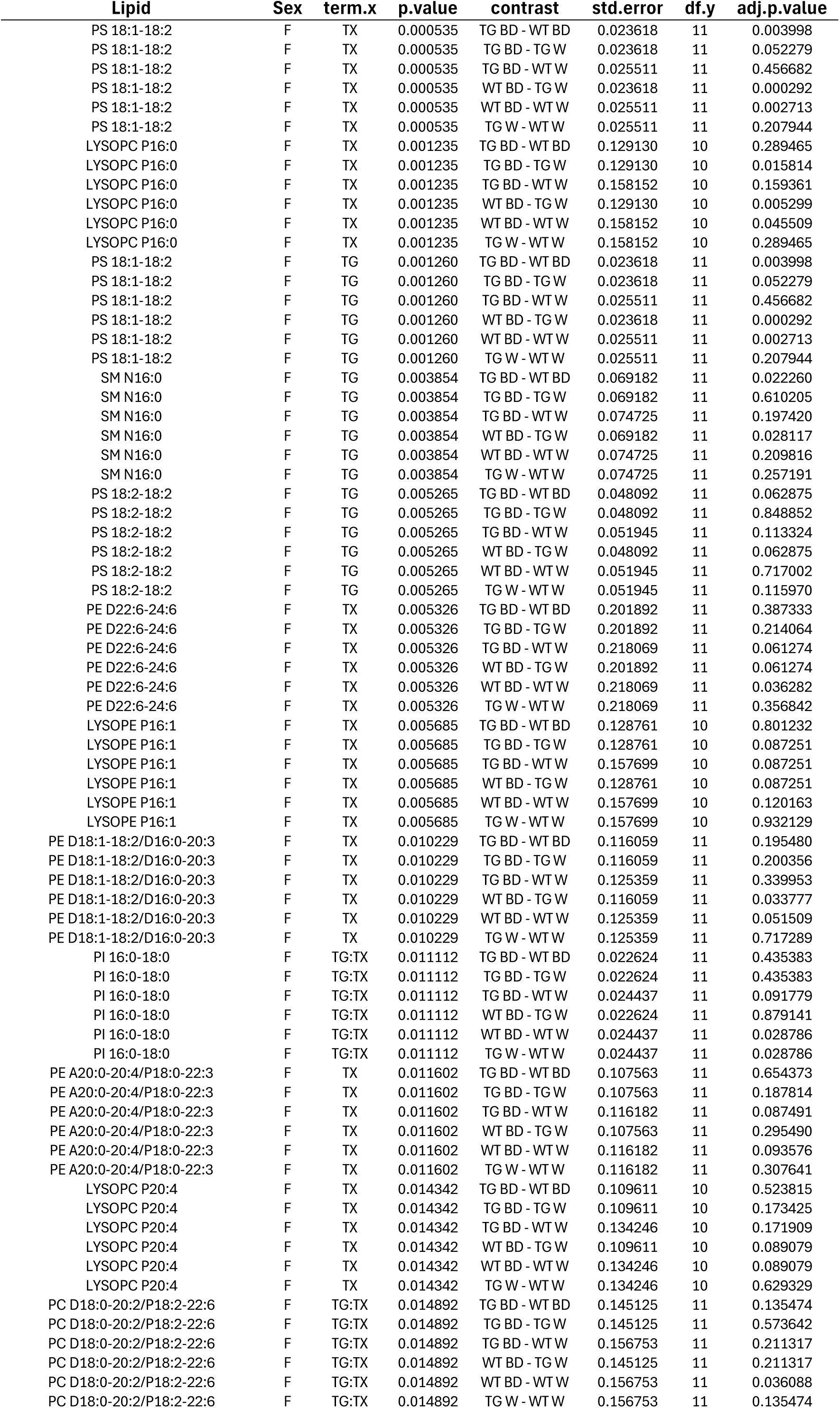
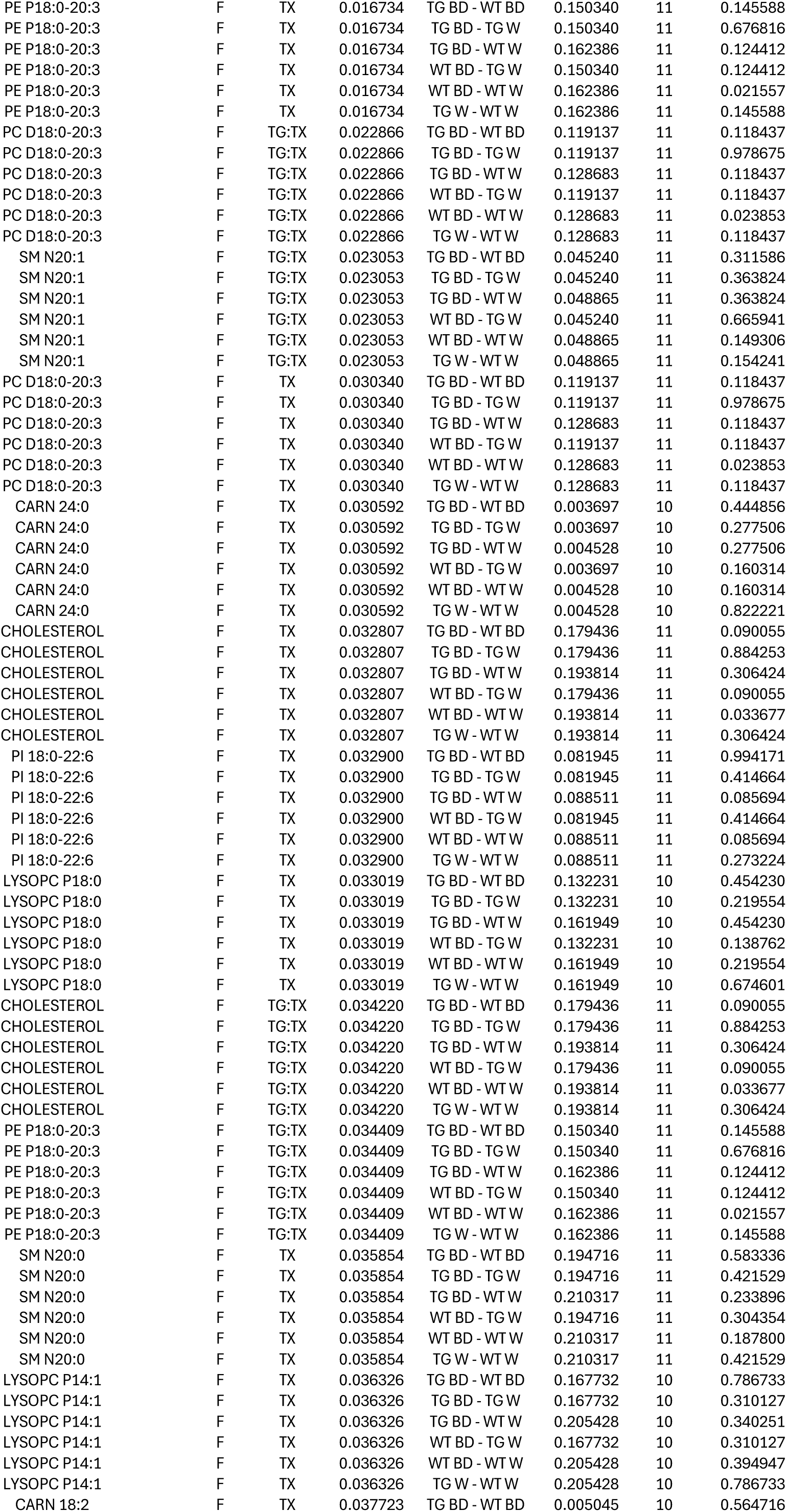
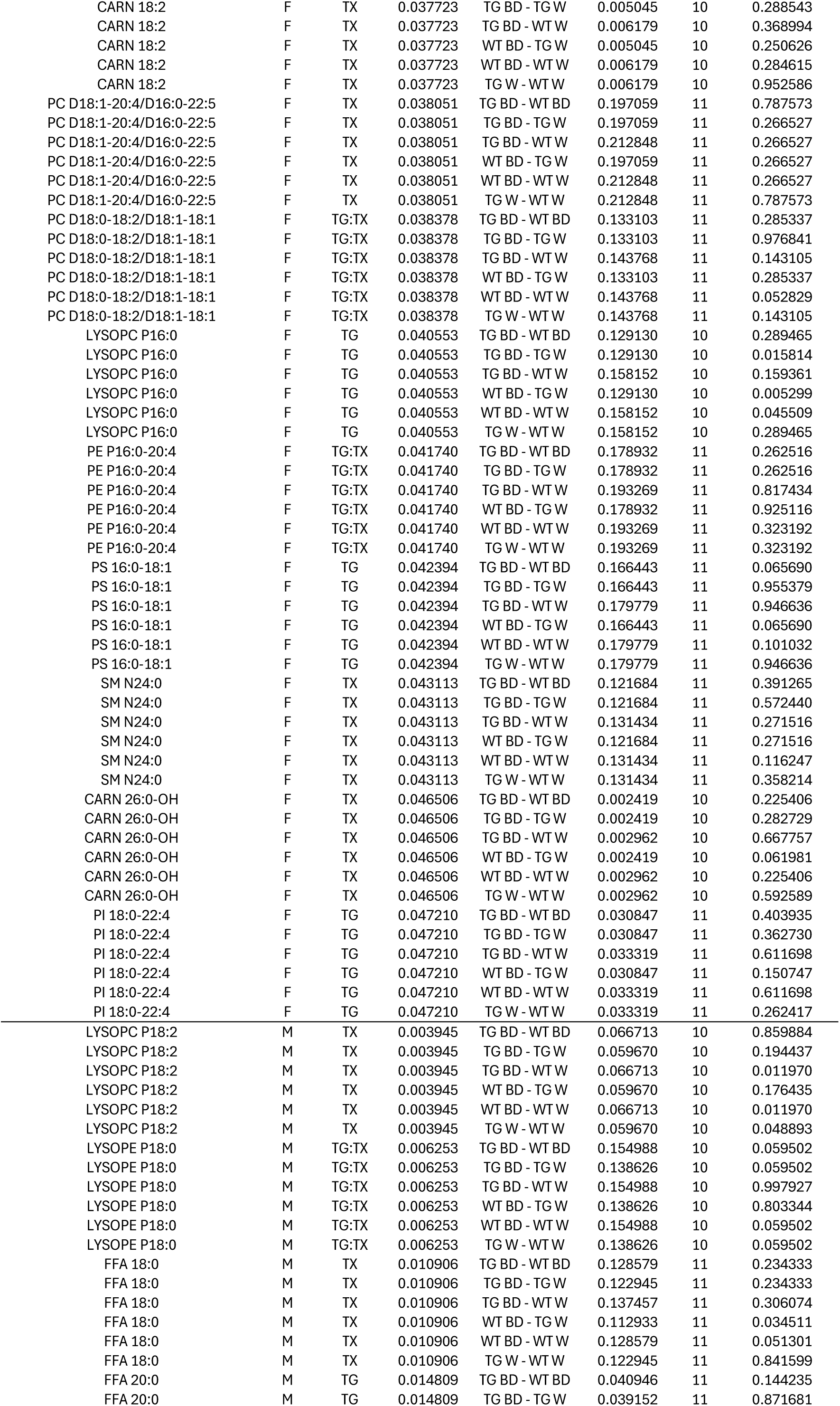
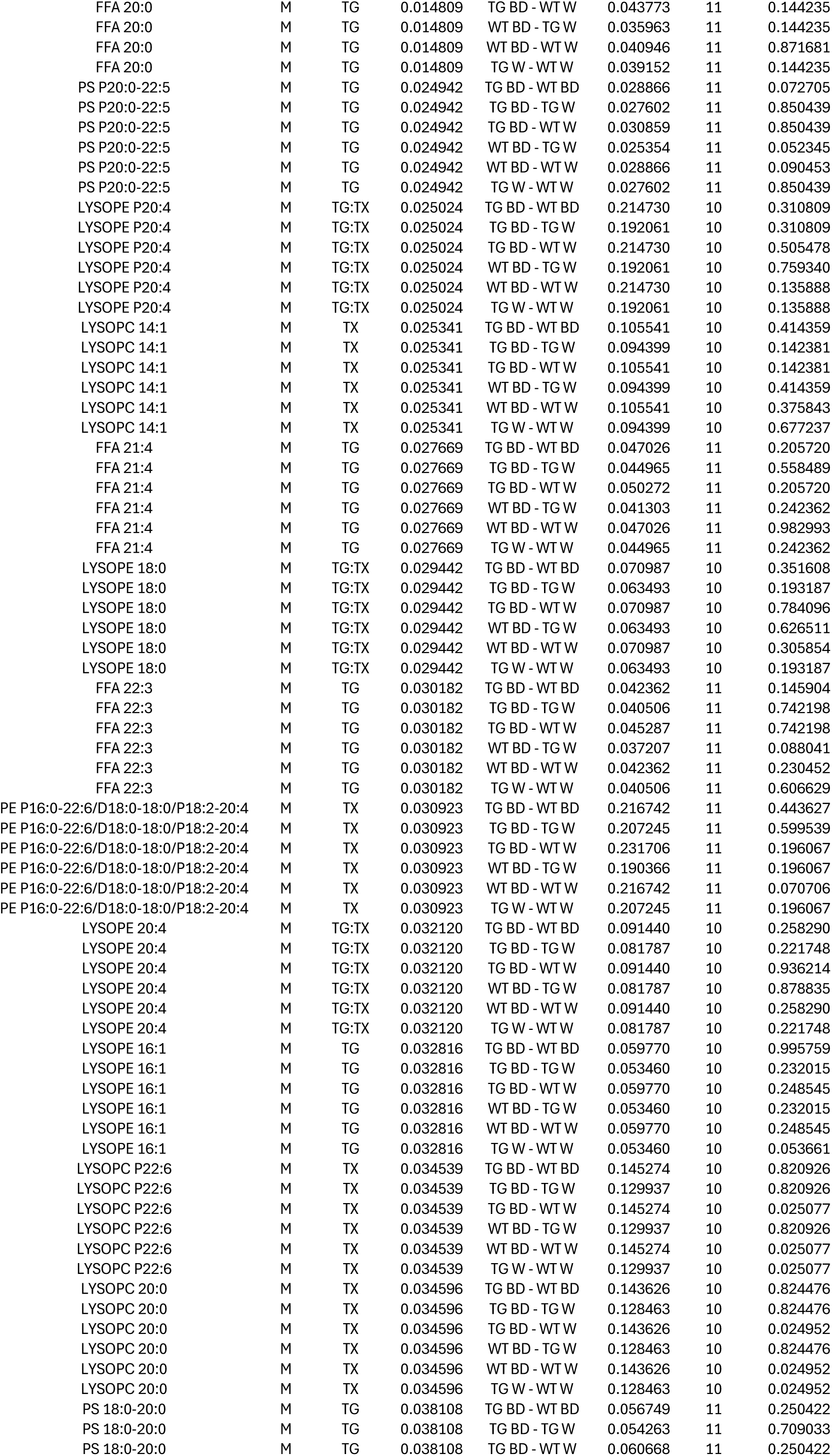
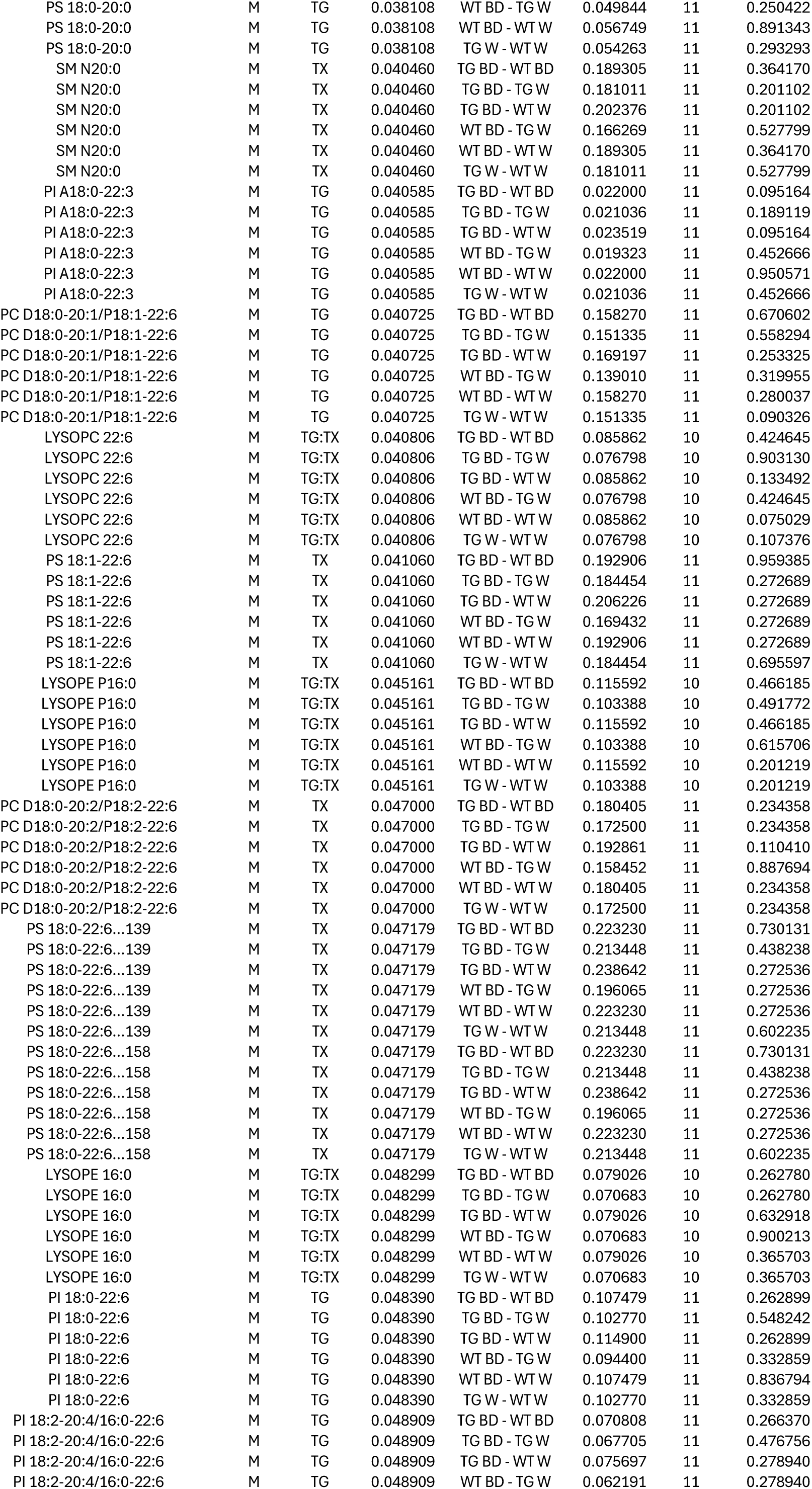
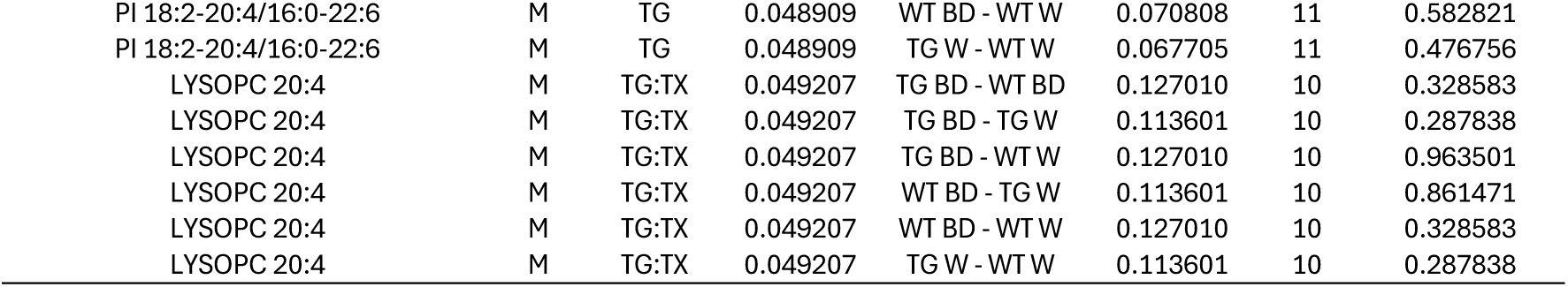
ANOVA results of meaningfully changed lipids in the brain. Absolute lipid concentrations in the brain were measured and compared within female or male cohorts. Data reflect two-way ANOVA with estimated marginal means and Benjamini-Hochberg false-discovery. term.x = specific model of interaction being tested, p.value = raw p-value for the interaction following ANOVA, contrast = specific pairwise comparison performed, std.error = standard error of estimated contrast, df.y = degrees of freedom associated with statistical test, adj.p.value = multiple-testing correction of p-value (q), F = female, M = male TG = transgenic, WT = nontransgenic, TX = treatment, W = water, BD = 1,3-butanediol, PC = phosphatidylcholine, PE = phosphatidylethanolamine, PA = phosphatidic acid, PI = phosphatidylinositol, PS = phosphatidylserine, SM = sphingomyelin, CBS = cerebroside, FFA = free fatty-acid, LPC = lysoPC, LPE = lysoPE, Car = acylcarnitine.

| Lipid | Sex | term.x | p.value | contrast | std.error | df.y | adj.p.value |
| --- | --- | --- | --- | --- | --- | --- | --- |
| PS 18:1-18:2 | F | TX | 0.000535 | TG BD - WT BD | 0.023618 | 11 | 0.003998 |
| PS 18:1-18:2 | F | TX | 0.000535 | TG BD - TG W | 0.023618 | 11 | 0.052279 |
| PS 18:1-18:2 | F | TX | 0.000535 | TG BD - WT W | 0.025511 | 11 | 0.456682 |
| PS 18:1-18:2 | F | TX | 0.000535 | WT BD - TG W | 0.023618 | 11 | 0.000292 |
| PS 18:1-18:2 | F | TX | 0.000535 | WT BD - WT W | 0.025511 | 11 | 0.002713 |
| PS 18:1-18:2 | F | TX | 0.000535 | TG W - WT W | 0.025511 | 11 | 0.207944 |
| LYSOPC P16:0 | F | TX | 0.001235 | TG BD - WT BD | 0.129130 | 10 | 0.289465 |
| LYSOPC P16:0 | F | TX | 0.001235 | TG BD - TG W | 0.129130 | 10 | 0.015814 |
| LYSOPC P16:0 | F | TX | 0.001235 | TG BD - WT W | 0.158152 | 10 | 0.159361 |
| LYSOPC P16:0 | F | TX | 0.001235 | WT BD - TG W | 0.129130 | 10 | 0.005299 |
| LYSOPC P16:0 | F | TX | 0.001235 | WT BD - WT W | 0.158152 | 10 | 0.045509 |
| LYSOPC P16:0 | F | TX | 0.001235 | TG W - WT W | 0.158152 | 10 | 0.289465 |
| PS 18:1-18:2 | F | TG | 0.001260 | TG BD - WT BD | 0.023618 | 11 | 0.003998 |
| PS 18:1-18:2 | F | TG | 0.001260 | TG BD - TG W | 0.023618 | 11 | 0.052279 |
| PS 18:1-18:2 | F | TG | 0.001260 | TG BD - WT W | 0.025511 | 11 | 0.456682 |
| PS 18:1-18:2 | F | TG | 0.001260 | WT BD - TG W | 0.023618 | 11 | 0.000292 |
| PS 18:1-18:2 | F | TG | 0.001260 | WT BD - WT W | 0.025511 | 11 | 0.002713 |
| PS 18:1-18:2 | F | TG | 0.001260 | TG W - WT W | 0.025511 | 11 | 0.207944 |
| SM N16:0 | F | TG | 0.003854 | TG BD - WT BD | 0.069182 | 11 | 0.022260 |
| SM N16:0 | F | TG | 0.003854 | TG BD - TG W | 0.069182 | 11 | 0.610205 |
| SM N16:0 | F | TG | 0.003854 | TG BD - WT W | 0.074725 | 11 | 0.197420 |
| SM N16:0 | F | TG | 0.003854 | WT BD - TG W | 0.069182 | 11 | 0.028117 |
| SM N16:0 | F | TG | 0.003854 | WT BD - WT W | 0.074725 | 11 | 0.209816 |
| SM N16:0 | F | TG | 0.003854 | TG W - WT W | 0.074725 | 11 | 0.257191 |
| PS 18:2-18:2 | F | TG | 0.005265 | TG BD - WT BD | 0.048092 | 11 | 0.062875 |
| PS 18:2-18:2 | F | TG | 0.005265 | TG BD - TG W | 0.048092 | 11 | 0.848852 |
| PS 18:2-18:2 | F | TG | 0.005265 | TG BD - WT W | 0.051945 | 11 | 0.113324 |
| PS 18:2-18:2 | F | TG | 0.005265 | WT BD - TG W | 0.048092 | 11 | 0.062875 |
| PS 18:2-18:2 | F | TG | 0.005265 | WT BD - WT W | 0.051945 | 11 | 0.717002 |
| PS 18:2-18:2 | F | TG | 0.005265 | TG W - WT W | 0.051945 | 11 | 0.115970 |
| PE D22:6-24:6 | F | TX | 0.005326 | TG BD - WT BD | 0.201892 | 11 | 0.387333 |
| PE D22:6-24:6 | F | TX | 0.005326 | TG BD - TG W | 0.201892 | 11 | 0.214064 |
| PE D22:6-24:6 | F | TX | 0.005326 | TG BD - WT W | 0.218069 | 11 | 0.061274 |
| PE D22:6-24:6 | F | TX | 0.005326 | WT BD - TG W | 0.201892 | 11 | 0.061274 |
| PE D22:6-24:6 | F | TX | 0.005326 | WT BD - WT W | 0.218069 | 11 | 0.036282 |
| PE D22:6-24:6 | F | TX | 0.005326 | TG W - WT W | 0.218069 | 11 | 0.356842 |
| LYSOPE P16:1 | F | TX | 0.005685 | TG BD - WT BD | 0.128761 | 10 | 0.801232 |
| LYSOPE P16:1 | F | TX | 0.005685 | TG BD - TG W | 0.128761 | 10 | 0.087251 |
| LYSOPE P16:1 | F | TX | 0.005685 | TG BD - WT W | 0.157699 | 10 | 0.087251 |
| LYSOPE P16:1 | F | TX | 0.005685 | WT BD - TG W | 0.128761 | 10 | 0.087251 |
| LYSOPE P16:1 | F | TX | 0.005685 | WT BD - WT W | 0.157699 | 10 | 0.120163 |
| LYSOPE P16:1 | F | TX | 0.005685 | TG W - WT W | 0.157699 | 10 | 0.932129 |
| PE D18:1-18:2/D16:0-20:3 | F | TX | 0.010229 | TG BD - WT BD | 0.116059 | 11 | 0.195480 |
| PE D18:1-18:2/D16:0-20:3 | F | TX | 0.010229 | TG BD - TG W | 0.116059 | 11 | 0.200356 |
| PE D18:1-18:2/D16:0-20:3 | F | TX | 0.010229 | TG BD - WT W | 0.125359 | 11 | 0.339953 |
| PE D18:1-18:2/D16:0-20:3 | F | TX | 0.010229 | WT BD - TG W | 0.116059 | 11 | 0.033777 |
| PE D18:1-18:2/D16:0-20:3 | F | TX | 0.010229 | WT BD - WT W | 0.125359 | 11 | 0.051509 |
| PE D18:1-18:2/D16:0-20:3 | F | TX | 0.010229 | TG W - WT W | 0.125359 | 11 | 0.717289 |
| PI 16:0-18:0 | F | TG:TX | 0.011112 | TG BD - WT BD | 0.022624 | 11 | 0.435383 |
| PI 16:0-18:0 | F | TG:TX | 0.011112 | TG BD - TG W | 0.022624 | 11 | 0.435383 |
| PI 16:0-18:0 | F | TG:TX | 0.011112 | TG BD - WT W | 0.024437 | 11 | 0.091779 |
| PI 16:0-18:0 | F | TG:TX | 0.011112 | WT BD - TG W | 0.022624 | 11 | 0.879141 |
| PI 16:0-18:0 | F | TG:TX | 0.011112 | WT BD - WT W | 0.024437 | 11 | 0.028786 |
| PI 16:0-18:0 | F | TG:TX | 0.011112 | TG W - WT W | 0.024437 | 11 | 0.028786 |
| PE A20:0-20:4/P18:0-22:3 | F | TX | 0.011602 | TG BD - WT BD | 0.107563 | 11 | 0.654373 |
| PE A20:0-20:4/P18:0-22:3 | F | TX | 0.011602 | TG BD - TG W | 0.107563 | 11 | 0.187814 |
| PE A20:0-20:4/P18:0-22:3 | F | TX | 0.011602 | TG BD - WT W | 0.116182 | 11 | 0.087491 |
| PE A20:0-20:4/P18:0-22:3 | F | TX | 0.011602 | WT BD - TG W | 0.107563 | 11 | 0.295490 |
| PE A20:0-20:4/P18:0-22:3 | F | TX | 0.011602 | WT BD - WT W | 0.116182 | 11 | 0.093576 |
| PE A20:0-20:4/P18:0-22:3 | F | TX | 0.011602 | TG W - WT W | 0.116182 | 11 | 0.307641 |
| LYSOPC P20:4 | F | TX | 0.014342 | TG BD - WT BD | 0.109611 | 10 | 0.523815 |
| LYSOPC P20:4 | F | TX | 0.014342 | TG BD - TG W | 0.109611 | 10 | 0.173425 |
| LYSOPC P20:4 | F | TX | 0.014342 | TG BD - WT W | 0.134246 | 10 | 0.171909 |
| LYSOPC P20:4 | F | TX | 0.014342 | WT BD - TG W | 0.109611 | 10 | 0.089079 |
| LYSOPC P20:4 | F | TX | 0.014342 | WT BD - WT W | 0.134246 | 10 | 0.089079 |
| LYSOPC P20:4 | F | TX | 0.014342 | TG W - WT W | 0.134246 | 10 | 0.629329 |
| PC D18:0-20:2/P18:2-22:6 | F | TG:TX | 0.014892 | TG BD - WT BD | 0.145125 | 11 | 0.135474 |
| PC D18:0-20:2/P18:2-22:6 | F | TG:TX | 0.014892 | TG BD - TG W | 0.145125 | 11 | 0.573642 |
| PC D18:0-20:2/P18:2-22:6 | F | TG:TX | 0.014892 | TG BD - WT W | 0.156753 | 11 | 0.211317 |
| PC D18:0-20:2/P18:2-22:6 | F | TG:TX | 0.014892 | WT BD - TG W | 0.145125 | 11 | 0.211317 |
| PC D18:0-20:2/P18:2-22:6 | F | TG:TX | 0.014892 | WT BD - WT W | 0.156753 | 11 | 0.036088 |
| PC D18:0-20:2/P18:2-22:6 | F | TG:TX | 0.014892 | TG W - WT W | 0.156753 | 11 | 0.135474 |
| PE P18:0-20:3 | F | TX | 0.016734 | TG BD - WT BD | 0.150340 | 11 | 0.145588 |
| PE P18:0-20:3 | F | TX | 0.016734 | TG BD - TG W | 0.150340 | 11 | 0.676816 |
| PE P18:0-20:3 | F | TX | 0.016734 | TG BD - WT W | 0.162386 | 11 | 0.124412 |
| PE P18:0-20:3 | F | TX | 0.016734 | WT BD - TG W | 0.150340 | 11 | 0.124412 |
| PE P18:0-20:3 | F | TX | 0.016734 | WT BD - WT W | 0.162386 | 11 | 0.021557 |
| PE P18:0-20:3 | F | TX | 0.016734 | TG W - WT W | 0.162386 | 11 | 0.145588 |
| PC D18:0-20:3 | F | TG:TX | 0.022866 | TG BD - WT BD | 0.119137 | 11 | 0.118437 |
| PC D18:0-20:3 | F | TG:TX | 0.022866 | TG BD - TG W | 0.119137 | 11 | 0.978675 |
| PC D18:0-20:3 | F | TG:TX | 0.022866 | TG BD - WT W | 0.128683 | 11 | 0.118437 |
| PC D18:0-20:3 | F | TG:TX | 0.022866 | WT BD - TG W | 0.119137 | 11 | 0.118437 |
| PC D18:0-20:3 | F | TG:TX | 0.022866 | WT BD - WT W | 0.128683 | 11 | 0.023853 |
| PC D18:0-20:3 | F | TG:TX | 0.022866 | TG W - WT W | 0.128683 | 11 | 0.118437 |
| SM N20:1 | F | TG:TX | 0.023053 | TG BD - WT BD | 0.045240 | 11 | 0.311586 |
| SM N20:1 | F | TG:TX | 0.023053 | TG BD - TG W | 0.045240 | 11 | 0.363824 |
| SM N20:1 | F | TG:TX | 0.023053 | TG BD - WT W | 0.048865 | 11 | 0.363824 |
| SM N20:1 | F | TG:TX | 0.023053 | WT BD - TG W | 0.045240 | 11 | 0.665941 |
| SM N20:1 | F | TG:TX | 0.023053 | WT BD - WT W | 0.048865 | 11 | 0.149306 |
| SM N20:1 | F | TG:TX | 0.023053 | TG W - WT W | 0.048865 | 11 | 0.154241 |
| PC D18:0-20:3 | F | TX | 0.030340 | TG BD - WT BD | 0.119137 | 11 | 0.118437 |
| PC D18:0-20:3 | F | TX | 0.030340 | TG BD - TG W | 0.119137 | 11 | 0.978675 |
| PC D18:0-20:3 | F | TX | 0.030340 | TG BD - WT W | 0.128683 | 11 | 0.118437 |
| PC D18:0-20:3 | F | TX | 0.030340 | WT BD - TG W | 0.119137 | 11 | 0.118437 |
| PC D18:0-20:3 | F | TX | 0.030340 | WT BD - WT W | 0.128683 | 11 | 0.023853 |
| PC D18:0-20:3 | F | TX | 0.030340 | TG W - WT W | 0.128683 | 11 | 0.118437 |
| CARN 24:0 | F | TX | 0.030592 | TG BD - WT BD | 0.003697 | 10 | 0.444856 |
| CARN 24:0 | F | TX | 0.030592 | TG BD - TG W | 0.003697 | 10 | 0.277506 |
| CARN 24:0 | F | TX | 0.030592 | TG BD - WT W | 0.004528 | 10 | 0.277506 |
| CARN 24:0 | F | TX | 0.030592 | WT BD - TG W | 0.003697 | 10 | 0.160314 |
| CARN 24:0 | F | TX | 0.030592 | WT BD - WT W | 0.004528 | 10 | 0.160314 |
| CARN 24:0 | F | TX | 0.030592 | TG W - WT W | 0.004528 | 10 | 0.822221 |
| CHOLESTEROL | F | TX | 0.032807 | TG BD - WT BD | 0.179436 | 11 | 0.090055 |
| CHOLESTEROL | F | TX | 0.032807 | TG BD - TG W | 0.179436 | 11 | 0.884253 |
| CHOLESTEROL | F | TX | 0.032807 | TG BD - WT W | 0.193814 | 11 | 0.306424 |
| CHOLESTEROL | F | TX | 0.032807 | WT BD - TG W | 0.179436 | 11 | 0.090055 |
| CHOLESTEROL | F | TX | 0.032807 | WT BD - WT W | 0.193814 | 11 | 0.033677 |
| CHOLESTEROL | F | TX | 0.032807 | TG W - WT W | 0.193814 | 11 | 0.306424 |
| PI 18:0-22:6 | F | TX | 0.032900 | TG BD - WT BD | 0.081945 | 11 | 0.994171 |
| PI 18:0-22:6 | F | TX | 0.032900 | TG BD - TG W | 0.081945 | 11 | 0.414664 |
| PI 18:0-22:6 | F | TX | 0.032900 | TG BD - WT W | 0.088511 | 11 | 0.085694 |
| PI 18:0-22:6 | F | TX | 0.032900 | WT BD - TG W | 0.081945 | 11 | 0.414664 |
| PI 18:0-22:6 | F | TX | 0.032900 | WT BD - WT W | 0.088511 | 11 | 0.085694 |
| PI 18:0-22:6 | F | TX | 0.032900 | TG W - WT W | 0.088511 | 11 | 0.273224 |
| LYSOPC P18:0 | F | TX | 0.033019 | TG BD - WT BD | 0.132231 | 10 | 0.454230 |
| LYSOPC P18:0 | F | TX | 0.033019 | TG BD - TG W | 0.132231 | 10 | 0.219554 |
| LYSOPC P18:0 | F | TX | 0.033019 | TG BD - WT W | 0.161949 | 10 | 0.454230 |
| LYSOPC P18:0 | F | TX | 0.033019 | WT BD - TG W | 0.132231 | 10 | 0.138762 |
| LYSOPC P18:0 | F | TX | 0.033019 | WT BD - WT W | 0.161949 | 10 | 0.219554 |
| LYSOPC P18:0 | F | TX | 0.033019 | TG W - WT W | 0.161949 | 10 | 0.674601 |
| CHOLESTEROL | F | TG:TX | 0.034220 | TG BD - WT BD | 0.179436 | 11 | 0.090055 |
| CHOLESTEROL | F | TG:TX | 0.034220 | TG BD - TG W | 0.179436 | 11 | 0.884253 |
| CHOLESTEROL | F | TG:TX | 0.034220 | TG BD - WT W | 0.193814 | 11 | 0.306424 |
| CHOLESTEROL | F | TG:TX | 0.034220 | WT BD - TG W | 0.179436 | 11 | 0.090055 |
| CHOLESTEROL | F | TG:TX | 0.034220 | WT BD - WT W | 0.193814 | 11 | 0.033677 |
| CHOLESTEROL | F | TG:TX | 0.034220 | TG W - WT W | 0.193814 | 11 | 0.306424 |
| PE P18:0-20:3 | F | TG:TX | 0.034409 | TG BD - WT BD | 0.150340 | 11 | 0.145588 |
| PE P18:0-20:3 | F | TG:TX | 0.034409 | TG BD - TG W | 0.150340 | 11 | 0.676816 |
| PE P18:0-20:3 | F | TG:TX | 0.034409 | TG BD - WT W | 0.162386 | 11 | 0.124412 |
| PE P18:0-20:3 | F | TG:TX | 0.034409 | WT BD - TG W | 0.150340 | 11 | 0.124412 |
| PE P18:0-20:3 | F | TG:TX | 0.034409 | WT BD - WT W | 0.162386 | 11 | 0.021557 |
| PE P18:0-20:3 | F | TG:TX | 0.034409 | TG W - WT W | 0.162386 | 11 | 0.145588 |
| SM N20:0 | F | TX | 0.035854 | TG BD - WT BD | 0.194716 | 11 | 0.583336 |
| SM N20:0 | F | TX | 0.035854 | TG BD - TG W | 0.194716 | 11 | 0.421529 |
| SM N20:0 | F | TX | 0.035854 | TG BD - WT W | 0.210317 | 11 | 0.233896 |
| SM N20:0 | F | TX | 0.035854 | WT BD - TG W | 0.194716 | 11 | 0.304354 |
| SM N20:0 | F | TX | 0.035854 | WT BD - WT W | 0.210317 | 11 | 0.187800 |
| SM N20:0 | F | TX | 0.035854 | TG W - WT W | 0.210317 | 11 | 0.421529 |
| LYSOPC P14:1 | F | TX | 0.036326 | TG BD - WT BD | 0.167732 | 10 | 0.786733 |
| LYSOPC P14:1 | F | TX | 0.036326 | TG BD - TG W | 0.167732 | 10 | 0.310127 |
| LYSOPC P14:1 | F | TX | 0.036326 | TG BD - WT W | 0.205428 | 10 | 0.340251 |
| LYSOPC P14:1 | F | TX | 0.036326 | WT BD - TG W | 0.167732 | 10 | 0.310127 |
| LYSOPC P14:1 | F | TX | 0.036326 | WT BD - WT W | 0.205428 | 10 | 0.394947 |
| LYSOPC P14:1 | F | TX | 0.036326 | TG W - WT W | 0.205428 | 10 | 0.786733 |
| CARN 18:2 | F | TX | 0.037723 | TG BD - WT BD | 0.005045 | 10 | 0.564716 |
| CARN 18:2 | F | TX | 0.037723 | TG BD - TG W | 0.005045 | 10 | 0.288543 |
| CARN 18:2 | F | TX | 0.037723 | TG BD - WT W | 0.006179 | 10 | 0.368994 |
| CARN 18:2 | F | TX | 0.037723 | WT BD - TG W | 0.005045 | 10 | 0.250626 |
| CARN 18:2 | F | TX | 0.037723 | WT BD - WT W | 0.006179 | 10 | 0.284615 |
| CARN 18:2 | F | TX | 0.037723 | TG W - WT W | 0.006179 | 10 | 0.952586 |
| PC D18:1-20:4/D16:0-22:5 | F | TX | 0.038051 | TG BD - WT BD | 0.197059 | 11 | 0.787573 |
| PC D18:1-20:4/D16:0-22:5 | F | TX | 0.038051 | TG BD - TG W | 0.197059 | 11 | 0.266527 |
| PC D18:1-20:4/D16:0-22:5 | F | TX | 0.038051 | TG BD - WT W | 0.212848 | 11 | 0.266527 |
| PC D18:1-20:4/D16:0-22:5 | F | TX | 0.038051 | WT BD - TG W | 0.197059 | 11 | 0.266527 |
| PC D18:1-20:4/D16:0-22:5 | F | TX | 0.038051 | WT BD - WT W | 0.212848 | 11 | 0.266527 |
| PC D18:1-20:4/D16:0-22:5 | F | TX | 0.038051 | TG W - WT W | 0.212848 | 11 | 0.787573 |
| PC D18:0-18:2/D18:1-18:1 | F | TG:TX | 0.038378 | TG BD - WT BD | 0.133103 | 11 | 0.285337 |
| PC D18:0-18:2/D18:1-18:1 | F | TG:TX | 0.038378 | TG BD - TG W | 0.133103 | 11 | 0.976841 |
| PC D18:0-18:2/D18:1-18:1 | F | TG:TX | 0.038378 | TG BD - WT W | 0.143768 | 11 | 0.143105 |
| PC D18:0-18:2/D18:1-18:1 | F | TG:TX | 0.038378 | WT BD - TG W | 0.133103 | 11 | 0.285337 |
| PC D18:0-18:2/D18:1-18:1 | F | TG:TX | 0.038378 | WT BD - WT W | 0.143768 | 11 | 0.052829 |
| PC D18:0-18:2/D18:1-18:1 | F | TG:TX | 0.038378 | TG W - WT W | 0.143768 | 11 | 0.143105 |
| LYSOPC P16:0 | F | TG | 0.040553 | TG BD - WT BD | 0.129130 | 10 | 0.289465 |
| LYSOPC P16:0 | F | TG | 0.040553 | TG BD - TG W | 0.129130 | 10 | 0.015814 |
| LYSOPC P16:0 | F | TG | 0.040553 | TG BD - WT W | 0.158152 | 10 | 0.159361 |
| LYSOPC P16:0 | F | TG | 0.040553 | WT BD - TG W | 0.129130 | 10 | 0.005299 |
| LYSOPC P16:0 | F | TG | 0.040553 | WT BD - WT W | 0.158152 | 10 | 0.045509 |
| LYSOPC P16:0 | F | TG | 0.040553 | TG W - WT W | 0.158152 | 10 | 0.289465 |
| PE P16:0-20:4 | F | TG:TX | 0.041740 | TG BD - WT BD | 0.178932 | 11 | 0.262516 |
| PE P16:0-20:4 | F | TG:TX | 0.041740 | TG BD - TG W | 0.178932 | 11 | 0.262516 |
| PE P16:0-20:4 | F | TG:TX | 0.041740 | TG BD - WT W | 0.193269 | 11 | 0.817434 |
| PE P16:0-20:4 | F | TG:TX | 0.041740 | WT BD - TG W | 0.178932 | 11 | 0.925116 |
| PE P16:0-20:4 | F | TG:TX | 0.041740 | WT BD - WT W | 0.193269 | 11 | 0.323192 |
| PE P16:0-20:4 | F | TG:TX | 0.041740 | TG W - WT W | 0.193269 | 11 | 0.323192 |
| PS 16:0-18:1 | F | TG | 0.042394 | TG BD - WT BD | 0.166443 | 11 | 0.065690 |
| PS 16:0-18:1 | F | TG | 0.042394 | TG BD - TG W | 0.166443 | 11 | 0.955379 |
| PS 16:0-18:1 | F | TG | 0.042394 | TG BD - WT W | 0.179779 | 11 | 0.946636 |
| PS 16:0-18:1 | F | TG | 0.042394 | WT BD - TG W | 0.166443 | 11 | 0.065690 |
| PS 16:0-18:1 | F | TG | 0.042394 | WT BD - WT W | 0.179779 | 11 | 0.101032 |
| PS 16:0-18:1 | F | TG | 0.042394 | TG W - WT W | 0.179779 | 11 | 0.946636 |
| SM N24:0 | F | TX | 0.043113 | TG BD - WT BD | 0.121684 | 11 | 0.391265 |
| SM N24:0 | F | TX | 0.043113 | TG BD - TG W | 0.121684 | 11 | 0.572440 |
| SM N24:0 | F | TX | 0.043113 | TG BD - WT W | 0.131434 | 11 | 0.271516 |
| SM N24:0 | F | TX | 0.043113 | WT BD - TG W | 0.121684 | 11 | 0.271516 |
| SM N24:0 | F | TX | 0.043113 | WT BD - WT W | 0.131434 | 11 | 0.116247 |
| SM N24:0 | F | TX | 0.043113 | TG W - WT W | 0.131434 | 11 | 0.358214 |
| CARN 26:0-OH | F | TX | 0.046506 | TG BD - WT BD | 0.002419 | 10 | 0.225406 |
| CARN 26:0-OH | F | TX | 0.046506 | TG BD - TG W | 0.002419 | 10 | 0.282729 |
| CARN 26:0-OH | F | TX | 0.046506 | TG BD - WT W | 0.002962 | 10 | 0.667757 |
| CARN 26:0-OH | F | TX | 0.046506 | WT BD - TG W | 0.002419 | 10 | 0.061981 |
| CARN 26:0-OH | F | TX | 0.046506 | WT BD - WT W | 0.002962 | 10 | 0.225406 |
| CARN 26:0-OH | F | TX | 0.046506 | TG W - WT W | 0.002962 | 10 | 0.592589 |
| PI 18:0-22:4 | F | TG | 0.047210 | TG BD - WT BD | 0.030847 | 11 | 0.403935 |
| PI 18:0-22:4 | F | TG | 0.047210 | TG BD - TG W | 0.030847 | 11 | 0.362730 |
| PI 18:0-22:4 | F | TG | 0.047210 | TG BD - WT W | 0.033319 | 11 | 0.611698 |
| PI 18:0-22:4 | F | TG | 0.047210 | WT BD - TG W | 0.030847 | 11 | 0.150747 |
| PI 18:0-22:4 | F | TG | 0.047210 | WT BD - WT W | 0.033319 | 11 | 0.611698 |
| PI 18:0-22:4 | F | TG | 0.047210 | TG W - WT W | 0.033319 | 11 | 0.262417 |
| LYSOPC P18:2 | M | TX | 0.003945 | TG BD - WT BD | 0.066713 | 10 | 0.859884 |
| LYSOPC P18:2 | M | TX | 0.003945 | TG BD - TG W | 0.059670 | 10 | 0.194437 |
| LYSOPC P18:2 | M | TX | 0.003945 | TG BD - WT W | 0.066713 | 10 | 0.011970 |
| LYSOPC P18:2 | M | TX | 0.003945 | WT BD - TG W | 0.059670 | 10 | 0.176435 |
| LYSOPC P18:2 | M | TX | 0.003945 | WT BD - WT W | 0.066713 | 10 | 0.011970 |
| LYSOPC P18:2 | M | TX | 0.003945 | TG W - WT W | 0.059670 | 10 | 0.048893 |
| LYSOPE P18:0 | M | TG:TX | 0.006253 | TG BD - WT BD | 0.154988 | 10 | 0.059502 |
| LYSOPE P18:0 | M | TG:TX | 0.006253 | TG BD - TG W | 0.138626 | 10 | 0.059502 |
| LYSOPE P18:0 | M | TG:TX | 0.006253 | TG BD - WT W | 0.154988 | 10 | 0.997927 |
| LYSOPE P18:0 | M | TG:TX | 0.006253 | WT BD - TG W | 0.138626 | 10 | 0.803344 |
| LYSOPE P18:0 | M | TG:TX | 0.006253 | WT BD - WT W | 0.154988 | 10 | 0.059502 |
| LYSOPE P18:0 | M | TG:TX | 0.006253 | TG W - WT W | 0.138626 | 10 | 0.059502 |
| FFA 18:0 | M | TX | 0.010906 | TG BD - WT BD | 0.128579 | 11 | 0.234333 |
| FFA 18:0 | M | TX | 0.010906 | TG BD - TG W | 0.122945 | 11 | 0.234333 |
| FFA 18:0 | M | TX | 0.010906 | TG BD - WT W | 0.137457 | 11 | 0.306074 |
| FFA 18:0 | M | TX | 0.010906 | WT BD - TG W | 0.112933 | 11 | 0.034511 |
| FFA 18:0 | M | TX | 0.010906 | WT BD - WT W | 0.128579 | 11 | 0.051301 |
| FFA 18:0 | M | TX | 0.010906 | TG W - WT W | 0.122945 | 11 | 0.841599 |
| FFA 20:0 | M | TG | 0.014809 | TG BD - WT BD | 0.040946 | 11 | 0.144235 |
| FFA 20:0 | M | TG | 0.014809 | TG BD - TG W | 0.039152 | 11 | 0.871681 |
| FFA 20:0 | M | TG | 0.014809 | TG BD - WT W | 0.043773 | 11 | 0.144235 |
| FFA 20:0 | M | TG | 0.014809 | WT BD - TG W | 0.035963 | 11 | 0.144235 |
| FFA 20:0 | M | TG | 0.014809 | WT BD - WT W | 0.040946 | 11 | 0.871681 |
| FFA 20:0 | M | TG | 0.014809 | TG W - WT W | 0.039152 | 11 | 0.144235 |
| PS P20:0-22:5 | M | TG | 0.024942 | TG BD - WT BD | 0.028866 | 11 | 0.072705 |
| PS P20:0-22:5 | M | TG | 0.024942 | TG BD - TG W | 0.027602 | 11 | 0.850439 |
| PS P20:0-22:5 | M | TG | 0.024942 | TG BD - WT W | 0.030859 | 11 | 0.850439 |
| PS P20:0-22:5 | M | TG | 0.024942 | WT BD - TG W | 0.025354 | 11 | 0.052345 |
| PS P20:0-22:5 | M | TG | 0.024942 | WT BD - WT W | 0.028866 | 11 | 0.090453 |
| PS P20:0-22:5 | M | TG | 0.024942 | TG W - WT W | 0.027602 | 11 | 0.850439 |
| LYSOPE P20:4 | M | TG:TX | 0.025024 | TG BD - WT BD | 0.214730 | 10 | 0.310809 |
| LYSOPE P20:4 | M | TG:TX | 0.025024 | TG BD - TG W | 0.192061 | 10 | 0.310809 |
| LYSOPE P20:4 | M | TG:TX | 0.025024 | TG BD - WT W | 0.214730 | 10 | 0.505478 |
| LYSOPE P20:4 | M | TG:TX | 0.025024 | WT BD - TG W | 0.192061 | 10 | 0.759340 |
| LYSOPE P20:4 | M | TG:TX | 0.025024 | WT BD - WT W | 0.214730 | 10 | 0.135888 |
| LYSOPE P20:4 | M | TG:TX | 0.025024 | TG W - WT W | 0.192061 | 10 | 0.135888 |
| LYSOPC 14:1 | M | TX | 0.025341 | TG BD - WT BD | 0.105541 | 10 | 0.414359 |
| LYSOPC 14:1 | M | TX | 0.025341 | TG BD - TG W | 0.094399 | 10 | 0.142381 |
| LYSOPC 14:1 | M | TX | 0.025341 | TG BD - WT W | 0.105541 | 10 | 0.142381 |
| LYSOPC 14:1 | M | TX | 0.025341 | WT BD - TG W | 0.094399 | 10 | 0.414359 |
| LYSOPC 14:1 | M | TX | 0.025341 | WT BD - WT W | 0.105541 | 10 | 0.375843 |
| LYSOPC 14:1 | M | TX | 0.025341 | TG W - WT W | 0.094399 | 10 | 0.677237 |
| FFA 21:4 | M | TG | 0.027669 | TG BD - WT BD | 0.047026 | 11 | 0.205720 |
| FFA 21:4 | M | TG | 0.027669 | TG BD - TG W | 0.044965 | 11 | 0.558489 |
| FFA 21:4 | M | TG | 0.027669 | TG BD - WT W | 0.050272 | 11 | 0.205720 |
| FFA 21:4 | M | TG | 0.027669 | WT BD - TG W | 0.041303 | 11 | 0.242362 |
| FFA 21:4 | M | TG | 0.027669 | WT BD - WT W | 0.047026 | 11 | 0.982993 |
| FFA 21:4 | M | TG | 0.027669 | TG W - WT W | 0.044965 | 11 | 0.242362 |
| LYSOPE 18:0 | M | TG:TX | 0.029442 | TG BD - WT BD | 0.070987 | 10 | 0.351608 |
| LYSOPE 18:0 | M | TG:TX | 0.029442 | TG BD - TG W | 0.063493 | 10 | 0.193187 |
| LYSOPE 18:0 | M | TG:TX | 0.029442 | TG BD - WT W | 0.070987 | 10 | 0.784096 |
| LYSOPE 18:0 | M | TG:TX | 0.029442 | WT BD - TG W | 0.063493 | 10 | 0.626511 |
| LYSOPE 18:0 | M | TG:TX | 0.029442 | WT BD - WT W | 0.070987 | 10 | 0.305854 |
| LYSOPE 18:0 | M | TG:TX | 0.029442 | TG W - WT W | 0.063493 | 10 | 0.193187 |
| FFA 22:3 | M | TG | 0.030182 | TG BD - WT BD | 0.042362 | 11 | 0.145904 |
| FFA 22:3 | M | TG | 0.030182 | TG BD - TG W | 0.040506 | 11 | 0.742198 |
| FFA 22:3 | M | TG | 0.030182 | TG BD - WT W | 0.045287 | 11 | 0.742198 |
| FFA 22:3 | M | TG | 0.030182 | WT BD - TG W | 0.037207 | 11 | 0.088041 |
| FFA 22:3 | M | TG | 0.030182 | WT BD - WT W | 0.042362 | 11 | 0.230452 |
| FFA 22:3 | M | TG | 0.030182 | TG W - WT W | 0.040506 | 11 | 0.606629 |
| PE P16:0-22:6/D18:0-18:0/P18:2-20:4 | M | TX | 0.030923 | TG BD - WT BD | 0.216742 | 11 | 0.443627 |
| PE P16:0-22:6/D18:0-18:0/P18:2-20:4 | M | TX | 0.030923 | TG BD - TG W | 0.207245 | 11 | 0.599539 |
| PE P16:0-22:6/D18:0-18:0/P18:2-20:4 | M | TX | 0.030923 | TG BD - WT W | 0.231706 | 11 | 0.196067 |
| PE P16:0-22:6/D18:0-18:0/P18:2-20:4 | M | TX | 0.030923 | WT BD - TG W | 0.190366 | 11 | 0.196067 |
| PE P16:0-22:6/D18:0-18:0/P18:2-20:4 | M | TX | 0.030923 | WT BD - WT W | 0.216742 | 11 | 0.070706 |
| PE P16:0-22:6/D18:0-18:0/P18:2-20:4 | M | TX | 0.030923 | TG W - WT W | 0.207245 | 11 | 0.196067 |
| LYSOPE 20:4 | M | TG:TX | 0.032120 | TG BD - WT BD | 0.091440 | 10 | 0.258290 |
| LYSOPE 20:4 | M | TG:TX | 0.032120 | TG BD - TG W | 0.081787 | 10 | 0.221748 |
| LYSOPE 20:4 | M | TG:TX | 0.032120 | TG BD - WT W | 0.091440 | 10 | 0.936214 |
| LYSOPE 20:4 | M | TG:TX | 0.032120 | WT BD - TG W | 0.081787 | 10 | 0.878835 |
| LYSOPE 20:4 | M | TG:TX | 0.032120 | WT BD - WT W | 0.091440 | 10 | 0.258290 |
| LYSOPE 20:4 | M | TG:TX | 0.032120 | TG W - WT W | 0.081787 | 10 | 0.221748 |
| LYSOPE 16:1 | M | TG | 0.032816 | TG BD - WT BD | 0.059770 | 10 | 0.995759 |
| LYSOPE 16:1 | M | TG | 0.032816 | TG BD - TG W | 0.053460 | 10 | 0.232015 |
| LYSOPE 16:1 | M | TG | 0.032816 | TG BD - WT W | 0.059770 | 10 | 0.248545 |
| LYSOPE 16:1 | M | TG | 0.032816 | WT BD - TG W | 0.053460 | 10 | 0.232015 |
| LYSOPE 16:1 | M | TG | 0.032816 | WT BD - WT W | 0.059770 | 10 | 0.248545 |
| LYSOPE 16:1 | M | TG | 0.032816 | TG W - WT W | 0.053460 | 10 | 0.053661 |
| LYSOPC P22:6 | M | TX | 0.034539 | TG BD - WT BD | 0.145274 | 10 | 0.820926 |
| LYSOPC P22:6 | M | TX | 0.034539 | TG BD - TG W | 0.129937 | 10 | 0.820926 |
| LYSOPC P22:6 | M | TX | 0.034539 | TG BD - WT W | 0.145274 | 10 | 0.025077 |
| LYSOPC P22:6 | M | TX | 0.034539 | WT BD - TG W | 0.129937 | 10 | 0.820926 |
| LYSOPC P22:6 | M | TX | 0.034539 | WT BD - WT W | 0.145274 | 10 | 0.025077 |
| LYSOPC P22:6 | M | TX | 0.034539 | TG W - WT W | 0.129937 | 10 | 0.025077 |
| LYSOPC 20:0 | M | TX | 0.034596 | TG BD - WT BD | 0.143626 | 10 | 0.824476 |
| LYSOPC 20:0 | M | TX | 0.034596 | TG BD - TG W | 0.128463 | 10 | 0.824476 |
| LYSOPC 20:0 | M | TX | 0.034596 | TG BD - WT W | 0.143626 | 10 | 0.024952 |
| LYSOPC 20:0 | M | TX | 0.034596 | WT BD - TG W | 0.128463 | 10 | 0.824476 |
| LYSOPC 20:0 | M | TX | 0.034596 | WT BD - WT W | 0.143626 | 10 | 0.024952 |
| LYSOPC 20:0 | M | TX | 0.034596 | TG W - WT W | 0.128463 | 10 | 0.024952 |
| PS 18:0-20:0 | M | TG | 0.038108 | TG BD - WT BD | 0.056749 | 11 | 0.250422 |
| PS 18:0-20:0 | M | TG | 0.038108 | TG BD - TG W | 0.054263 | 11 | 0.709033 |
| PS 18:0-20:0 | M | TG | 0.038108 | TG BD - WT W | 0.060668 | 11 | 0.250422 |
| PS 18:0-20:0 | M | TG | 0.038108 | WT BD - TG W | 0.049844 | 11 | 0.250422 |
| PS 18:0-20:0 | M | TG | 0.038108 | WT BD - WT W | 0.056749 | 11 | 0.891343 |
| PS 18:0-20:0 | M | TG | 0.038108 | TG W - WT W | 0.054263 | 11 | 0.293293 |
| SM N20:0 | M | TX | 0.040460 | TG BD - WT BD | 0.189305 | 11 | 0.364170 |
| SM N20:0 | M | TX | 0.040460 | TG BD - TG W | 0.181011 | 11 | 0.201102 |
| SM N20:0 | M | TX | 0.040460 | TG BD - WT W | 0.202376 | 11 | 0.201102 |
| SM N20:0 | M | TX | 0.040460 | WT BD - TG W | 0.166269 | 11 | 0.527799 |
| SM N20:0 | M | TX | 0.040460 | WT BD - WT W | 0.189305 | 11 | 0.364170 |
| SM N20:0 | M | TX | 0.040460 | TG W - WT W | 0.181011 | 11 | 0.527799 |
| PI A18:0-22:3 | M | TG | 0.040585 | TG BD - WT BD | 0.022000 | 11 | 0.095164 |
| PI A18:0-22:3 | M | TG | 0.040585 | TG BD - TG W | 0.021036 | 11 | 0.189119 |
| PI A18:0-22:3 | M | TG | 0.040585 | TG BD - WT W | 0.023519 | 11 | 0.095164 |
| PI A18:0-22:3 | M | TG | 0.040585 | WT BD - TG W | 0.019323 | 11 | 0.452666 |
| PI A18:0-22:3 | M | TG | 0.040585 | WT BD - WT W | 0.022000 | 11 | 0.950571 |
| PI A18:0-22:3 | M | TG | 0.040585 | TG W - WT W | 0.021036 | 11 | 0.452666 |
| PC D18:0-20:1/P18:1-22:6 | M | TG | 0.040725 | TG BD - WT BD | 0.158270 | 11 | 0.670602 |
| PC D18:0-20:1/P18:1-22:6 | M | TG | 0.040725 | TG BD - TG W | 0.151335 | 11 | 0.558294 |
| PC D18:0-20:1/P18:1-22:6 | M | TG | 0.040725 | TG BD - WT W | 0.169197 | 11 | 0.253325 |
| PC D18:0-20:1/P18:1-22:6 | M | TG | 0.040725 | WT BD - TG W | 0.139010 | 11 | 0.319955 |
| PC D18:0-20:1/P18:1-22:6 | M | TG | 0.040725 | WT BD - WT W | 0.158270 | 11 | 0.280037 |
| PC D18:0-20:1/P18:1-22:6 | M | TG | 0.040725 | TG W - WT W | 0.151335 | 11 | 0.090326 |
| LYSOPC 22:6 | M | TG:TX | 0.040806 | TG BD - WT BD | 0.085862 | 10 | 0.424645 |
| LYSOPC 22:6 | M | TG:TX | 0.040806 | TG BD - TG W | 0.076798 | 10 | 0.903130 |
| LYSOPC 22:6 | M | TG:TX | 0.040806 | TG BD - WT W | 0.085862 | 10 | 0.133492 |
| LYSOPC 22:6 | M | TG:TX | 0.040806 | WT BD - TG W | 0.076798 | 10 | 0.424645 |
| LYSOPC 22:6 | M | TG:TX | 0.040806 | WT BD - WT W | 0.085862 | 10 | 0.075029 |
| LYSOPC 22:6 | M | TG:TX | 0.040806 | TG W - WT W | 0.076798 | 10 | 0.107376 |
| PS 18:1-22:6 | M | TX | 0.041060 | TG BD - WT BD | 0.192906 | 11 | 0.959385 |
| PS 18:1-22:6 | M | TX | 0.041060 | TG BD - TG W | 0.184454 | 11 | 0.272689 |
| PS 18:1-22:6 | M | TX | 0.041060 | TG BD - WT W | 0.206226 | 11 | 0.272689 |
| PS 18:1-22:6 | M | TX | 0.041060 | WT BD - TG W | 0.169432 | 11 | 0.272689 |
| PS 18:1-22:6 | M | TX | 0.041060 | WT BD - WT W | 0.192906 | 11 | 0.272689 |
| PS 18:1-22:6 | M | TX | 0.041060 | TG W - WT W | 0.184454 | 11 | 0.695597 |
| LYSOPE P16:0 | M | TG:TX | 0.045161 | TG BD - WT BD | 0.115592 | 10 | 0.466185 |
| LYSOPE P16:0 | M | TG:TX | 0.045161 | TG BD - TG W | 0.103388 | 10 | 0.491772 |
| LYSOPE P16:0 | M | TG:TX | 0.045161 | TG BD - WT W | 0.115592 | 10 | 0.466185 |
| LYSOPE P16:0 | M | TG:TX | 0.045161 | WT BD - TG W | 0.103388 | 10 | 0.615706 |
| LYSOPE P16:0 | M | TG:TX | 0.045161 | WT BD - WT W | 0.115592 | 10 | 0.201219 |
| LYSOPE P16:0 | M | TG:TX | 0.045161 | TG W - WT W | 0.103388 | 10 | 0.201219 |
| PC D18:0-20:2/P18:2-22:6 | M | TX | 0.047000 | TG BD - WT BD | 0.180405 | 11 | 0.234358 |
| PC D18:0-20:2/P18:2-22:6 | M | TX | 0.047000 | TG BD - TG W | 0.172500 | 11 | 0.234358 |
| PC D18:0-20:2/P18:2-22:6 | M | TX | 0.047000 | TG BD - WT W | 0.192861 | 11 | 0.110410 |
| PC D18:0-20:2/P18:2-22:6 | M | TX | 0.047000 | WT BD - TG W | 0.158452 | 11 | 0.887694 |
| PC D18:0-20:2/P18:2-22:6 | M | TX | 0.047000 | WT BD - WT W | 0.180405 | 11 | 0.234358 |
| PC D18:0-20:2/P18:2-22:6 | M | TX | 0.047000 | TG W - WT W | 0.172500 | 11 | 0.234358 |
| PS 18:0-22:6...139 | M | TX | 0.047179 | TG BD - WT BD | 0.223230 | 11 | 0.730131 |
| PS 18:0-22:6...139 | M | TX | 0.047179 | TG BD - TG W | 0.213448 | 11 | 0.438238 |
| PS 18:0-22:6...139 | M | TX | 0.047179 | TG BD - WT W | 0.238642 | 11 | 0.272536 |
| PS 18:0-22:6...139 | M | TX | 0.047179 | WT BD - TG W | 0.196065 | 11 | 0.272536 |
| PS 18:0-22:6...139 | M | TX | 0.047179 | WT BD - WT W | 0.223230 | 11 | 0.272536 |
| PS 18:0-22:6...139 | M | TX | 0.047179 | TG W - WT W | 0.213448 | 11 | 0.602235 |
| PS 18:0-22:6...158 | M | TX | 0.047179 | TG BD - WT BD | 0.223230 | 11 | 0.730131 |
| PS 18:0-22:6...158 | M | TX | 0.047179 | TG BD - TG W | 0.213448 | 11 | 0.438238 |
| PS 18:0-22:6...158 | M | TX | 0.047179 | TG BD - WT W | 0.238642 | 11 | 0.272536 |
| PS 18:0-22:6...158 | M | TX | 0.047179 | WT BD - TG W | 0.196065 | 11 | 0.272536 |
| PS 18:0-22:6...158 | M | TX | 0.047179 | WT BD - WT W | 0.223230 | 11 | 0.272536 |
| PS 18:0-22:6...158 | M | TX | 0.047179 | TG W - WT W | 0.213448 | 11 | 0.602235 |
| LYSOPE 16:0 | M | TG:TX | 0.048299 | TG BD - WT BD | 0.079026 | 10 | 0.262780 |
| LYSOPE 16:0 | M | TG:TX | 0.048299 | TG BD - TG W | 0.070683 | 10 | 0.262780 |
| LYSOPE 16:0 | M | TG:TX | 0.048299 | TG BD - WT W | 0.079026 | 10 | 0.632918 |
| LYSOPE 16:0 | M | TG:TX | 0.048299 | WT BD - TG W | 0.070683 | 10 | 0.900213 |
| LYSOPE 16:0 | M | TG:TX | 0.048299 | WT BD - WT W | 0.079026 | 10 | 0.365703 |
| LYSOPE 16:0 | M | TG:TX | 0.048299 | TG W - WT W | 0.070683 | 10 | 0.365703 |
| PI 18:0-22:6 | M | TG | 0.048390 | TG BD - WT BD | 0.107479 | 11 | 0.262899 |
| PI 18:0-22:6 | M | TG | 0.048390 | TG BD - TG W | 0.102770 | 11 | 0.548242 |
| PI 18:0-22:6 | M | TG | 0.048390 | TG BD - WT W | 0.114900 | 11 | 0.262899 |
| PI 18:0-22:6 | M | TG | 0.048390 | WT BD - TG W | 0.094400 | 11 | 0.332859 |
| PI 18:0-22:6 | M | TG | 0.048390 | WT BD - WT W | 0.107479 | 11 | 0.836794 |
| PI 18:0-22:6 | M | TG | 0.048390 | TG W - WT W | 0.102770 | 11 | 0.332859 |
| PI 18:2-20:4/16:0-22:6 | M | TG | 0.048909 | TG BD - WT BD | 0.070808 | 11 | 0.266370 |
| PI 18:2-20:4/16:0-22:6 | M | TG | 0.048909 | TG BD - TG W | 0.067705 | 11 | 0.476756 |
| PI 18:2-20:4/16:0-22:6 | M | TG | 0.048909 | TG BD - WT W | 0.075697 | 11 | 0.278940 |
| PI 18:2-20:4/16:0-22:6 | M | TG | 0.048909 | WT BD - TG W | 0.062191 | 11 | 0.278940 |
| PI 18:2-20:4/16:0-22:6 | M | TG | 0.048909 | WT BD - WT W | 0.070808 | 11 | 0.582821 |
| PI 18:2-20:4/16:0-22:6 | M | TG | 0.048909 | TG W - WT W | 0.067705 | 11 | 0.476756 |
| LYSOPC 20:4 | M | TG:TX | 0.049207 | TG BD - WT BD | 0.127010 | 10 | 0.328583 |
| LYSOPC 20:4 | M | TG:TX | 0.049207 | TG BD - TG W | 0.113601 | 10 | 0.287838 |
| LYSOPC 20:4 | M | TG:TX | 0.049207 | TG BD - WT W | 0.127010 | 10 | 0.963501 |
| LYSOPC 20:4 | M | TG:TX | 0.049207 | WT BD - TG W | 0.113601 | 10 | 0.861471 |
| LYSOPC 20:4 | M | TG:TX | 0.049207 | WT BD - WT W | 0.127010 | 10 | 0.328583 |
| LYSOPC 20:4 | M | TG:TX | 0.049207 | TG W - WT W | 0.113601 | 10 | 0.287838 |

**Supplemental Table 5:**
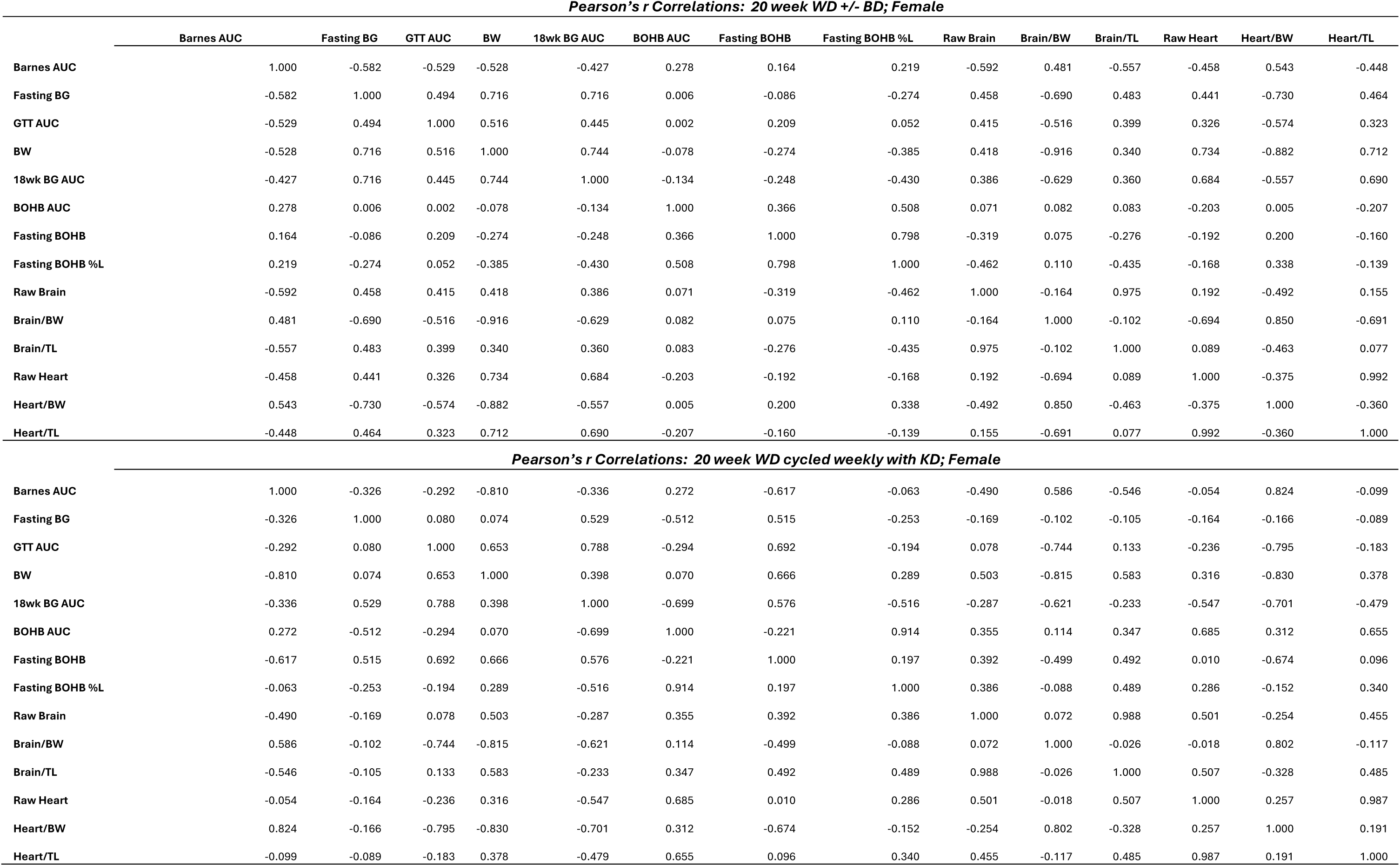
Pearson’s correlation of metabolic variables with cognitive outcome in Western Diet-fed female mice. Pearson’s correlation coefficient, r, for various metabolic variables following a 20-week Western diet (WD) with or without 10% BD dissolved in drinking water, or WD cycling weekly with ketogenic diet (KD). Positive values indicate positive correlation and negative values indicate negative correlation. *n*=10-21 females in correlation calculations. BD = 1,3-butanediol, AUC = area under curve, GTT = glucose tolerance test, BG = blood glucose, BW = body weight, TL = tibia length, Glc = glucose, βOHB = total beta-hydroxybutyrate (both D and L enantiomers), AcAc = acetoacetate.

**Pearson's r Correlations: 20 week WD +/- BD; Female**
|  | Barnes AUC | Fasting BG | GTT AUC | BW | 18wk BG AUC | BOHB AUC | Fasting BOHB | Fasting BOHB %L | Raw Brain | Brain/BW | Brain/TL | Raw Heart | Heart/BW | Heart/TL |
| --- | --- | --- | --- | --- | --- | --- | --- | --- | --- | --- | --- | --- | --- | --- |
| Barnes AUC | 1.000 | -0.582 | -0.529 | -0.528 | -0.427 | 0.278 | 0.164 | 0.219 | -0.592 | 0.481 | -0.557 | -0.458 | 0.543 | -0.448 |
| Fasting BG | -0.582 | 1.000 | 0.494 | 0.716 | 0.716 | 0.006 | -0.086 | -0.274 | 0.458 | -0.690 | 0.483 | 0.441 | -0.730 | 0.464 |
| GTT AUC | -0.529 | 0.494 | 1.000 | 0.516 | 0.445 | 0.002 | 0.209 | 0.052 | 0.415 | -0.516 | 0.399 | 0.326 | -0.574 | 0.323 |
| BW | -0.528 | 0.716 | 0.516 | 1.000 | 0.744 | -0.078 | -0.274 | -0.385 | 0.418 | -0.916 | 0.340 | 0.734 | -0.882 | 0.712 |
| 18wk BG AUC | -0.427 | 0.716 | 0.445 | 0.744 | 1.000 | -0.134 | -0.248 | -0.430 | 0.386 | -0.629 | 0.360 | 0.684 | -0.557 | 0.690 |
| BOHB AUC | 0.278 | 0.006 | 0.002 | -0.078 | -0.134 | 1.000 | 0.366 | 0.508 | 0.071 | 0.082 | 0.083 | -0.203 | 0.005 | -0.207 |
| Fasting BOHB | 0.164 | -0.086 | 0.209 | -0.274 | -0.248 | 0.366 | 1.000 | 0.798 | -0.319 | 0.075 | -0.276 | -0.192 | 0.200 | -0.160 |
| Fasting BOHB %L | 0.219 | -0.274 | 0.052 | -0.385 | -0.430 | 0.508 | 0.798 | 1.000 | -0.462 | 0.110 | -0.435 | -0.168 | 0.338 | -0.139 |
| Raw Brain | -0.592 | 0.458 | 0.415 | 0.418 | 0.386 | 0.071 | -0.319 | -0.462 | 1.000 | -0.164 | 0.975 | 0.192 | -0.492 | 0.155 |
| Brain/BW | 0.481 | -0.690 | -0.516 | -0.916 | -0.629 | 0.082 | 0.075 | 0.110 | -0.164 | 1.000 | -0.102 | -0.694 | 0.850 | -0.691 |
| Brain/TL | -0.557 | 0.483 | 0.399 | 0.340 | 0.360 | 0.083 | -0.276 | -0.435 | 0.975 | -0.102 | 1.000 | 0.089 | -0.463 | 0.077 |
| Raw Heart | -0.458 | 0.441 | 0.326 | 0.734 | 0.684 | -0.203 | -0.192 | -0.168 | 0.192 | -0.694 | 0.089 | 1.000 | -0.375 | 0.992 |
| Heart/BW | 0.543 | -0.730 | -0.574 | -0.882 | -0.557 | 0.005 | 0.200 | 0.338 | -0.492 | 0.850 | -0.463 | -0.375 | 1.000 | -0.360 |
| Heart/TL | -0.448 | 0.464 | 0.323 | 0.712 | 0.690 | -0.207 | -0.160 | -0.139 | 0.155 | -0.691 | 0.077 | 0.992 | -0.360 | 1.000 |

**Supplemental Table 5: Pearson's correlation of metabolic variables with cognitive outcome in Western Diet-fed female mice.**
|  |  |  |  |  |  |  |  |  |  |  |  |  |  |  |
| --- | --- | --- | --- | --- | --- | --- | --- | --- | --- | --- | --- | --- | --- | --- |
| Barnes AUC | 1.000 | -0.326 | -0.292 | -0.810 | -0.336 | 0.272 | -0.617 | -0.063 | -0.490 | 0.586 | -0.546 | -0.054 | 0.824 | -0.099 |
| Fasting BG | -0.326 | 1.000 | 0.080 | 0.074 | 0.529 | -0.512 | 0.515 | -0.253 | -0.169 | -0.102 | -0.105 | -0.164 | -0.166 | -0.089 |
| GTT AUC | -0.292 | 0.080 | 1.000 | 0.653 | 0.788 | -0.294 | 0.692 | -0.194 | 0.078 | -0.744 | 0.133 | -0.236 | -0.795 | -0.183 |
| BW | -0.810 | 0.074 | 0.653 | 1.000 | 0.398 | 0.070 | 0.666 | 0.289 | 0.503 | -0.815 | 0.583 | 0.316 | -0.830 | 0.378 |
| 18wk BG AUC | -0.336 | 0.529 | 0.788 | 0.398 | 1.000 | -0.699 | 0.576 | -0.516 | -0.287 | -0.621 | -0.233 | -0.547 | -0.701 | -0.479 |
| BOHB AUC | 0.272 | -0.512 | -0.294 | 0.070 | -0.699 | 1.000 | -0.221 | 0.914 | 0.355 | 0.114 | 0.347 | 0.685 | 0.312 | 0.655 |
| Fasting BOHB | -0.617 | 0.515 | 0.692 | 0.666 | 0.576 | -0.221 | 1.000 | 0.197 | 0.392 | -0.499 | 0.492 | 0.010 | -0.674 | 0.096 |
| Fasting BOHB %L | -0.063 | -0.253 | -0.194 | 0.289 | -0.516 | 0.914 | 0.197 | 1.000 | 0.386 | -0.088 | 0.489 | 0.286 | -0.152 | 0.340 |
| Raw Brain | -0.490 | -0.169 | 0.078 | 0.503 | -0.287 | 0.355 | 0.392 | 0.386 | 1.000 | 0.072 | 0.988 | 0.501 | -0.254 | 0.455 |
| Brain/BW | 0.586 | -0.102 | -0.744 | -0.815 | -0.621 | 0.114 | -0.499 | -0.088 | 0.072 | 1.000 | -0.026 | -0.018 | 0.802 | -0.117 |
| Brain/TL | -0.546 | -0.105 | 0.133 | 0.583 | -0.233 | 0.347 | 0.492 | 0.489 | 0.988 | -0.026 | 1.000 | 0.507 | -0.328 | 0.485 |
| Raw Heart | -0.054 | -0.164 | -0.236 | 0.316 | -0.547 | 0.685 | 0.010 | 0.286 | 0.501 | -0.018 | 0.507 | 1.000 | 0.257 | 0.987 |
| Heart/BW | 0.824 | -0.166 | -0.795 | -0.830 | -0.701 | 0.312 | -0.674 | -0.152 | -0.254 | 0.802 | -0.328 | 0.257 | 1.000 | 0.191 |
| Heart/TL | -0.099 | -0.089 | -0.183 | 0.378 | -0.479 | 0.655 | 0.096 | 0.340 | 0.455 | -0.117 | 0.485 | 0.987 | 0.191 | 1.000 |

**Supplemental Table 6:** Pearson’s correlation of metabolic variables with cognitive outcome in Western Diet-fed male mice. Pearson’s correlation coefficient, r, for various metabolic variables following a 20-week Western diet (WD) with or without 10% BD dissolved in drinking water, or WD cycling weekly with ketogenic diet (KD). Positive values indicate positive correlation and negative values indicate negative correlation. *n*=10-26 males in correlation calculations. BD = 1,3-butanediol, AUC = area under curve, GTT = glucose tolerance test, BG = blood glucose, BW = body weight, TL = tibia length, Glc = glucose, βOHB = total beta-hydroxybutyrate (both D and L enantiomers), AcAc = acetoacetate.

|  | Barnes AUC | Fasting BG | GTT AUC | BW | 18wk BG AUC | BOHB AUC | Fasting BOHB | Fasting BOHB %L | Raw Brain | Brain/BW | Brain/TL | Raw Heart | Heart/BW | Heart/TL |
| --- | --- | --- | --- | --- | --- | --- | --- | --- | --- | --- | --- | --- | --- | --- |
| Barnes AUC | 1.000 | -0.065 | -0.156 | -0.318 | -0.394 | 0.396 | 0.172 | 0.295 | -0.450 | 0.138 | -0.304 | -0.003 | 0.299 | 0.092 |
| Fasting BG | -0.065 | 1.000 | 0.552 | 0.350 | 0.647 | -0.113 | 0.074 | -0.048 | -0.291 | -0.484 | -0.300 | -0.082 | -0.439 | -0.076 |
| GTT AUC | -0.156 | 0.552 | 1.000 | 0.116 | 0.245 | -0.372 | -0.220 | -0.294 | -0.396 | -0.403 | -0.489 | -0.539 | -0.609 | -0.588 |
| BW | -0.318 | 0.350 | 0.116 | 1.000 | 0.597 | 0.016 | 0.228 | -0.333 | 0.420 | -0.903 | 0.135 | 0.481 | -0.735 | 0.333 |
| 18wk BG AUC | -0.394 | 0.647 | 0.245 | 0.597 | 1.000 | -0.079 | 0.182 | -0.210 | 0.404 | -0.425 | 0.193 | 0.313 | -0.368 | 0.194 |
| BOHB AUC | 0.396 | -0.113 | -0.372 | 0.016 | -0.079 | 1.000 | -0.067 | 0.466 | 0.120 | 0.076 | 0.123 | 0.180 | 0.176 | 0.185 |
| Fasting BOHB | 0.172 | 0.074 | -0.220 | 0.228 | 0.182 | -0.067 | 1.000 | -0.244 | 0.029 | -0.195 | -0.112 | 0.351 | -0.014 | 0.274 |
| Fasting BOHB %L | 0.295 | -0.048 | -0.294 | -0.333 | -0.210 | 0.466 | -0.244 | 1.000 | 0.105 | 0.384 | 0.237 | 0.417 | 0.602 | 0.485 |
| Raw Brain | -0.450 | -0.291 | -0.396 | 0.420 | 0.404 | 0.120 | 0.029 | 0.105 | 1.000 | -0.029 | 0.894 | 0.536 | -0.019 | 0.451 |
| Brain/BW | 0.138 | -0.484 | -0.403 | -0.903 | -0.425 | 0.076 | -0.195 | 0.384 | -0.029 | 1.000 | 0.244 | -0.240 | 0.854 | -0.110 |
| Brain/TL | -0.304 | -0.300 | -0.489 | 0.135 | 0.193 | 0.123 | -0.112 | 0.237 | 0.894 | 0.244 | 1.000 | 0.475 | 0.247 | 0.500 |
| Raw Heart | -0.003 | -0.082 | -0.539 | 0.481 | 0.313 | 0.180 | 0.351 | 0.417 | 0.536 | -0.240 | 0.475 | 1.000 | 0.217 | 0.970 |
| Heart/BW | 0.299 | -0.439 | -0.609 | -0.735 | -0.368 | 0.176 | -0.014 | 0.602 | -0.019 | 0.854 | 0.247 | 0.217 | 1.000 | 0.355 |
| Heart/TL | 0.092 | -0.076 | -0.588 | 0.333 | 0.194 | 0.185 | 0.274 | 0.485 | 0.451 | -0.110 | 0.500 | 0.970 | 0.355 | 1.000 |

**Supplemental Table 6: Pearson's correlation of metabolic variables with cognitive outcome in Western Diet-fed male mice.**
|  | Barnes AUC | Fasting BG | GTT AUC | BW | 18wk BG AUC | TKB AUC | Fasting BOHB | Fasting BOHB %L | Raw Brain | Brain/BW | Brain/TL | Raw Heart | Heart/BW | Heart/TL |
| --- | --- | --- | --- | --- | --- | --- | --- | --- | --- | --- | --- | --- | --- | --- |
| Barnes AUC | 1.000 | 0.068 | -0.760 | -0.722 | -0.750 | -0.010 | 0.428 | -0.267 | -0.664 | 0.442 | -0.621 | 0.592 | 0.761 | 0.630 |
| Fasting BG | 0.068 | 1.000 | 0.289 | 0.082 | 0.221 | 0.572 | -0.243 | -0.709 | 0.171 | 0.068 | 0.249 | 0.093 | 0.065 | 0.146 |
| GTT AUC | -0.760 | 0.289 | 1.000 | 0.864 | 0.808 | 0.334 | -0.379 | -0.018 | 0.588 | -0.667 | 0.550 | -0.557 | -0.832 | -0.592 |
| BW | -0.722 | 0.082 | 0.864 | 1.000 | 0.828 | 0.384 | -0.304 | -0.119 | 0.592 | -0.845 | 0.479 | -0.415 | -0.868 | -0.501 |
| 18wk BG AUC | -0.750 | 0.221 | 0.808 | 0.828 | 1.000 | 0.143 | -0.235 | -0.063 | 0.482 | -0.720 | 0.448 | -0.632 | -0.888 | -0.662 |
| TKB AUC | -0.010 | 0.572 | 0.334 | 0.384 | 0.143 | 1.000 | -0.371 | -0.491 | 0.624 | -0.054 | 0.597 | 0.309 | -0.040 | 0.277 |
| Fasting BOHB | 0.428 | -0.243 | -0.379 | -0.304 | -0.235 | -0.371 | 1.000 | 0.097 | -0.668 | -0.110 | -0.721 | -0.038 | 0.125 | -0.064 |
| Fasting BOHB %L | -0.267 | -0.709 | -0.018 | -0.119 | -0.063 | -0.491 | 0.097 | 1.000 | 0.047 | 0.148 | 0.067 | -0.744 | -0.271 | -0.717 |
| Raw Brain | -0.664 | 0.171 | 0.588 | 0.592 | 0.482 | 0.624 | -0.668 | 0.047 | 1.000 | -0.083 | 0.981 | -0.031 | -0.373 | -0.065 |
| Brain/BW | 0.442 | 0.068 | -0.667 | -0.845 | -0.720 | -0.054 | -0.110 | 0.148 | -0.083 | 1.000 | 0.045 | 0.487 | 0.840 | 0.574 |
| Brain/TL | -0.621 | 0.249 | 0.550 | 0.479 | 0.448 | 0.597 | -0.721 | 0.067 | 0.981 | 0.045 | 1.000 | -0.038 | -0.299 | -0.045 |
| Raw Heart | 0.592 | 0.093 | -0.557 | -0.415 | -0.632 | 0.309 | -0.038 | -0.744 | -0.031 | 0.487 | -0.038 | 1.000 | 0.796 | 0.991 |
| Heart/BW | 0.761 | 0.065 | -0.832 | -0.868 | -0.888 | -0.040 | 0.125 | -0.271 | -0.373 | 0.840 | -0.299 | 0.796 | 1.000 | 0.850 |
| Heart/TL | 0.630 | 0.146 | -0.592 | -0.501 | -0.662 | 0.277 | -0.064 | -0.717 | -0.065 | 0.574 | -0.045 | 0.991 | 0.850 | 1.000 |

**Supplemental Table 7:**
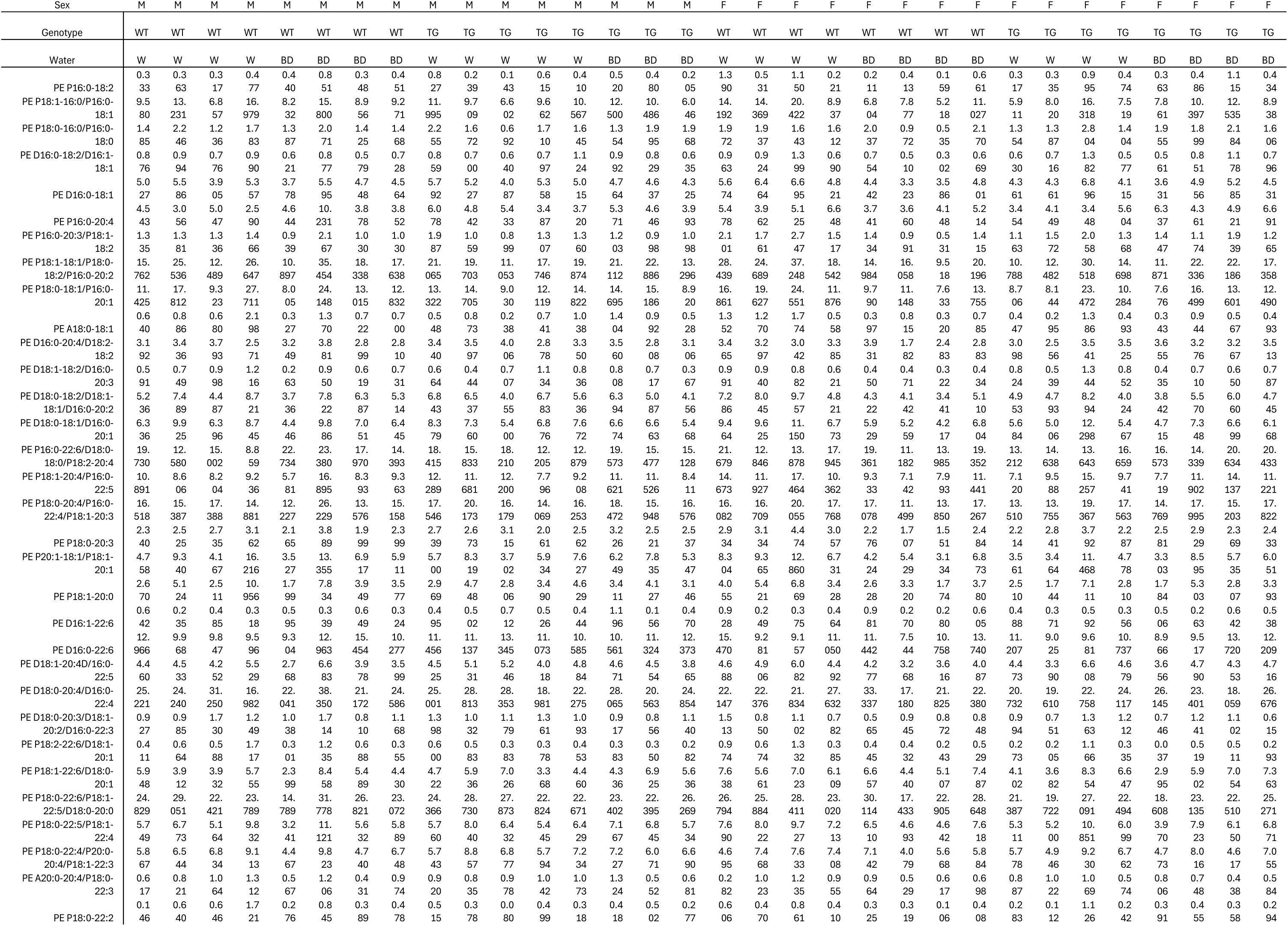
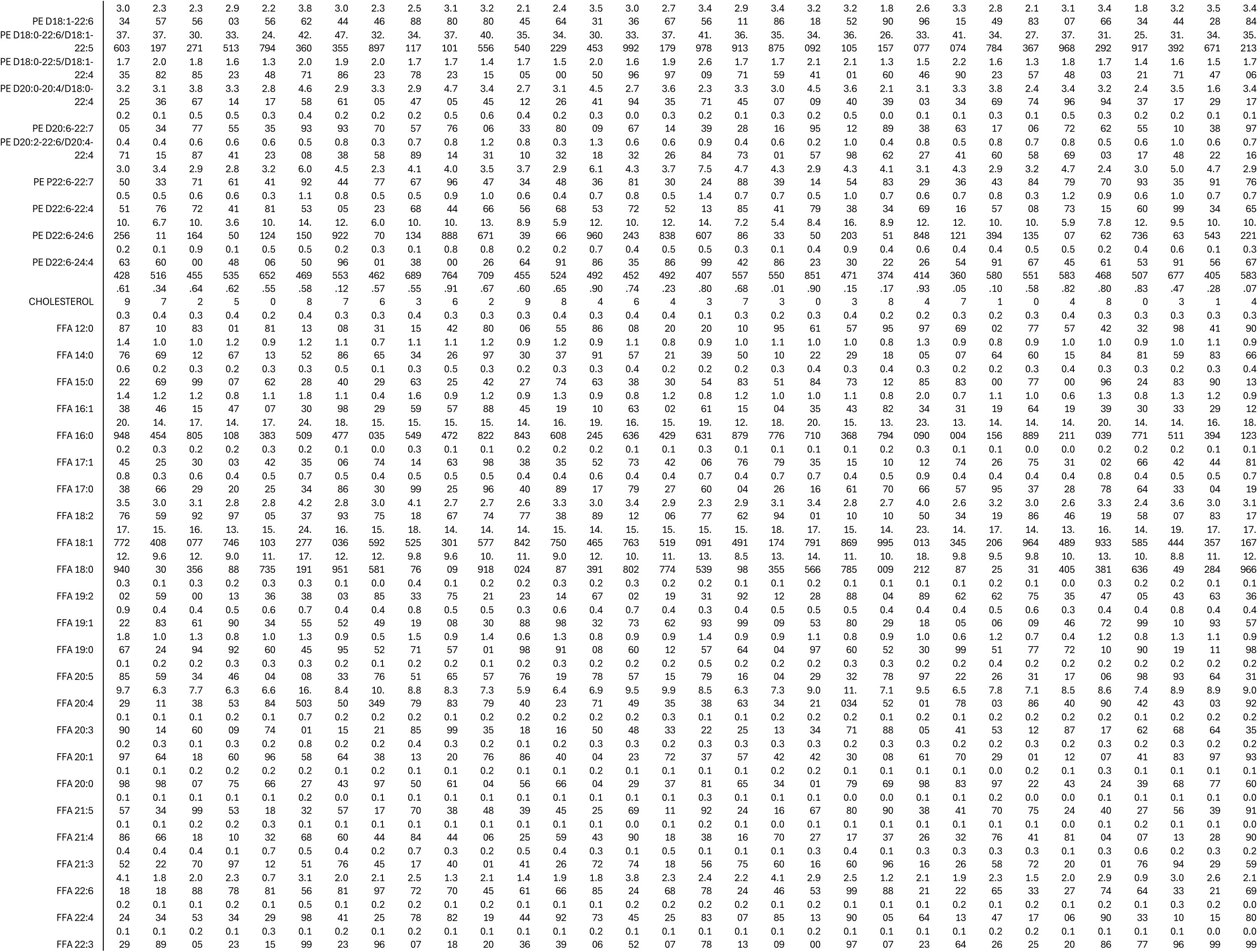
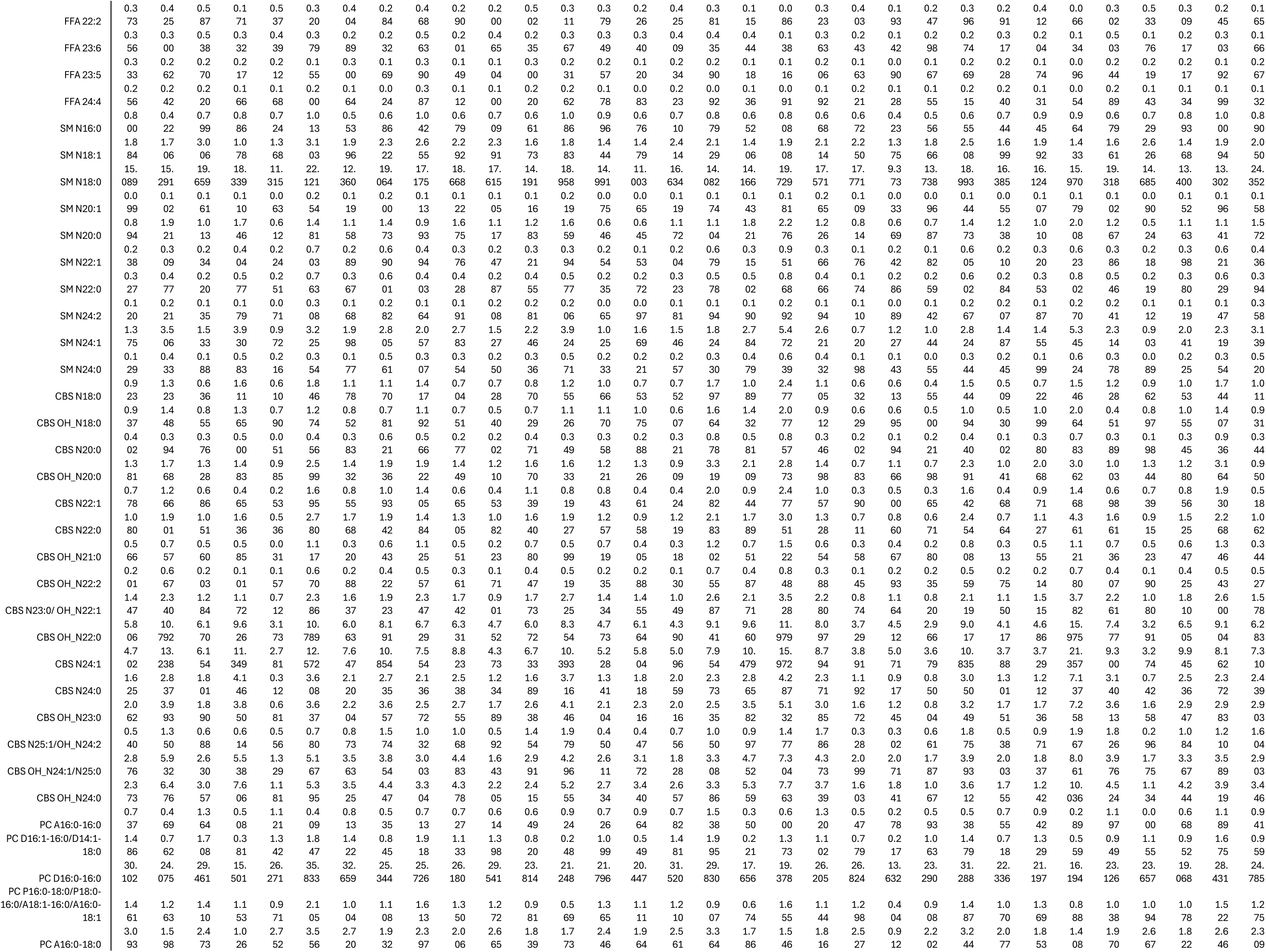
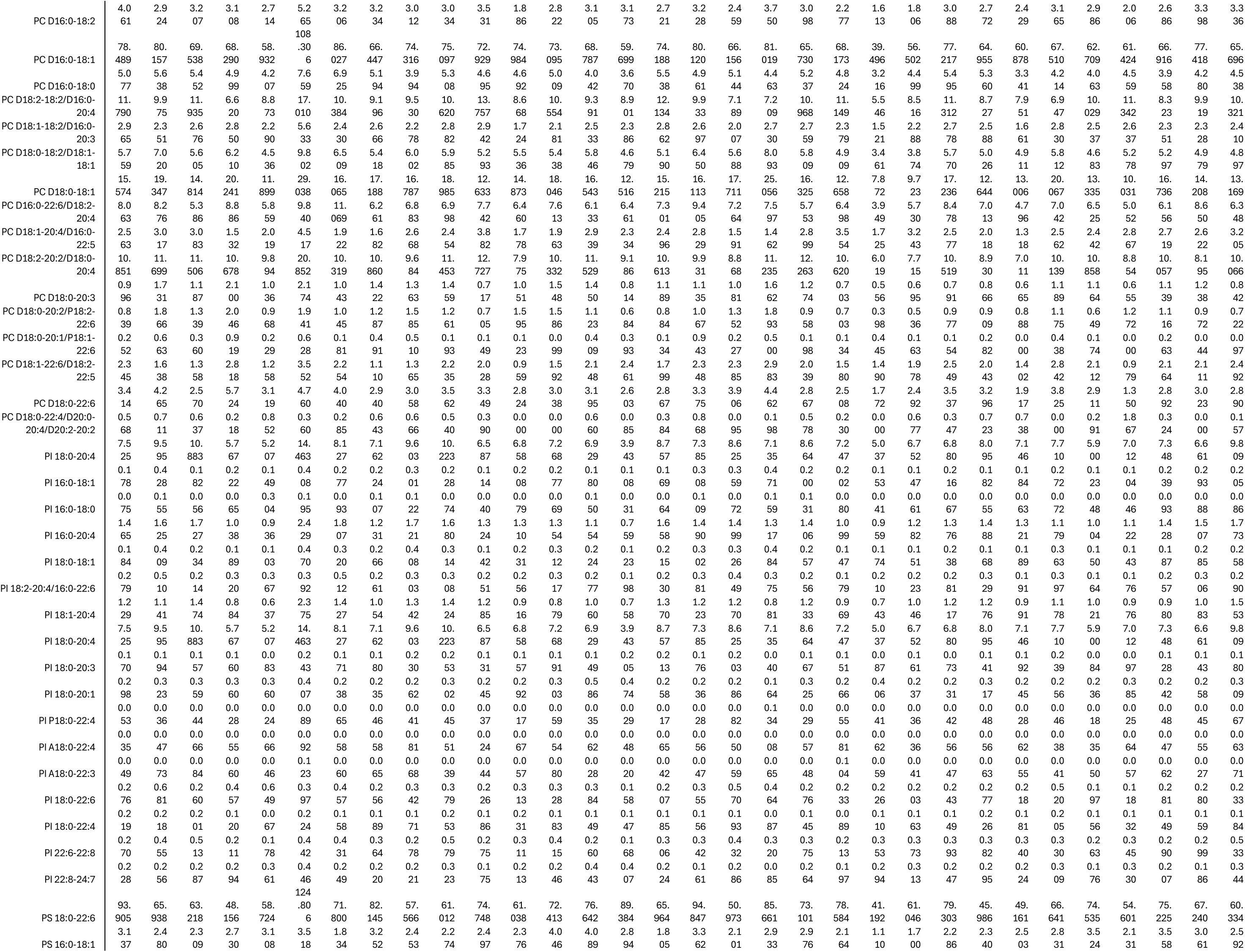
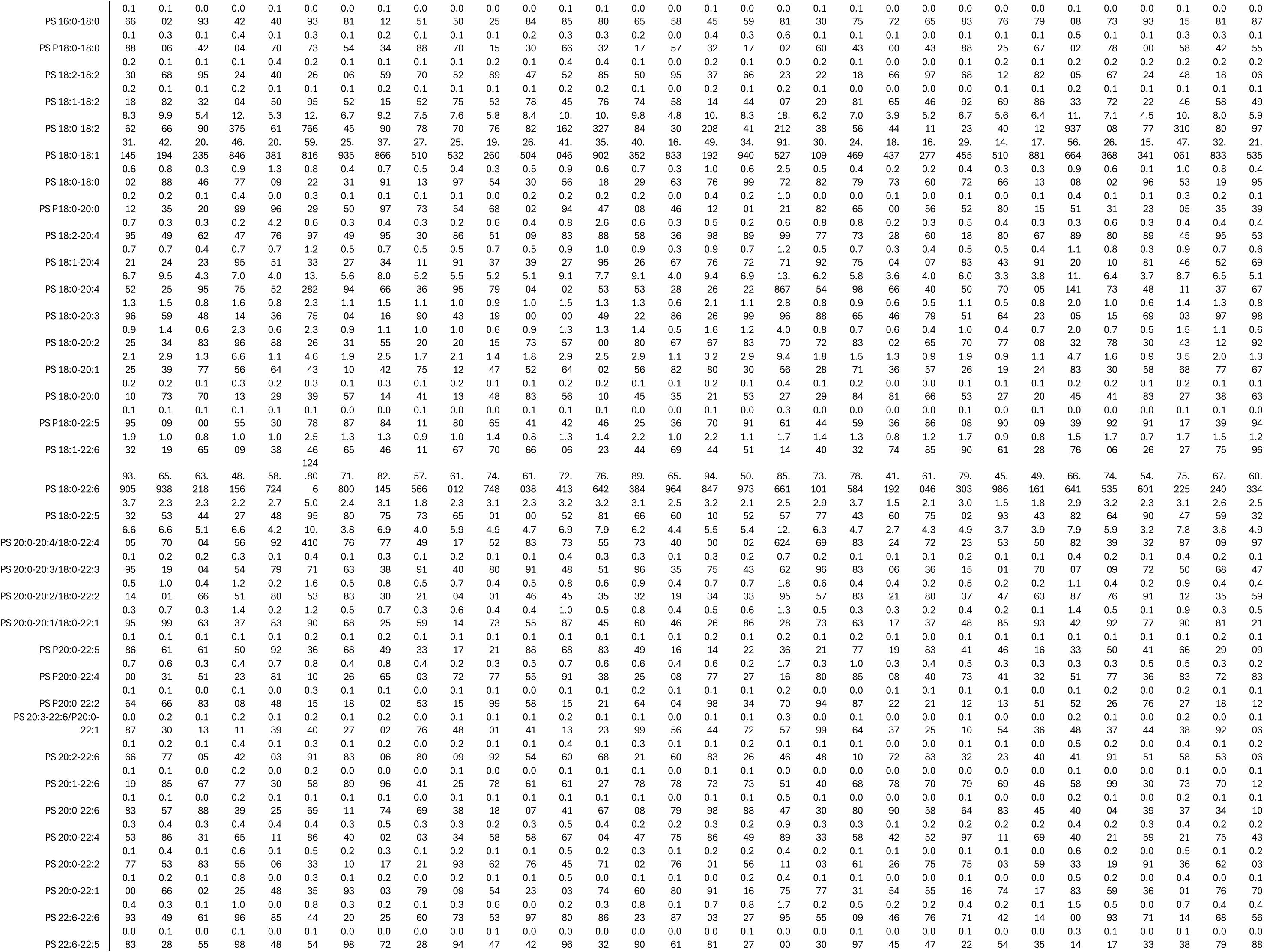
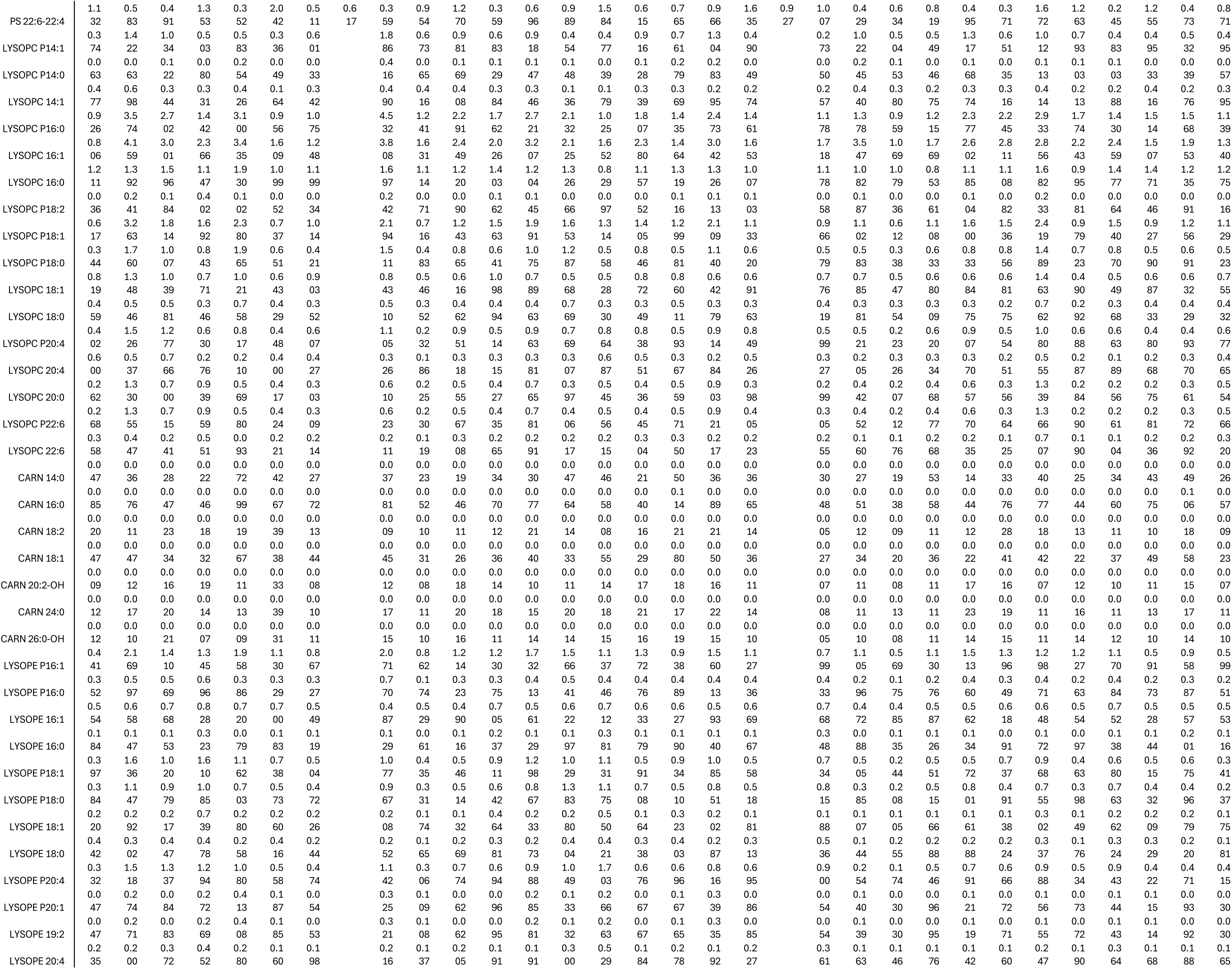
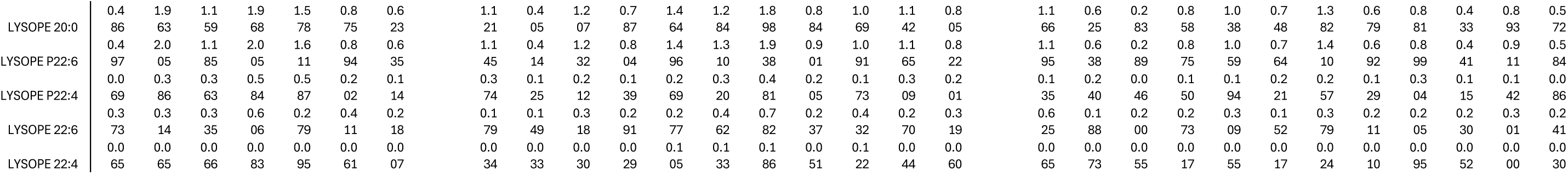
Absolute concentrations of lipid species. Quantification of lipid species (nmol / g protein) from freeze-clamped, forebrain regions of male and female, tau-transgenic and non-transgenic mice consuming 10% BD or normal drinking water for 20 weeks. M = male, F = female, WT = non-transgenic mice, TG = tau-transgenic mice, W = water, BD = 1,3-butanediol, PC = phosphatidylcholine, PE = phosphatidylethanolamine, PI = phosphatidylinositol, PS = phosphatidylserine, SM = sphingomyelin, CBS = cerebroside, FFA = free fatty-acid, LPC = lysoPC, LPE = lysoPE, Car = acylcarnitine

